# Being Born versus Being Born Early: Disentangling Birth from Prematurity in Perinatal Brain Development

**DOI:** 10.64898/2026.11.21.768316

**Authors:** Ying-Qiu Zheng, Matteo Bastiani, Frederik J. Lange, Vyacheslav Karolis, Alex Fung, Hossein Rafipoor, Michiel Cottaar, Haoming Liu, Jianwei Cui, Zeju Li, Rebeccah Slater, Saad Jbabdi

## Abstract

Studies of preterm birth report both delayed and accelerated brain maturation, depending on whether preterm infants are compared with term newborns or with fetuses. We propose that these mixed observations arise because such comparisons conflate two distinct influences, the effect of birth on term-aged infants and the independent effects of being born prematurely. Using MRI data from 330 term and 118 preterm neonates in the developing Human Connectome Project (dHCP), we developed a simple piecewise linear model that estimates prenatal and postnatal rates of development separately for term and preterm infants across volumetric, microstructural and connectivity measures. In term infants, birth accelerated volumetric growth, with total brain volume growing 44% faster post-birth, with regionally specific alterations in microstructural trajectories. Deep white matter continued to mature, whereas superficial and cerebellar white matter reversed direction. Changes in cortical connectivity at birth followed the sensorimotor to association axis, shifting towards specialisation in sensory and insular cortex and towards more diverse connectivity in association cortex. In contrast, the effect of birth on preterm infants was much weaker, where development largely continued at prenatal rates. These results show that the impact of birth depends on brain region and maturational state at birth. Separating birth from prematurity offers a framework for interpreting mixed findings in the perinatal literature.

## INTRODUCTION

The perinatal period is a critical transition in human brain development. It is marked by rapid microstructural and morphological changes that establish the neuroanatomical foundation for lifelong cognitive abilities and behaviours (Knickmeyer et al. 2008; Gao et al. 2008; Dubois et al. 2014; Lyall et al. 2014). Disruptions during this vulnerable window are associated with elevated risks for neuropsychiatric disorders that may not manifest until years or decades later (Nosarti et al. 2012; Johnson and Marlow 2011).

Preterm birth (before 37 weeks of gestation) is a major disruption to perinatal neurodevelopment, particularly when it occurs before 32 weeks (Volpe 2018). Typically, the transition from intrauterine to extrauterine life occurs at term when the infant is physiologically mature enough to breathe, feed, and regulate body temperature independently. Preterm birth exposes the brain to extrauterine conditions before it has completed key stages of maturation. Histological studies show that preterm birth disrupts neurogenesis, which continues into the third trimester (Malik et al. 2013; Tibrewal et al. 2017). This period is also marked by rapid synaptogenesis, dendritic arborisation, and the onset of myelination (Tau and Peterson 2009; Volpe 2018). Neuroimaging studies have complemented these microscopic findings, revealing widespread alterations in brain development following preterm birth. These include disrupted cerebral metabolism (Vinall et al. 2013; Kwon et al. 2014; Basu et al. 2019), reduced cortical development (Ajayi-Obe et al. 2000; Lefèvre et al. 2015; Orasanu et al. 2016), altered functional connectivity (Bouyssi-Kobar et al. 2019; De Asis-Cruz et al. 2020), delayed white matter maturation (Grotheer et al. 2023), and impaired overall brain growth trajectories (Padilla et al. 2014; Bouyssi-Kobar et al. 2016).

However, neuroimaging studies have produced mixed findings on how preterm birth affects maturation. In the neonatal period, preterm infants at term equivalent age show weaker and less complex resting state connectivity than term-born newborns, including altered thalamocortical connections (Smyser et al. 2010). Yet when very preterm infants are scanned before term and compared with gestational age matched fetuses, several interhemispheric connections are stronger ex utero, particularly in occipital, superior temporal and medial temporal regions (De Asis-Cruz et al. 2020). The visual system shows a similar pattern. Very preterm infants have larger occipital volumes (Padilla et al. 2014) and more mature visual behaviour at term equivalent age (Ricci et al. 2008), and their visual evoked responses mature faster with longer extrauterine exposure (Schwindt et al. 2018). Preterm infants therefore appear advanced in some measures and delayed in others, even when compared with the same reference group of term born infants at term equivalent age. Advanced maturation has been attributed to early extrauterine sensory experience (De Asis-Cruz et al. 2020; Schwindt et al. 2018; Schmidt Mellado et al. 2022), whereas delays are typically attributed to brain injury and adverse exposures in the neonatal period (Volpe 2018).

These contradictory findings may reflect a methodological limitation. Comparisons between preterm and term infants confound two separate developmental influences, being born and being born prematurely. Grotheer et al. (Grotheer et al. 2023) examined the effect of birth directly, reporting uniformly slower white matter growth ex utero than in utero in the developing Human Connectome Project (dHCP) cohort. However, they examined a single imaging measure (myelination) and assumed that birth has the same effect regardless of gestational age at birth. Whether the effect of birth differs between term and preterm infants and across structural domains therefore remains unknown.

Animal studies suggest three possible hypotheses about how birth itself affects neurodevelopment (Bourgeois et al. 1989; Toda et al. 2013). First, the “acceleration hypothesis” proposes that the dramatic increase in incoming sensory information at birth, such as the increase in visual input, provokes an increased rate of growth ex utero compared to in utero. Second, the “protection hypothesis” suggests that the protective environment of the womb, with its consistent supply of nutrients, oxygen and warmth, could promote more rapid growth in utero compared to ex utero. Third, the null hypothesis suggests that birth does not impact growth trajectories (Figure 1A).

**Figure 1.**
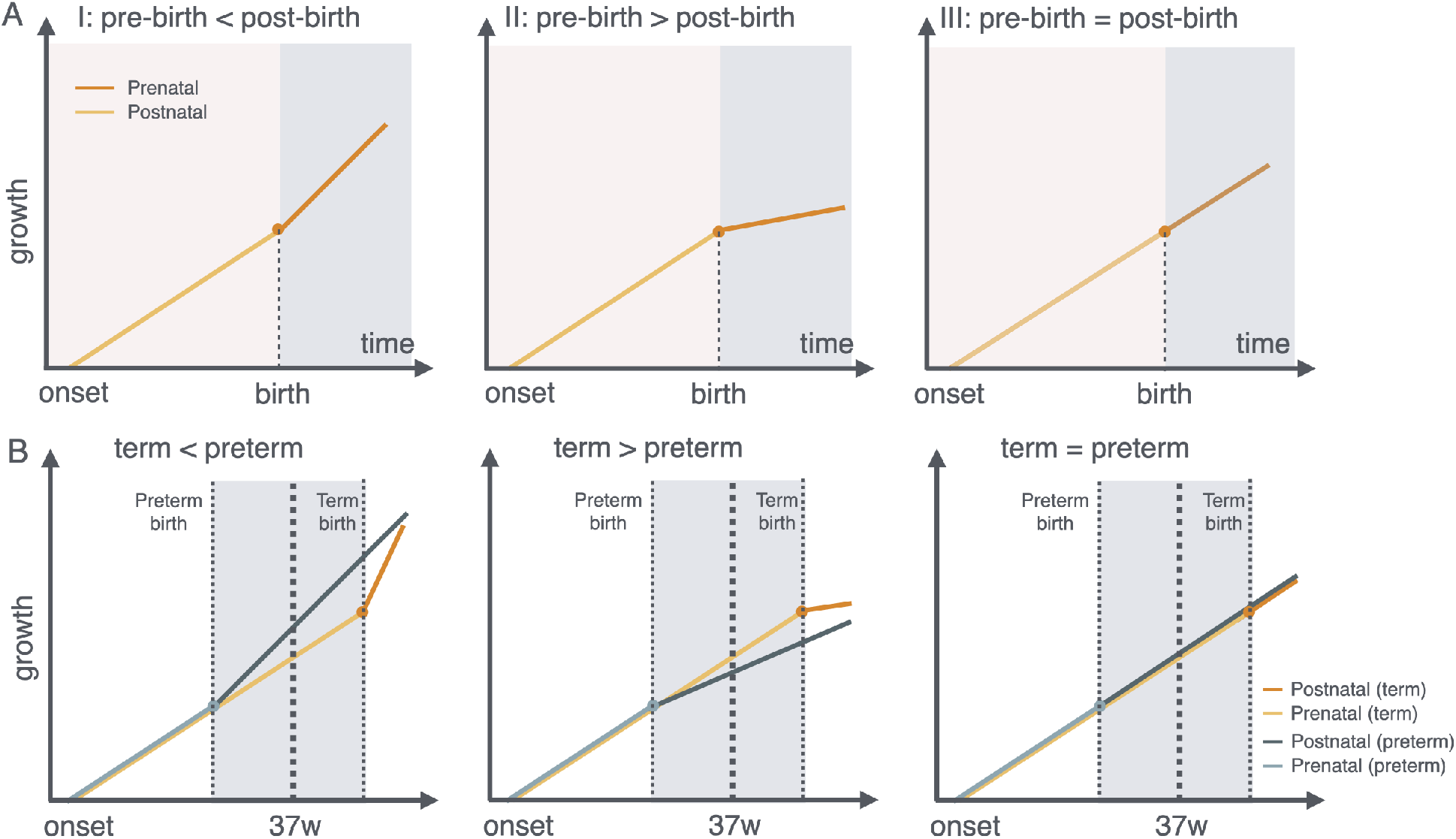
Hypotheses for the effect of birth on brain development. **(A)** Possible effects of birth on a developmental trajectory: (I) birth accelerates growth (acceleration hypothesis), (II) birth slows growth (protection hypothesis), or (III) birth has no effect (null hypothesis). **(B)** Expected trajectories of term and preterm infants under each hypothesis. Because preterm infants are born earlier, they would appear advanced, delayed or similar to term born infants at the same postmenstrual age, depending on which hypothesis holds.

These competing, or perhaps complementary, hypotheses become particularly important when considering preterm birth. Different hypotheses yield different pre- and postnatal growth trajectories for term and preterm newborns (Figure 1B). Under the acceleration hypothesis, preterm infants’ extended exposure to sensory stimulation might explain findings of advanced connectivity in certain domains. Under the protection hypothesis, early loss of the intrauterine environment would consistently impair development. The mixed findings in the literature, with some domains showing acceleration while others showing delays, suggest heterogeneous responses to the birth transition across different neural systems. Furthermore, the effect of birth may depend on the brain’s maturational state at the time of birth, so that it differs between preterm and term infants.

To characterise the impact of birth and preterm birth on neurodevelopmental trajectories, we developed a piecewise linear regression model that tracks perinatal brain development while modelling the effects of birth and prematurity separately across multiple measures of brain structure and connectivity. This approach specifically tests three developmental hypotheses regarding birth’s impact: whether it accelerates maturation, delays maturation, or produces no significant change in developmental trajectories. Using cross-sectional MRI data from term and preterm neonates in dHCP, we quantified developmental trajectories across volumetric, microstructural, and connectivity measures. We found that birth altered developmental trajectories in term infants in a measure- and region-specific manner. Volumetric growth accelerated after birth to varying degrees across brain structures. Microstructure continued to mature in deep white matter, although more slowly than before birth in most tracts, whereas superficial and cerebellar white matter reversed their prenatal trends. The change in cortical connectivity at birth followed the sensorimotor to association axis of cortical maturation (Dubois et al. 2014; Sydnor et al. 2021), shifting towards specialisation in sensory and insular cortex and towards more diverse connectivity in association cortex. In contrast, preterm infants largely maintained their prenatal trajectories, with only modest changes after birth. By separating the effects of birth and prematurity, this work helps reconcile contradictory findings in the literature and provides a framework for understanding perinatal brain maturation.

## RESULTS

### Asynchronous Volume Growth Trajectories of Brain Tissues

We characterised brain volume trajectories across the perinatal period using a piecewise linear Bayesian regression model. For each measure, the model estimates three parameters: (i) growth onset, i.e. the gestational age at which measurable expansion begins; (ii) the prenatal slope, the weekly growth rate before birth; and (iii) the postnatal slope, the weekly growth rate after birth. Since gestational age at birth and postmenstrual age at scan are decorrelated in the dHCP design, prenatal and postnatal rates can be estimated separately from postnatal scans alone (see Methods). We first applied the model to cortical grey matter (GM), white matter (WM) and total brain volumes. Although estimated only from postnatal data, the model-predicted growth rates aligned well with volume growths from independent fetal atlases at equivalent gestational ages (Figure 2B). Both prenatal and postnatal rates predicted by our model were validated in two ways: first by comparing to neonates scanned within days of birth, which approximates prenatal rates (Figure S1), and by examining a subset of infants with longitudinal scans, which aligns with our cross-sectionally estimated postnatal rates (Figure S2).

**Figure 2.**
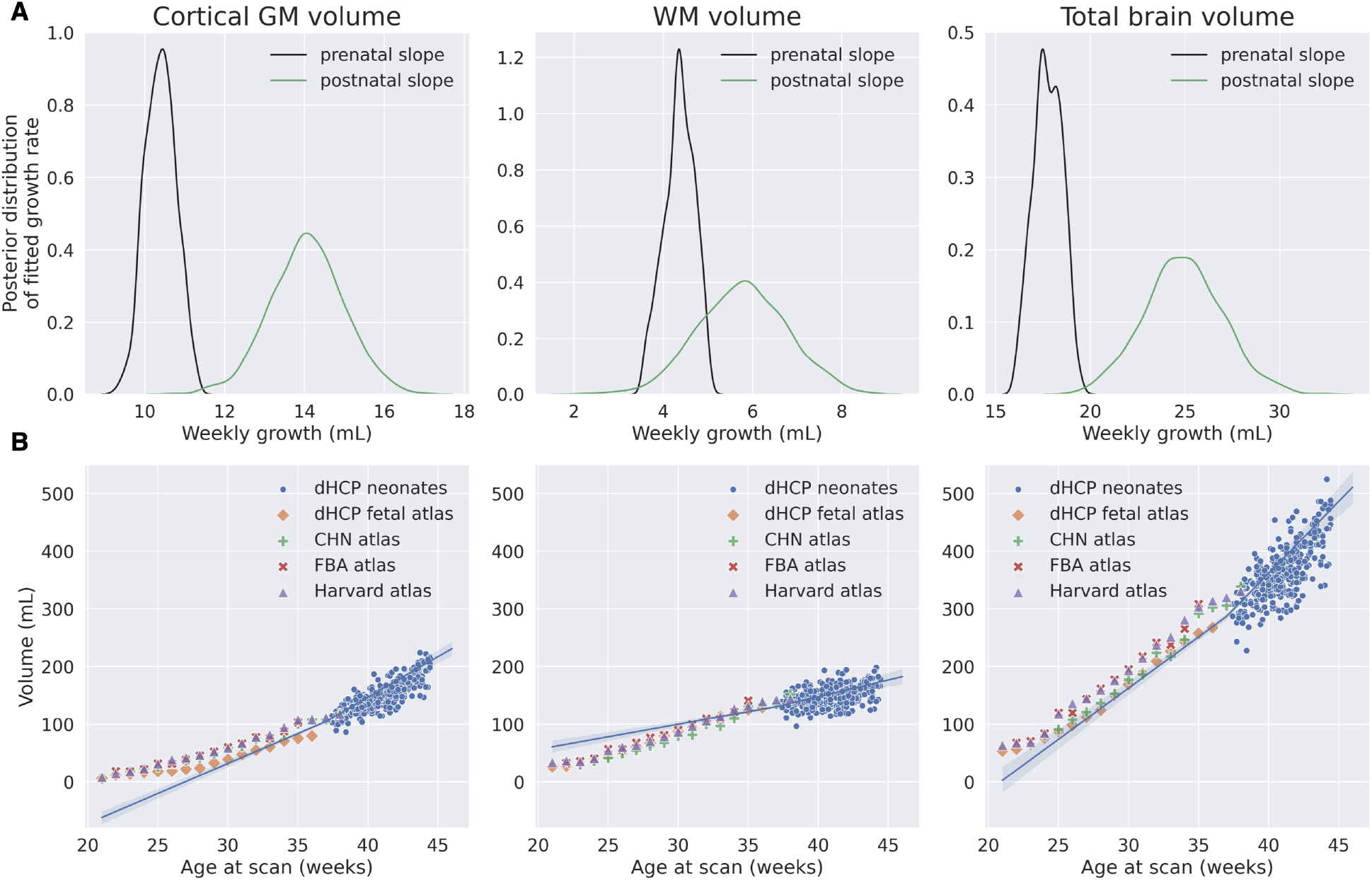
Prenatal and postnatal volume growth in term infants. **(A)** Posterior distributions of the prenatal and postnatal slopes (weekly growth rate) for cortical grey matter, white matter and total brain volume. **(B)** Fitted growth trajectories (blue curves). Blue dots show volumes of individual dHCP neonates; other dots show volumes from external fetal atlases, including the dHCP fetal atlas (Uus et al. 2023), the CHN atlas (Wu et al. 2021), the FBA atlas (Li et al. 2021), and the Harvard atlas (Gholipour et al. 2017).

Postnatal growth exceeded prenatal growth for all three measures (Figure 2A). Total brain volume grew by 17.4 mL/week before birth and 25.1 mL/week after birth, a 44% increase. These rates are consistent with previous estimates of 16.1 mL/week in utero (Gong et al. 1998) and approximately 1% per day shortly after birth (Holland et al. 2014), equivalent to about 24 mL/week. GM grew faster than WM both before and after birth, in line with reports that WM develops more slowly than cortical GM (Knickmeyer et al. 2008).

White matter fibre bundles showed similar patterns. Using FSL’s XTRACT (Warrington et al. 2020), we reconstructed 42 major bundles, comprising 19 bilateral association, limbic and projection tracts and 4 commissural tracts (Table S1). To allow comparison across tracts of different sizes, we normalised each tract’s volume by its adult volume, so that growth rates express weekly growth as a percentage of adult tract volume.

The fitted perinatal trajectories agreed with the observed tract volumes from dHCP neonates and with volumes derived from a series of independent fetal atlases (Figure 3A) (Gholipour et al. 2017; Wu et al. 2021; Li et al. 2021; Uus et al. 2023). Weekly volume growth differed across fibre categories (Figure 3B). In all tracts except the anterior thalamic radiation (ATR), volume grew faster after birth than before. Limbic tracts showed the largest postnatal acceleration, whereas association, projection and commissural tracts accelerated only modestly. This difference was also present in absolute tract volumes (Figure S3).

**Figure 3.**
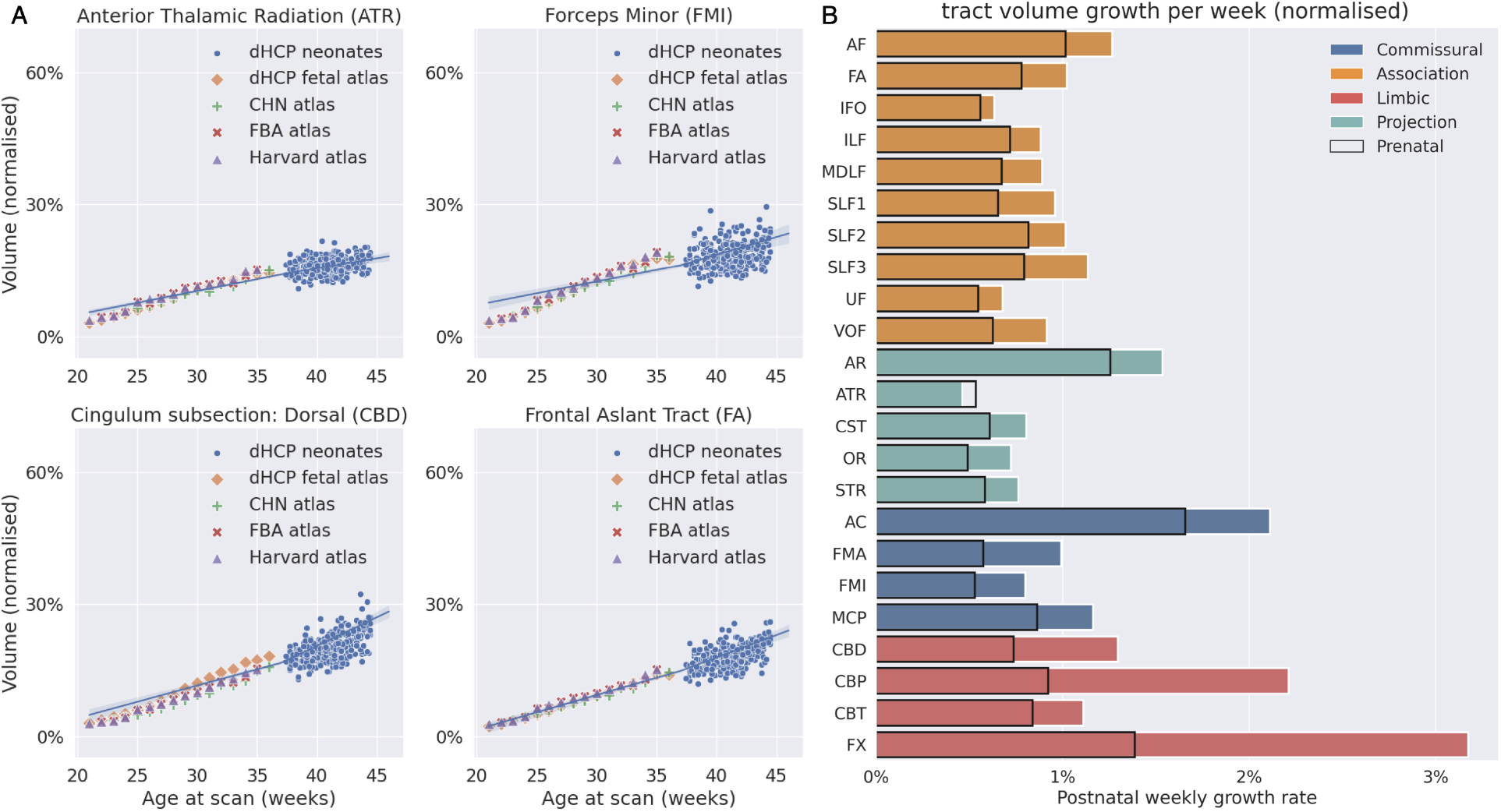
Prenatal and postnatal volume growth of white matter fibre bundles. **(A)** Example growth trajectories of four fibre bundles, one each from the commissural, association, limbic and projection systems. Blue curves show the piecewise linear model fitted to term infants in the dHCP. Blue dots show volumes of individual neonates; other dots show volumes from external fetal atlases, which were not used for model fitting. **(B)** Weekly growth rates of all fibre bundles, grouped by fibre system. Black: prenatal growth rate. Coloured: postnatal growth rate. See Figure S3 for the equivalent plots in absolute volumes.

### Preterm infants largely maintain prenatal growth rates after birth

We next asked whether the effect of birth differs in preterm infants. We extended the model to estimate separate postnatal slopes for term (≥37 weeks) and preterm (<37 weeks) infants. This allowed us to test directly whether birth accelerates, slows or does not change growth in each group. Unlike term infants, preterm infants largely maintained their prenatal growth rates after birth (Figure 4). Total brain volume grew only 7% faster after birth (17.4 to 18.6 mL/week) and GM only 5% faster (10.5 to 11 mL/week) (Figures 4B). The same contrast held across most fibre bundles. Limbic tracts were the exception. They also accelerated after birth in preterm infants, although much less than in term infants (Figures 4D). Because preterm infants’ postnatal data span the same postmenstrual ages as term infants’ prenatal data, this lack of acceleration also indicates that the acceleration in term infants reflects birth rather than a nonlinear effect of age.

**Figure 4.**
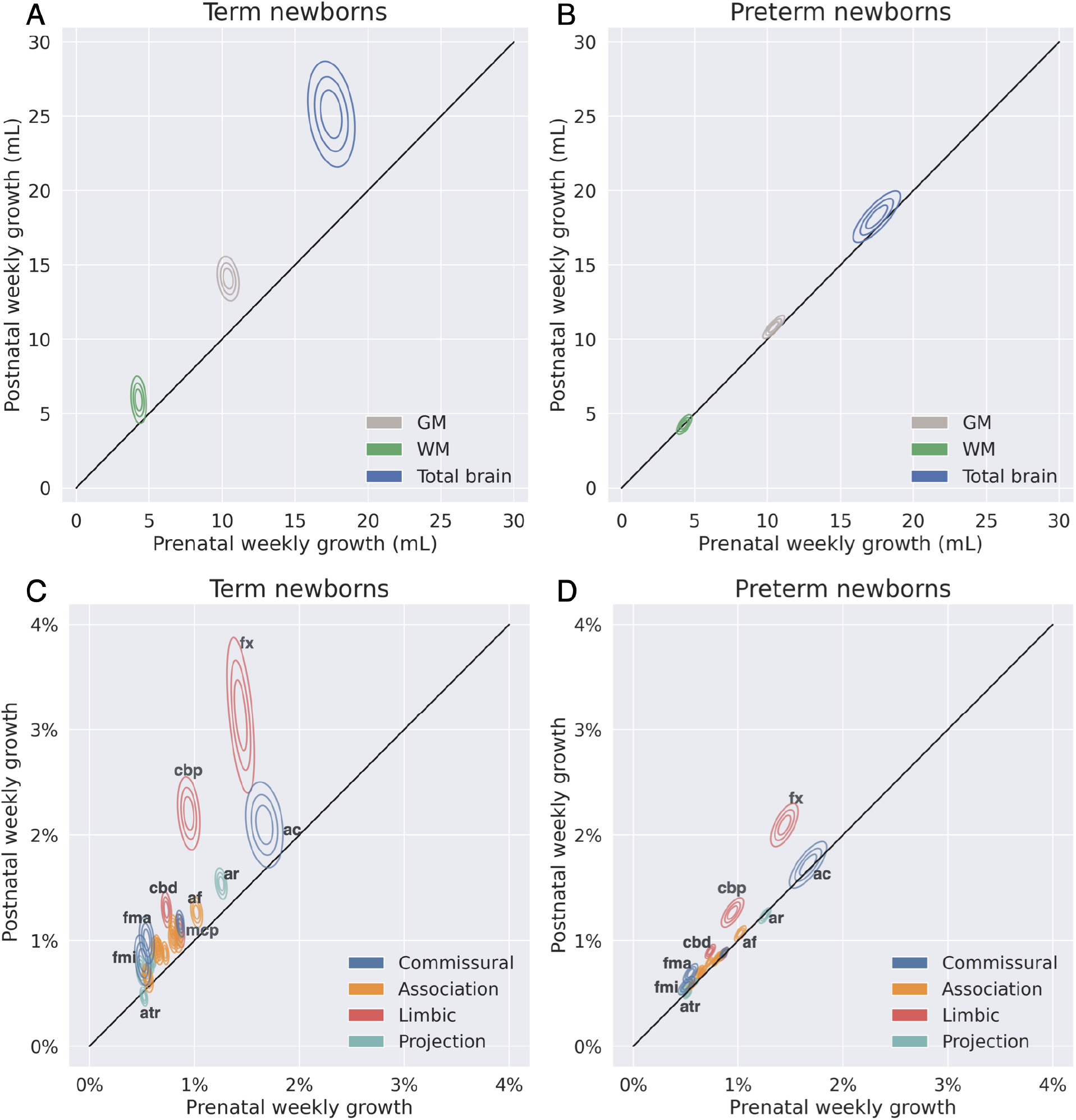
Prenatal and postnatal volume growth in term and preterm infants. **(A)** Prenatal and postnatal weekly growth rates (mL/week) of cortical grey matter, white matter and total brain volume in term infants. **(B)** Same as (A) for preterm infants. **(C)** Prenatal and postnatal weekly growth rates of white matter fibre bundles in term infants, normalised by the corresponding adult tract volume (% of adult volume per week). **(D)** Same as (C) for preterm infants. Contours show the posterior density of the prenatal (x axis) and postnatal (y axis) growth rates; points above the diagonal indicate faster growth after birth.

The absence of acceleration in preterm infants may indicate that the immature brain cannot yet support faster growth and continues at its intrauterine pace. The limbic exception parallels the stronger hippocampal and insular connectivity reported in preterm infants relative to fetuses, which has been attributed to early stress exposure (De Asis-Cruz et al. 2020).

### White Matter Microstructure Changes of Term and Preterm Newborns

To characterise microstructural development, we fitted diffusion tensors to the neonatal dMRI data using FSL DTIFIT and applied the piecewise linear model voxel-wise to fractional anisotropy (FA) and mean diffusivity (MD) maps. We describe the direction of change using the sign of each slope, and the effect of birth as the difference between postnatal and prenatal slopes. This analysis revealed spatially asynchronous prenatal and postnatal developmental trajectories across the brain for both term and preterm cohorts.

Prenatally, FA increased and MD decreased throughout most white matter (Figures 5A and 6A). In term infants, the effect of birth varied across white matter (Figures 5B and 6B). In deep white matter, FA continued to increase and MD continued to decrease after birth, in some voxels faster than before birth (Figure S5 and S6). In superficial white matter, however, FA reversed and decreased after birth. Deep cerebellar white matter also reversed, with FA decreasing and MD increasing after birth.

**Figure 5.**
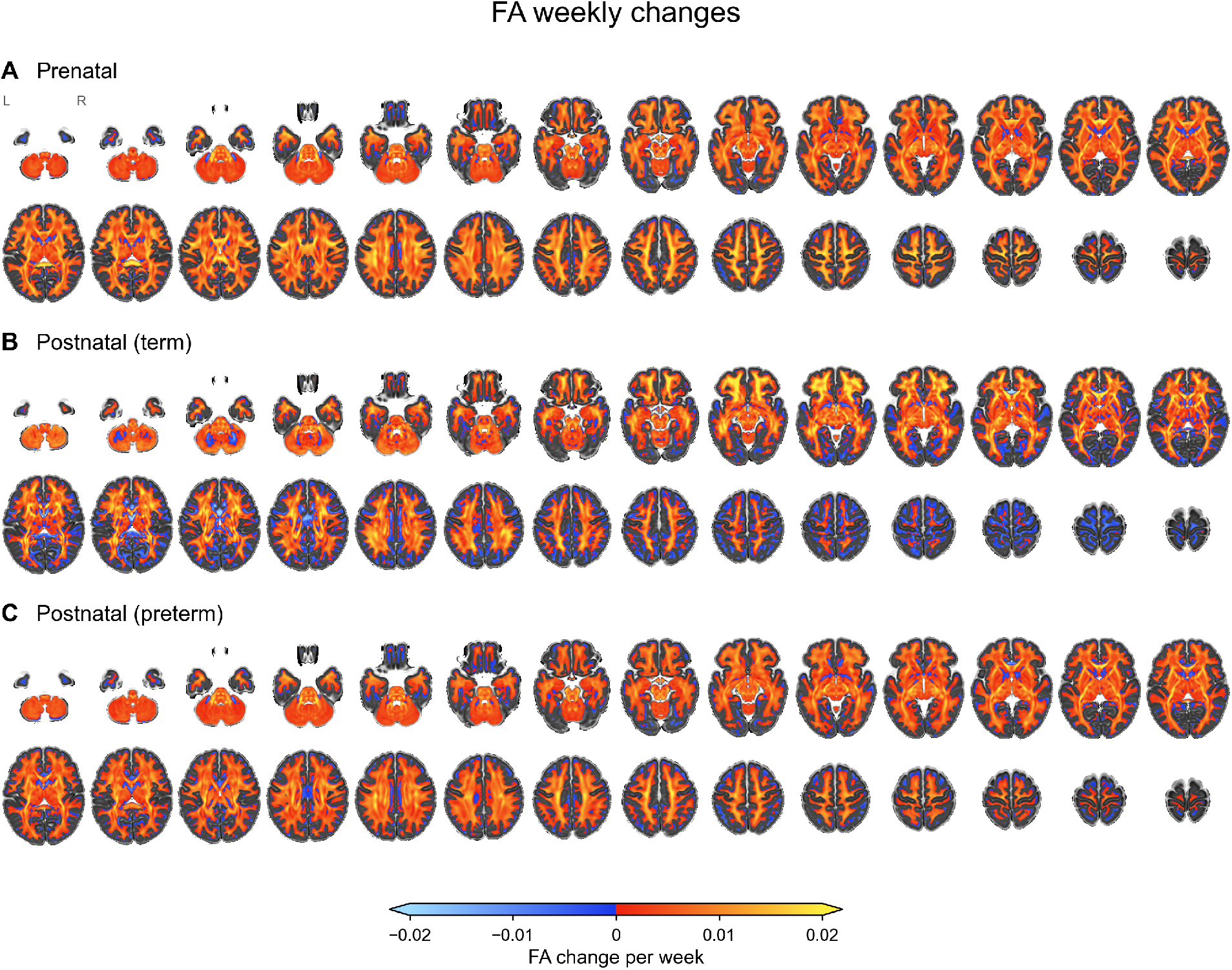
Prenatal and postnatal weekly change in FA. Voxelwise prenatal and postnatal slopes of FA for term and preterm newborns, shown on axial slices of the 40-week template (Schuh et al. 2018). Warm colours indicate increasing FA and cool colours decreasing FA. Maps are masked to white matter (Makropoulos et al. 2014; Makropoulos et al. 2016). **(A)** Prenatal slope. **(B)** Postnatal slope in term infants. **(C)** Postnatal slope in preterm infants. The prenatal slope is estimated jointly for term and preterm infants.

**Figure 6.**
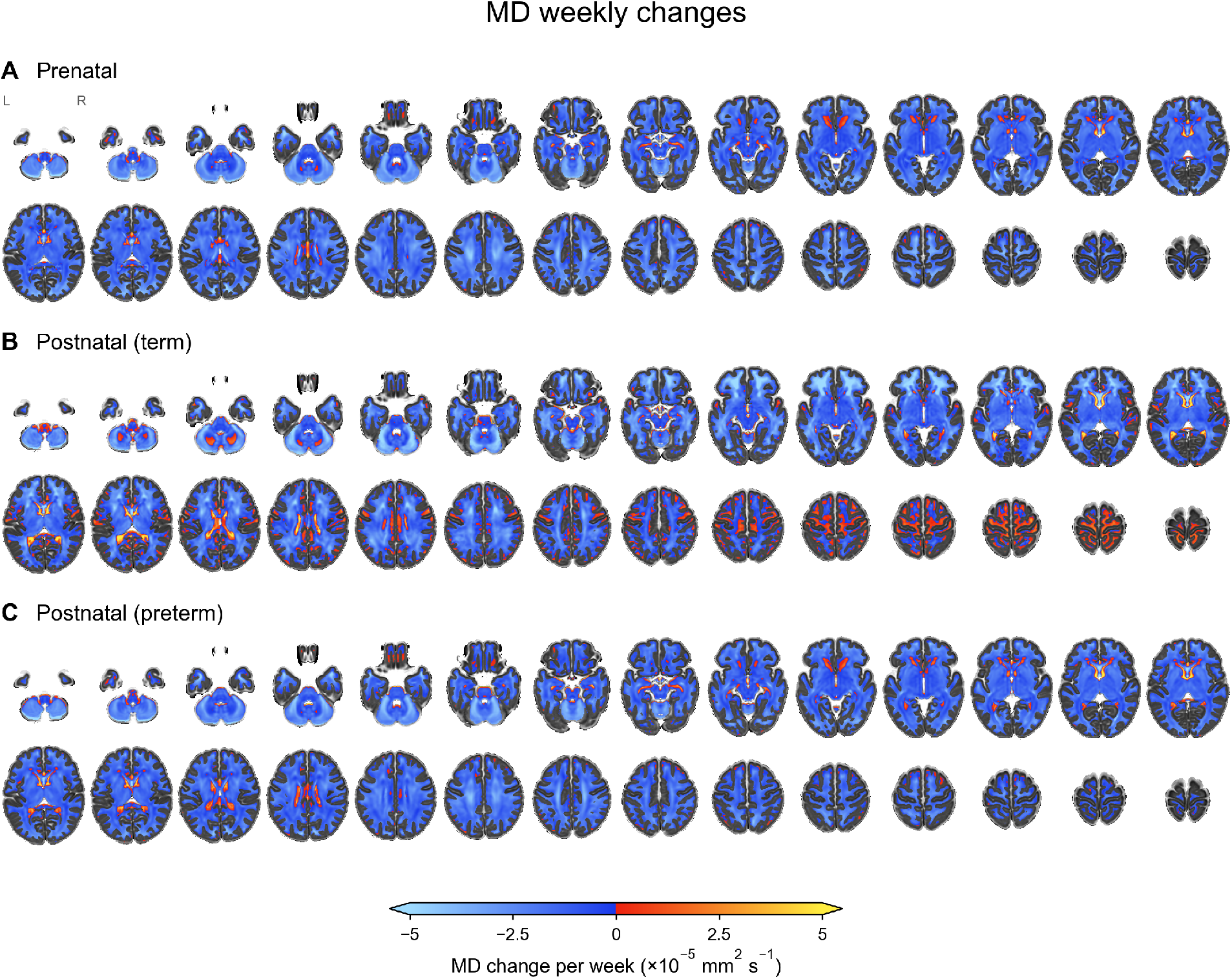
Prenatal and postnatal weekly change in MD. Voxelwise prenatal and postnatal slopes of MD for term and preterm newborns, shown on axial slices of the 40-week template (Schuh et al. 2018). Warm colours indicate increasing MD and cool colours decreasing MD. Maps are masked to white matter (Makropoulos et al. 2014; Makropoulos et al. 2016). **(A)** Prenatal slope. **(B)** Postnatal slope in term infants. **(C)** Postnatal slope in preterm infants. The prenatal slope is estimated jointly for term and preterm infants.

To examine whether these changes reflect fibre architecture, we quantified crossing fibre complexity. We fitted the ball and stick model with FSL BedpostX (Behrens et al. 2007; Jbabdi et al. 2012) to estimate volume fractions for up to three fibre populations per voxel, normalised the fractions to sum to one, and computed their Shannon entropy. Entropy ranges from 0, a single dominant fibre population, to log_2_ 3 ≈ 1.58, three equally weighted populations. Whereas FA can decrease for several reasons, including crossing fibres and low axonal density, entropy specifically measures how evenly the signal is distributed across fibre orientations.

Prenatally, entropy increased in most white matter (Figure 7A). In term infants, the effect of birth again varied by region (Figure 7B). Entropy reversed and decreased in regions dominated by parallel fibres, such as the corpus callosum, and increased faster in regions with extensive fibre crossing in the mature brain, such as parts of limbic and association pathways.

**Figure 7.**
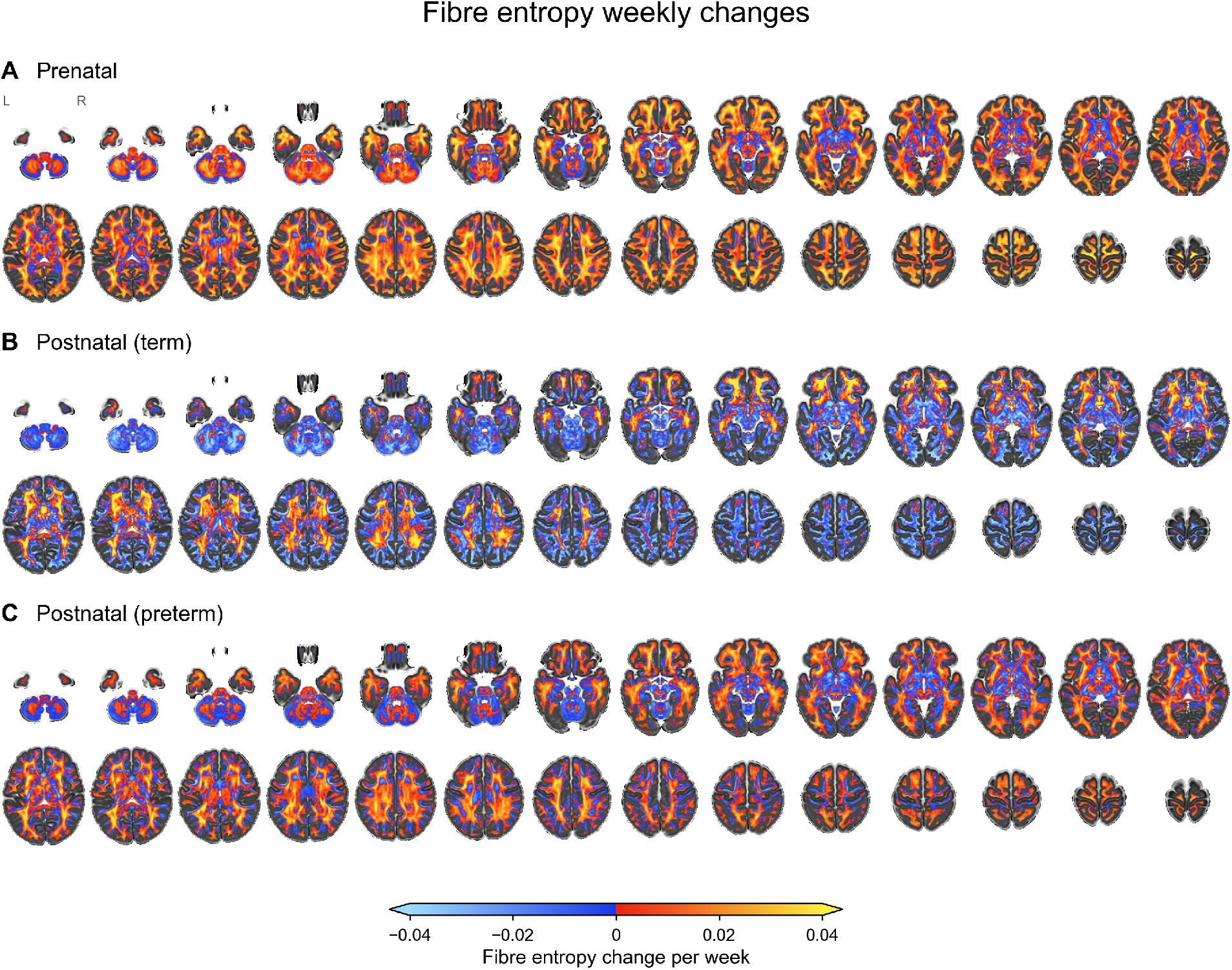
Prenatal and postnatal weekly change in crossing fibre entropy. Voxelwise slopes (change per week) of the Shannon entropy of fibre volume fractions estimated with the ball and stick model. Entropy ranges from 0 (a single dominant fibre population) to log_2_ 3 ≈ 1.58 (three equally weighted populations). Warm colours indicate increasing entropy and cool colours decreasing entropy. **(A)** Prenatal slope. **(B)** Postnatal slope in term infants. **(C)** Postnatal slope in preterm infants.

In preterm infants, the effect of birth was much smaller across all three measures (Figure 5C, 6C and 7C). FA, MD and entropy continued close to their prenatal rates, and none of the voxelwise reversals seen in term infants occurred. In superficial white matter, FA continued to increase, but more slowly than before birth. For entropy, postnatal changes shifted toward the term pattern without reaching it. The preterm brain therefore responded to birth, but far less than the term brain, consistent with the volumetric results.

We next examined FA and MD trajectories within the 42 tracts. Prenatally, FA increased and MD decreased in all tracts (Figures 8A and 8B). After birth, term infants showed two distinct patterns. In dorsal association tracts (AF, SLF1 to SLF3), most projection tracts (AR, CST, STR), commissural tracts other than FMI (AC, FMA, MCP) and most limbic tracts, FA increased and MD decreased more slowly than before birth. In contrast, ventral association tracts (IFO, ILF, UF), the optic radiation (OR), the anterior thalamic radiation (ATR) and the forceps minor (FMI) kept or exceeded their prenatal rate of FA increase, and their MD decreased faster after birth. These tracts may correspond to the deep white matter regions that changed faster after birth at the voxel level (Figures 5B and 6B). The remaining association tracts (frontal aslant, MdLF and VOF) showed a slower FA increase but a faster MD decrease. FA decreased after birth only in the dorsal (CBD) and perigenual (CBP) cingulum.

**Figure 8.**
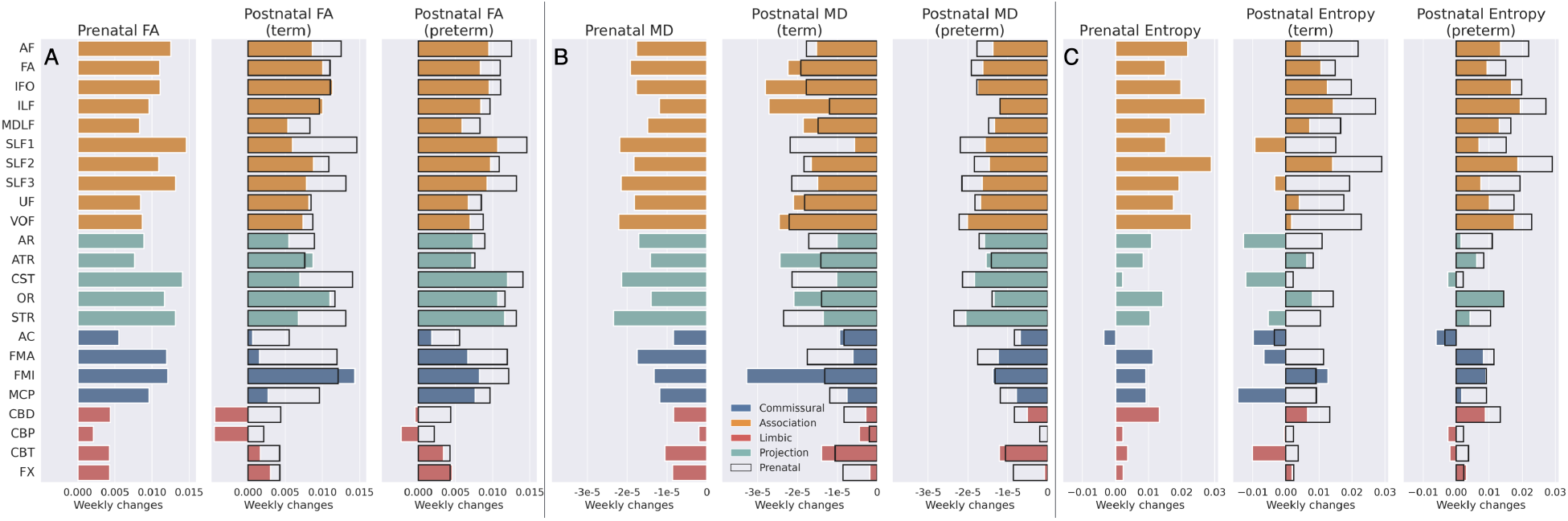
Prenatal and postnatal weekly change in microstructure of major fibre bundles. Weekly change in tract averaged **(A)** FA, **(B)** MD, and **(C)** crossing fibre entropy for each fibre bundle, grouped by fibre system. Bars show the prenatal slope and the postnatal slopes for term and preterm infants. Values are averaged over voxels within each tract.

Limbic tracts had the smallest rates of FA increase and MD decrease of the four systems, both before and after birth, although their volume accelerated most after birth (Figures 3B and 4C). Volume and microstructure can change independently. Limbic tracts may expand through axonal growth before becoming more coherent and myelinated. Such growth would increase volume without a comparable change in FA or MD.

Mean entropy within tracts also changed heterogeneously after birth (Figure 8C). In term infants, entropy reversed and decreased in three projection tracts (AR, CST, STR), two commissural tracts (FMA, MCP), two association tracts (SLF1, SLF3) and one limbic tract (CBT). In AC, entropy already decreased before birth and decreased faster afterwards. In the remaining tracts, entropy continued to increase at a similar or slower rate. FMI was the only tract in which entropy increased faster after birth. The faster increases seen at the voxel level (Figure 7B) are therefore confined to parts of tracts and do not dominate tract averages. Entropy reversed mainly in tracts whose FA and MD changes also slowed after birth.

In preterm infants, FA increased more slowly after birth than before in almost all tracts. In tracts that slowed in term infants, preterm infants mostly showed a milder form of the same deceleration in FA, MD and entropy. However, preterm infants did not show the postnatal acceleration seen in term infants. In ATR, OR, FMI, IFO and ILF, their rate of FA increase fell below the prenatal rate, and their rate of MD decrease stayed close to it. FA also decreased after birth in CBP. Overall, preterm infants stayed closer to their prenatal microstructural trajectory than term infants did, although they did not maintain it exactly.

### Developmental Trajectories of Fibre Innervation and Connectivity Specialisation

We next examined how white matter connectivity to the cortex develops during the perinatal period. Using probabilistic tractography, we constructed connectivity matrices between cortical vertices and white matter voxels (cortex × WM), and between white matter voxels and the 42 major fibre tracts (WM × tracts). Multiplying these matrices gave a cortical connectivity blueprint (cortex × tracts), where each row represents the innervation density, i.e., how strongly each tract reaches a cortical vertex (Mars et al. 2018). To quantify cortical connectivity specialisation, we calculated the Shannon entropy of each vertex’s tract innervation distribution (Figure 9A). High entropy indicates that a vertex is reached by many tracts in similar proportions (low specialisation); low entropy indicates that a few tracts dominate (high specialisation).

**Figure 9.**
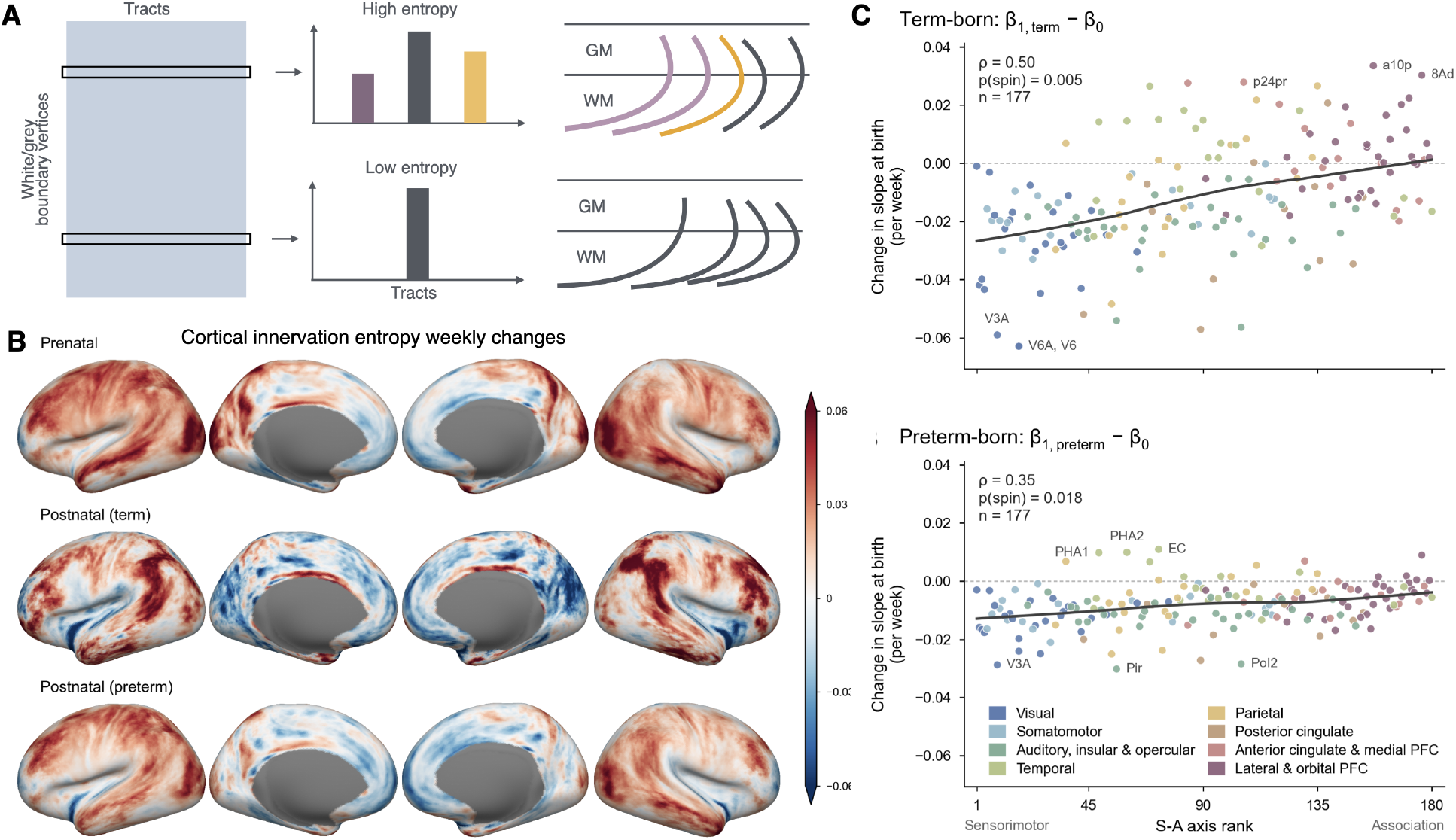
Development of cortical connectivity blueprints. **(A)** Illustration of connectivity entropy at each cortical vertex. Higher values indicate that the vertex is reached by a more even mix of tracts. **(B)** Prenatal and postnatal weekly changes in cortical connectivity entropy in term and preterm newborns. **(C)** Change in slope at birth (*β*_1,term_ − *β*_0_) for each area against its rank on the sensorimotor to association axis. *ρ* and *p*_spin_ from 5,000 spatial rotations.

Prenatally, cortical innervation entropy increased across most of the lateral surface and decreased across much of the medial surface (Figure 9B). After term birth, entropy increased most in lateral association cortex and decreased in the insula, dorsal visual cortex and V1. In preterm infants, the postnatal pattern remained close to the prenatal one.

To test whether the effect of birth followed the sensorimotor to association axis of cortical maturation (Sydnor et al. 2021), we correlated the change in slope at birth (*β*_1_ − *β*_0_) with each area’s rank on this axis. In term infants, the change at birth aligned with the axis (*ρ* = 0.50, *p*_spin_ = 0.005; Figure 9C). After birth, the rate of entropy change fell at the sensorimotor end of the axis, most in dorsal visual and posterior parietal areas (V6, V6A, V3A, V7) and the insula (PoI2), and rose at the association end, most in prefrontal and inferior parietal areas (a10p, 8Ad, a47r, PFm, IP2). The same alignment was present after preterm birth (*ρ* = 0.35, *p*_spin_ = 0.018). Neither the prenatal nor the term postnatal slopes aligned with the axis on their own (Figure S7, S8 and S9), so the alignment is specific to the change at birth.

Innervation slopes for all 42 tracts are provided in Figures S10 to S51. Because each tract’s cortical map sums to one, these slopes describe how terminations are redistributed across the cortex. For example, corticospinal terminations became concentrated over dorsal precentral and postcentral cortex after term birth (Figure S42 and S43).

## DISCUSSION

In this study, we used a piecewise linear model to separate the effects of birth and preterm birth on brain development, addressing a confound in previous neuroimaging studies of preterm infants. Analysing cross-sectional MRI data from term and preterm neonates in the dHCP, we found that birth altered developmental trajectories in term infants, but the direction of this effect depended on the measure and region. Volumetric growth accelerated after birth, most strongly in limbic tracts. Microstructure continued to mature in deep white matter with increasing FA and decreasing MD, although more slowly than before birth in most tracts. However, this pattern reversed in superficial and cerebellar white matter, where FA started to decrease and MD started to increase after birth. Birth also changed how cortical connectivity developed, and this change followed the sensorimotor to association axis, in which primary sensory and motor cortices mature earliest and association cortices supporting higher cognition follow more protracted timelines (Gogtay et al. 2004; Shaw et al. 2008; Valk et al. 2020; Dong et al. 2021; Sydnor et al. 2021; Larsen et al. 2023). After term birth, the rate of entropy change fell at the sensorimotor end of the axis, most in dorsal visual and insular cortex, and rose at the association end, most in prefrontal and inferior parietal cortex. A shift towards specialisation in sensory cortex is consistent with the elimination of exuberant axonal projections during early development (Innocenti and Price 2005). Notably, neither the prenatal nor the postnatal rates aligned with the axis on their own; the alignment appeared only in their difference. Evidence for this axis comes largely from childhood and adolescence (Tau and Peterson 2009; Deoni et al. 2015), with emerging evidence in infancy (Natu et al. 2021; Nishio et al. 2025). Our results suggest that the axis already organises the cortical response to birth before it is evident in developmental rates themselves.

In contrast to term newborns, preterm infants largely maintained prenatal developmental trajectories after birth across volumetric, microstructural and connectivity measures, as if development continued in utero. However, preterm brains still show partial responsiveness. For example, limbic tracts accelerated in volume, although much less than in term infants. Microstructure still changed modestly after birth. The change in cortical connectivity followed the same sensorimotor to association axis as in term infants. The preterm brain may therefore respond to the extrauterine environment, but within maturational constraints. This offers an explanation for reports of accelerated maturation after preterm birth. If birth slows or reverses development in the more mature brain while the preterm brain continues at its prenatal rate, preterm infants will appear more “advanced” at matched age without any acceleration of their own. For example, infants born very preterm are reported to show larger visual and smaller tactile evoked responses than infants born late preterm at the same postmenstrual age, which has been interpreted as accelerated sensory maturation (Schmidt Mellado et al. 2022). Visual responses increase and tactile responses decrease with age. If birth slows these changes in late preterm infants, who are born closer to term, their visual responses would continue to increase and their tactile responses to decrease after birth, but more slowly than before. Very preterm infants may continue at their prenatal rates, as a result showing larger visual and smaller tactile responses at the same postmenstrual age without any birth-triggered alteration of their own. Separating the effect of birth from that of prematurity may therefore help reconcile contradictory reports in the literature.

The prenatal increase in white matter FA and decrease in MD are consistent with increasing axonal organisation and the onset of myelination (Dubois et al. 2014). In post mortem adult tissue, FA increases and MD decreases with axonal density (Schmierer et al. 2007). After term birth, these changes continued in deep white matter but slowed in most tracts, in line with slower T1w/T2w growth ex utero than in utero (Grotheer et al. 2023). In contrast, FA reversed in superficial white matter, which contains short association fibres and remnants of the transient subplate zone and undergoes extensive reorganisation around term (Kostović and Jovanov-Milošević 2006). Because superficial white matter borders the cortex, partial volume effects could also contribute to this reversal as the cortex expands after birth. Deep cerebellar white matter also reversed after birth. The cerebellum grows particularly rapidly around term (Limperopoulos et al. 2005), but diffusion measures in the neonatal cerebellum are sensitive to noise and partial volume effects, so this finding should be interpreted with caution.

Limbic tracts showed the largest postnatal volumetric acceleration but the smallest rates of FA increase and MD decrease, both before and after birth. Volume and diffusion measures capture different aspects of development. Volume reflects the amount of tissue, whereas FA and MD reflect its organisation and water content. Limbic tracts may therefore grow through axonal elongation and the addition of fibres before becoming more coherent and myelinated.

Note that we fitted our models exclusively to postnatal data despite the availability of fetal scans in dHCP. Fetal MRI faces unique challenges (Rajagopalan et al. 2021; Vahedifard et al. 2023). For example, the intrauterine environment and lower SNR necessitate different acquisition protocols. The maternal and fetal motion requires specialised motion correction algorithms. Furthermore, immature tissue contrast in fetal scans also poses significant challenges to algorithms that work on neonates. These differences extend to subsequent preprocessing procedures, introducing further technical variations to the true anatomical variability. Therefore, combining fetal and neonatal data in a single model fitting will introduce bias, potentially misattributing protocol-driven measurement differences to biological trajectories. However, our approach of fitting models to postnatal data alone while validating against external fetal atlases (Gholipour et al. 2017; Wu et al. 2021; Li et al. 2021; Uus et al. 2023) provides evidence of model validity. The close alignment between our predicted prenatal trajectories and atlas-derived fetal volumes for most measures demonstrates that the framework captures true developmental processes rather than protocol-driven heterogeneity. Two additional model validations further support this approach. First, we selected a set of babies that were scanned very shortly after birth, but at different gestational ages. This gave us an approximation to pre-birth growth trajectories which matched our model predictions very well (Figure S1). Second, we compared postnatal growth rates to longitudinal trajectories in a subset of babies to validate post-birth slopes and found agreement (Figure S2).

These findings suggest that the transition to extrauterine life does not uniformly accelerate brain maturation, but that the developmental response to birth depends on the maturational state of the brain. While birth at term is associated with widespread changes in developmental trajectories, earlier exposure to the extrauterine environment appears insufficient to trigger comparable changes in preterm infants. Nevertheless, the selective responsiveness of limbic pathways and the alignment of cortical connectivity changes with the sensorimotor to association axis in both groups, suggest that sensitivity to the extrauterine environment varies across neural systems and their stages of maturation.

An important implication is that the maintenance of prenatal developmental trajectories following preterm birth should not be interpreted as delayed or impaired maturation. Instead, these findings suggest that ex utero brain development reflects an interaction between intrinsic maturational processes and environmental influences, with their relative contributions varying across neural systems and across gestational age. Integrating structural and functional measures will be important for determining if these differing developmental trajectories have lasting impact on brain function.

## METHODS

### Subjects and Image Pre-processing

We analysed T2-weighted and Diffusion MRI (dMRI) data from the developing Human Connectome Project (dHCP) (Edwards et al. 2022), including 330 term-born neonates (38-44 weeks postmenstrual age) and 118 preterm neonates (45 scanned at term-equivalent age). T2w images were reconstructed at 0.5 mm isotropic resolution using retrospective motion correction combining sagittal and axial acquisitions (Kuklisova-Murgasova et al. 2012; Cordero-Grande et al. 2017). We used the dHCP pipeline v1.1 (Makropoulos et al. 2018) for subsequent bias correction and tissue segmentation.

Diffusion images were corrected for motion, eddy currents, and susceptibility distortions using FSL topup and EDDY (Andersson et al. 2003; Andersson and Sotiropoulos 2016), as implemented in the dHCP diffusion pipeline (Bastiani et al. 2019), and then reconstructed at 1.5 mm isotropic resolution from 3 mm overlapping slices (Bastiani et al. 2019). We registered the dMRI data to the T2w structural images using boundary-based registration (Greve and Fischl 2009). Diffusion tensors were fitted with FSL DTIFIT using the b=0,400 and 1000 s/mm^2^ volumes.

To register individual brain images to a 40-week template (Schuh et al. 2018), we used FSL’s MultiModal Registration Framework (MMORF) (Lange et al. 2024). MMORF registers the T2w image and the diffusion tensor simultaneously, combining anatomical contrast with white matter orientation, which improves the alignment of major fibre tracts compared with registration of T2w images alone (Lange et al. 2024).

### Tissue Volumetry

We obtained cortical GM, WM and CSF volumes by segmenting T2w images using Draw-EM (Makropoulos et al. 2014; Makropoulos et al. 2016), both for the individual neonates and the fetal atlases (Uus et al. 2023; Wu et al. 2021; Li et al. 2021; Gholipour et al. 2017).

To calculate fibre bundle volumes, we adopted an atlas-based approach instead of direct tractography. This is because neonatal brains have incomplete white matter maturation (especially in preterm newborns), making tractography often fail to accurately delineate immature pathways and thus underestimate fibre bundle volumes. We reconstructed 42 major fibre bundles in each subject with XTRACT (Warrington et al. 2020), warped each reconstruction to the dHCP template at 40 weeks (Schuh et al. 2018) and averaged across subjects, which reduces the influence of incomplete reconstructions in individual subjects. The resulting atlas was warped back into each subject’s native space using the inverse of the nonlinear transform estimated with MMORF (Lange et al. 2024). Bundle volume was the number of voxels in the warped mask multiplied by the voxel volume.

To compare neonatal volumes with fetal development, we measured bundle volumes in the same way in four independent fetal atlases (**?**; Wu et al. 2021; Li et al. 2021; Gholipour et al. 2017). Each weekly fetal template was registered to the 40 week template with MMORF, and the neonatal tract atlas was warped into the fetal template space where we counted voxels.

Finally, we expressed each bundle volume as a fraction of the corresponding bundle volume in the adult XTRACT atlas (Warrington et al. 2020). This places bundles of very different sizes on a common scale, so that growth rates can be compared across bundles.

### Microstructure Properties

We calculated FA and MD maps in native space using FSL DTIFIT. Fibre orientation distributions were estimated using FSL BedpostX (Behrens et al. 2007; Jbabdi et al. 2012), modelling up to three fibre populations per voxel with their associated volume fractions (i.e., the proportion of diffusion signal from each fibre population). We quantified local fibre configuration complexity using Shannon entropy of the signal fraction distribution across the three modelled fibre orientations in each voxel. This entropy metric provides a continuous measure of crossing fibre complexity: high entropy values (approaching log_2_(3) ≈ 1.58 for three equally-weighted populations) indicate that signal is distributed equally across all three fibre populations, reflecting complex multi-directional crossing patterns. Conversely, low entropy values (approaching 0) indicate that one fibre population strongly dominates the signal, suggesting a simpler fibre architecture with either a single coherent bundle or a primary fibre direction with minimal crossing.

#### Alignment with the sensorimotor to association axis

We averaged the posterior mean slopes within areas of the HCP multimodal parcellation (Glasser et al. 2016) separately for each hemisphere. Vertices without estimates were excluded. A hemisphere’s area mean was used only if at least 30% of the area’s vertices had estimates. Left and right means were then averaged, otherwise the single available hemisphere was used. Three areas failed this criterion in both hemispheres, leaving 177. For each map, we computed the Spearman correlation with the sensorimotor to association axis rank (Sydnor et al. 2021) and assessed significance with a spin test (Alexander-Bloch et al. 2018) using 5,000 rotations, assigning each area the value of the nearest rotated area. Because our hypothesis concerned the effect of birth, the primary tests used the change in slope at birth for term and preterm infants (*β*_1,term_ − *β*_0_ and *β*_1,preterm_ − *β*_0_), with Bonferroni correction for these two tests. Correlations for the individual slope maps are reported for comparison.

### Developmental Innervation and Blueprints

To characterise cortical-white matter connectivity development, we built two connectivity matrices per subject using probabilistic tractography. The first matrix, C (cortex × white matter), quantified connectivity between cortical vertices and white matter voxels. The second matrix, T (white matter × tracts), characterised relationships between white matter voxels and the 42 major fibre bundles. Multiplying these matrices (B = C × T) yielded a cortical blueprint matrix B (cortex × tracts). Each column of B, representing a tract’s cortical terminations, was normalised to sum to one, so that innervation density describes the proportion of a tract’s terminations at each vertex. Each row then represents the “connectivity fingerprints” of a single cortical vertex, i.e., the distribution of streamline densities to each of the 42 major white matter fibre bundles (Mars et al. 2018). To calculate entropy, each fingerprint was further normalised to sum to one.

To quantify cortical specialisation development, we calculated the entropy of each vertex’s tract distribution (i.e., each row of B). High entropy indicates that a cortical region receives relatively equal innervation from multiple tracts (i.e., low specialisation), while low entropy values indicate that innervation is dominated by one or a few specific tracts (i.e., high specialisation). This provides a continuous metric of cortical connectivity specialisation that can be mapped across the cortex and compared across developmental stages.

### Piecewise Linear Regression Model

We developed a piecewise linear model to characterise brain development trajectories across the perinatal transition. The model describes a brain measure (e.g., brain tissue volume), *y*(*t*), as a function of age at scan (*t*_scan_, in weeks) and age at birth (*t*_birth_):

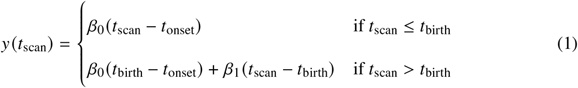

where *β*_0_ represents the prenatal growth rate (changes per week), *β*_1_ represents the postnatal growth rate, and *t*_onset_ is the gestational age at which measurable development begins. This formulation allows us to estimate separate pre- and postnatal growth rates despite only having postnatal scan data. This is possible because gestational age at birth (*t*_birth_) and age at scan (*t*_scan_) are decorrelated by experimental design in dHCP, i.e. infants are born at different gestational ages but scanned at similar postnatal ages, and vice versa. Therefore, the model simultaneously fits both the prenatal trajectories (reflected in volume differences among infants born at different gestational ages when scanned at the same postnatal age) and the postnatal trajectories (reflected in volume changes across postnatal ages within individual infants or across infants scanned at different postnatal ages).

The model can be extended to estimate separate postnatal growth rates for each group:

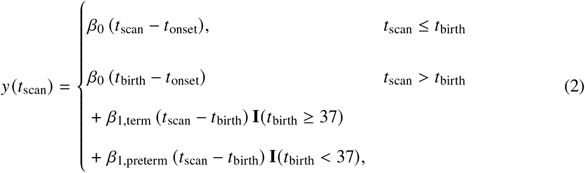

where *β*_1,term_ represents the postnatal growth rate for term-born infants (≥37 weeks gestational age) and *β*_1,preterm_ represents the postnatal growth rate for preterm infants (<37 weeks gestational age). The indicator functions **I**() ensure that each infant contributes to the appropriate growth rate parameter based on their gestational age at birth. The prenatal slope (*β*_0_) and onset age (*t*_onset_) remain shared across both groups, reflecting the assumption that prenatal development follows the same trajectory for all fetuses prior to birth. The posterior parameter distributions were estimated using Markov Chain Monte Carlo (MCMC) sampling with a Metropolis-Hastings algorithm. We used flat priors constrained to physiologically plausible bounds (e.g., onset time between 0-40 weeks postmenstrual age). For each parameter, we drew 10,000 MCMC samples after a 1,000-iteration burn-in, retaining every 10th sample to reduce autocorrelation, yielding 1,000 posterior samples.

## DATA AVAILABILITY

The neonatal MRI data used in this study are from dHCP and are available to researchers who agree to its data use terms. Group-level maps of all estimated parameters (growth onset, prenatal and postnatal slopes for term and preterm infants) for tissue volumes, FA, MD, fibre entropy and cortical innervation are available on OSF at https://osf.io/n6rav/overview.

## ACKNOWLEDGEMENT

This project was supported by the NIHR Oxford Health Biomedical Research Centre (NIHR203316). The Centre for Integrative Neuroimaging was supported by core funding from the Wellcome Trust (203139/Z/16/Z and 203139/A/16/Z). SJ is supported by a Wellcome Senior Fellowship (221933/Z/20/Z) and Wellcome Collaborative Award (215573/Z/19/Z).

## SUPPLEMENTARY MATERIALS

**TABLE S1.** The list of 42 fibre bundles, including 19 bilateral and 4 commissural (Warrington et al. 2020).

|  | **Tract** | **Abbreviation** |
|---|---|---|
| **Association Fibres** | Arcuate Fasciculus | AF |
|  | Frontal Aslant Tract | FA |
|  | Inferior Longitudinal Fasciculus | ILF |
|  | Inferior Fronto-Occipital Fasciculus | IFO |
|  | Middle Longitudinal Fasciculus | MdLF |
|  | Superior Longitudinal Fasciculus I | SLF1 |
|  | Superior Longitudinal Fasciculus II | SLF2 |
|  | Superior Longitudinal Fasciculus III | SLF3 |
|  | Uncinate Fasciculus | UF |
|  | Vertical Occipital Fasciculus | VOF |
| **Commissural Fibres** | Anterior Commissure | AC |
|  | Forceps Major | FMA |
|  | Forceps Minor | FMI |
|  | Middle Cerebellar Peduncle | MCP |
| **Limbic Fibres** | Cingulum subsection: Dorsal | CBD |
|  | Cingulum subsection: Peri-genual | CBP |
|  | Cingulum subsection: Temporal | CBT |
|  | Fornix | FX |
| **Projection Fibres** | Acoustic Radiation | AR |
|  | Anterior Thalamic Radiation | ATR |
|  | Corticospinal Tract | CST |
|  | Optic Radiation | OR |
|  | Superior Thalamic Radiation | STR |

**Figure S1.**
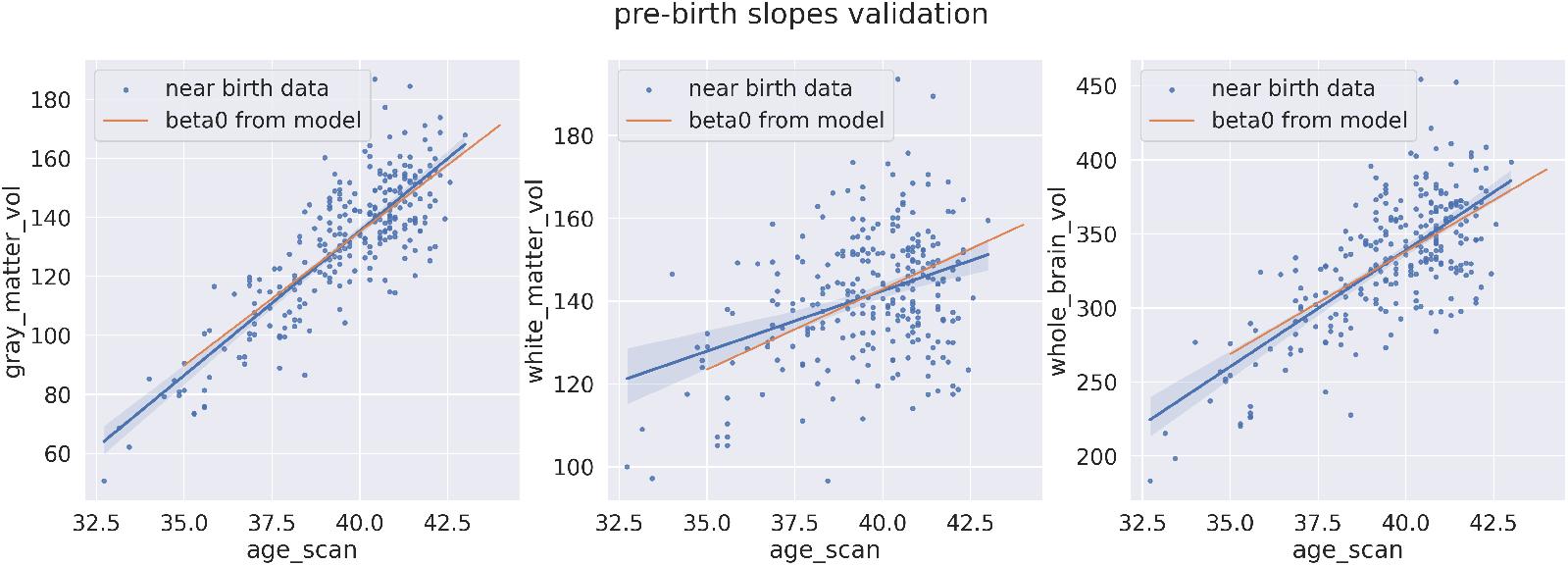
Modelled prenatal slopes agreed with the actual prenatal weekly growth rate estimated from near-birth neonates (scanned <1 week after birth).

**Figure S2.**
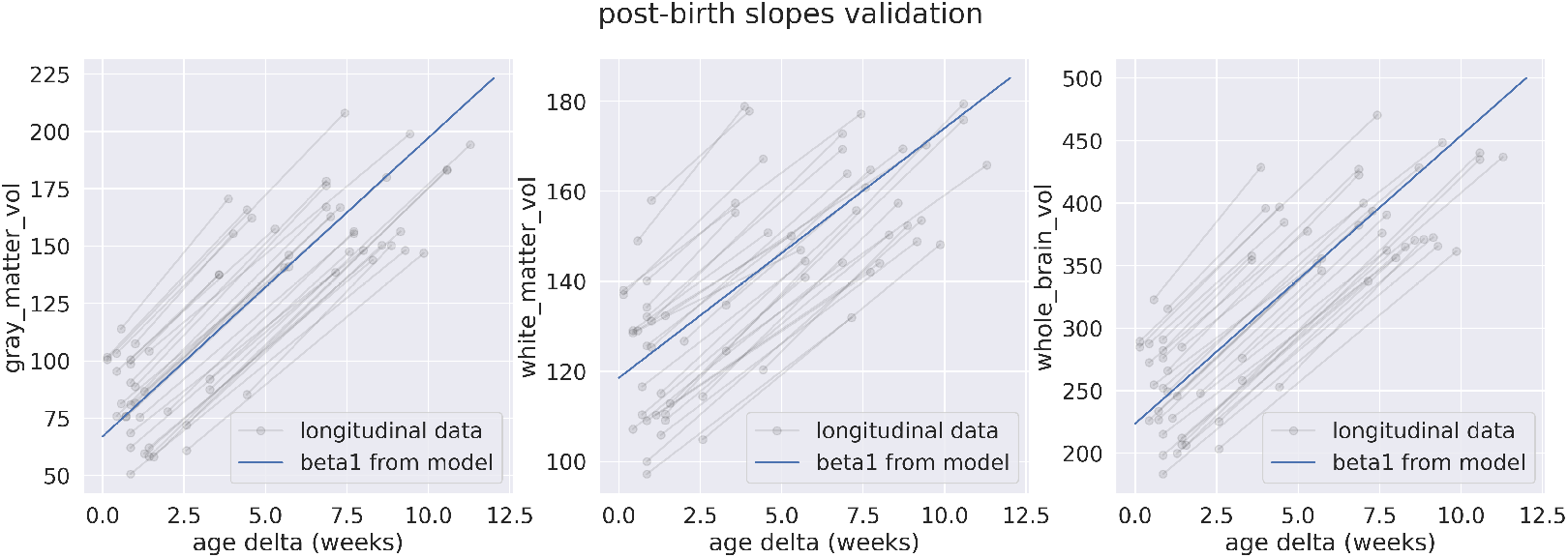
Modelled preterm postnatal slopes agreed with the actual postnatal weekly growth rate estimated from longitudinal scans of preterm newborns.

**Figure S3.**
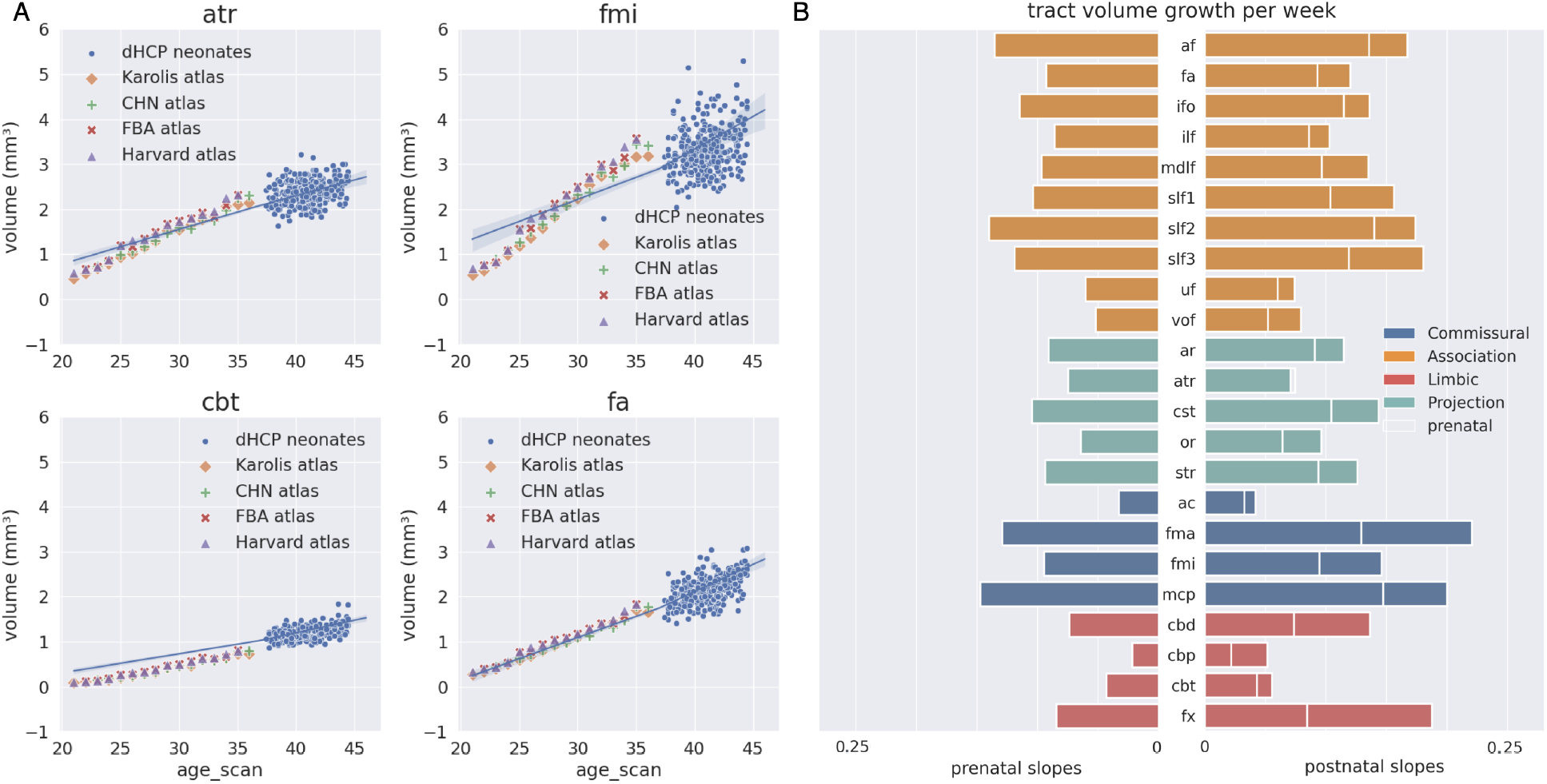
Equivalent plot of Figure 3, where each tract’s volume is un-normalised by its adult volume, so that slopes represent weekly change in absolute volume. See S4 for trajectories of all 42 tracts.

**Figure S4.**
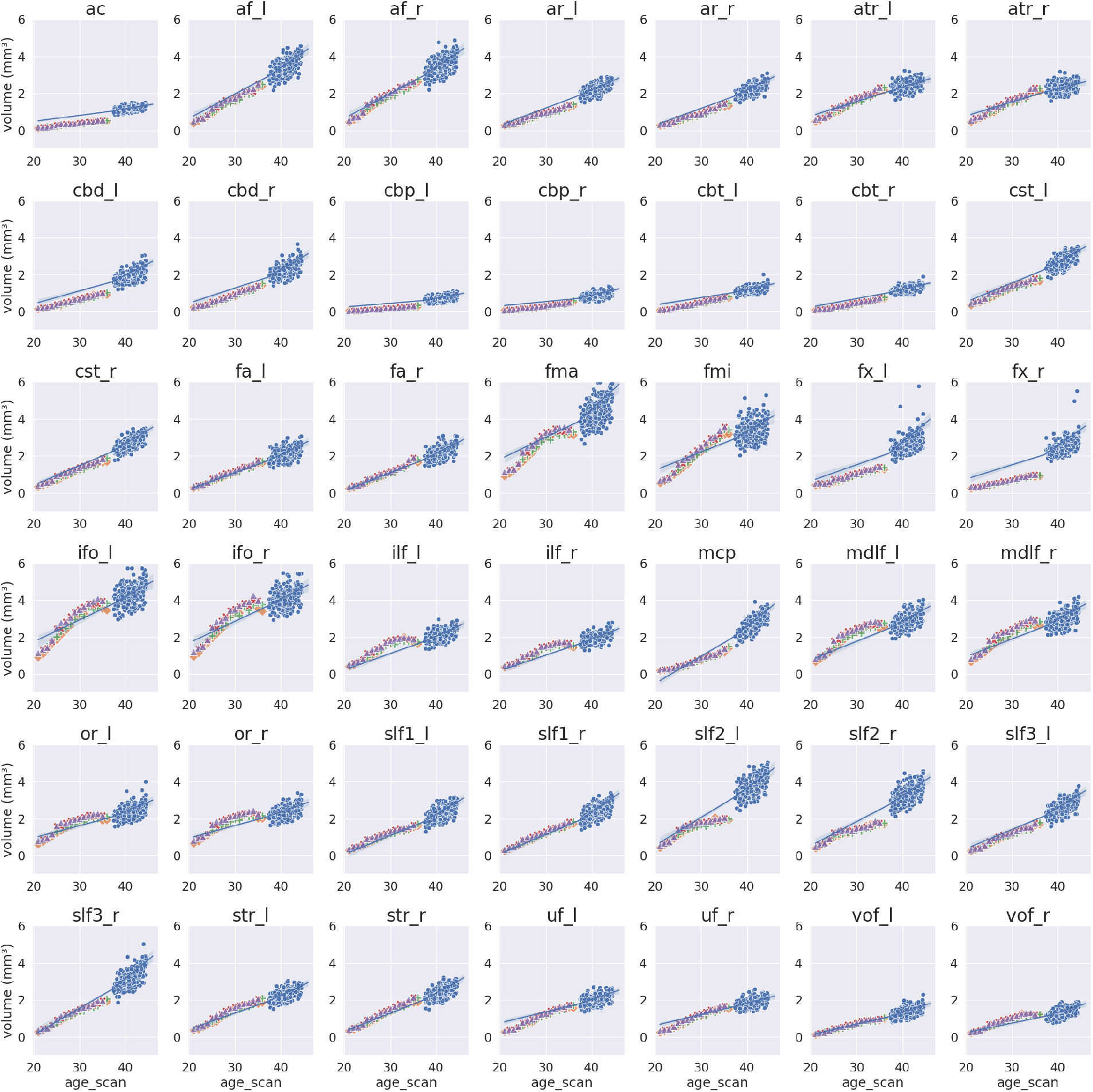
Volume growth trajectories of all 42 white matter fibre bundles. Absolute tract volume as a function of postmenstrual age (weeks). Blue curves show the piecewise linear model fitted to term infants in the dHCP. Blue dots show volumes of individual neonates; other dots show volumes from independent fetal atlases as in Figure 3.

**Figure S5.**
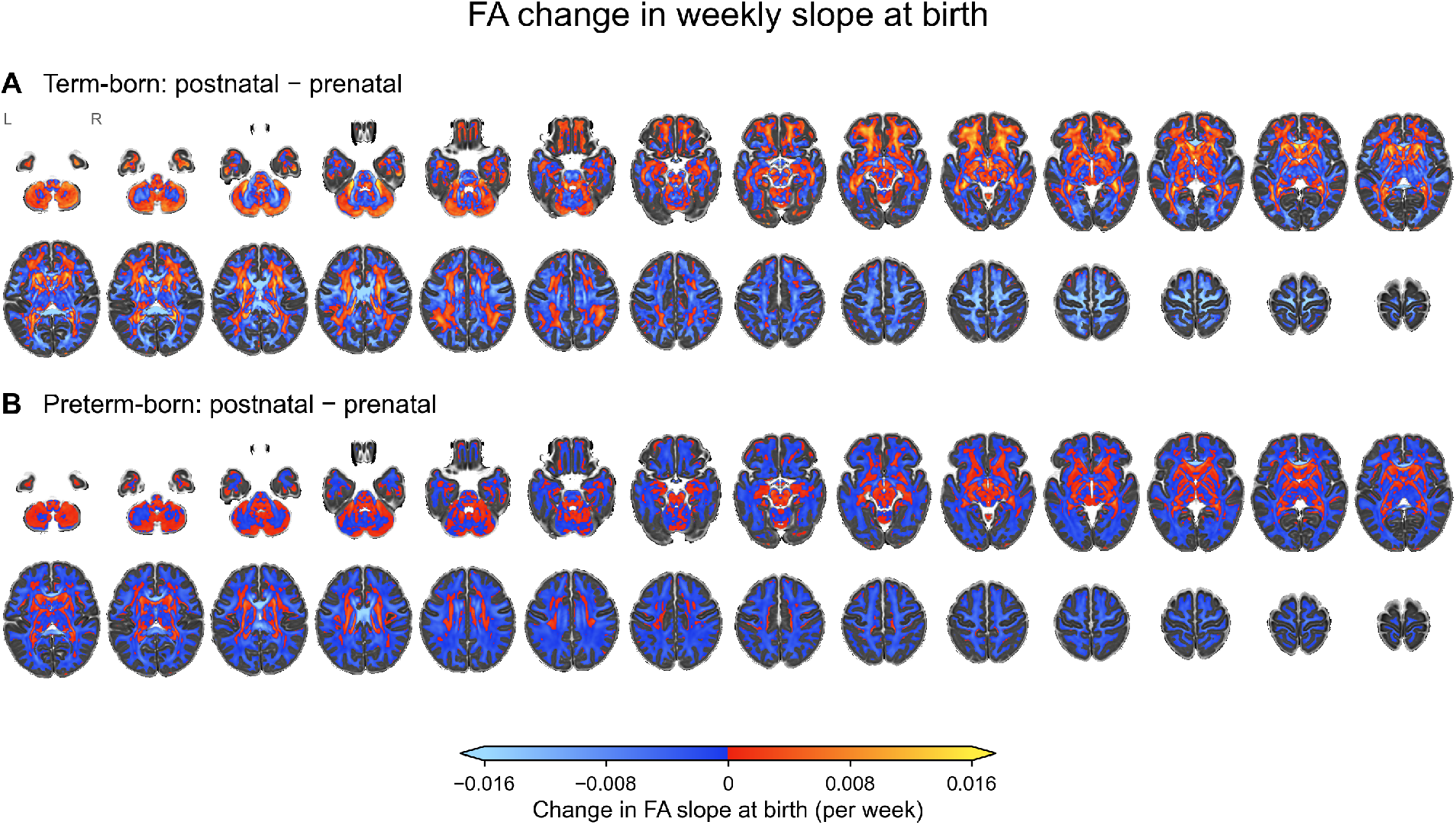
Change in the weekly FA slope at birth. Voxel-wise difference between the postnatal and prenatal slopes of FA for (A) term-born (*β*_1,term_ − *β*_0_) and (B) preterm-born infants (*β*_1,preterm_ − *β*_0_).

**Figure S6.**
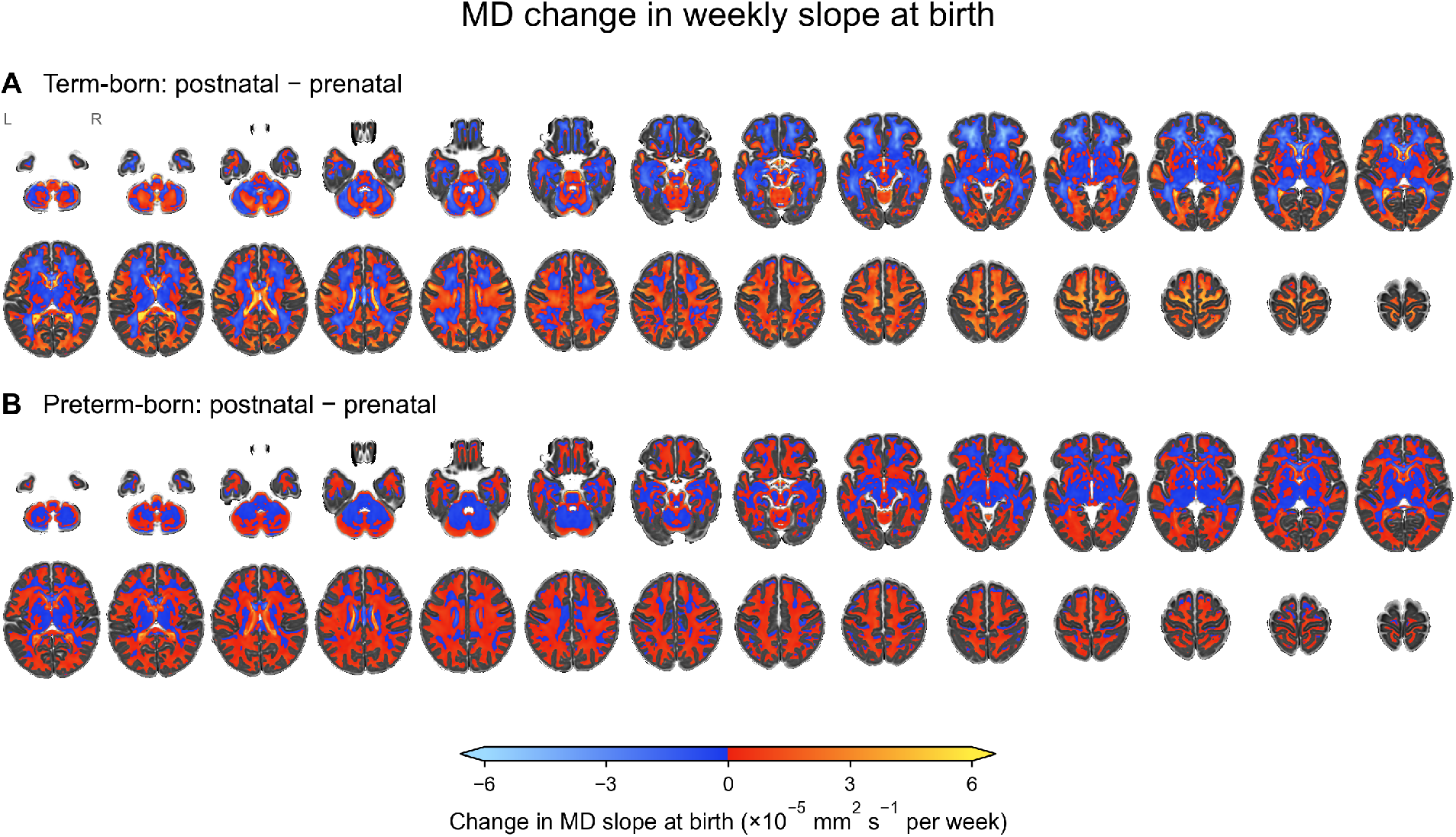
Change in the weekly MD slope at birth. Voxel-wise difference between the postnatal and prenatal slopes of MD for (A) term-born (*β*_1,term_ − *β*_0_) and (B) preterm-born infants (*β*_1,preterm_ − *β*_0_).

**Figure S7.**
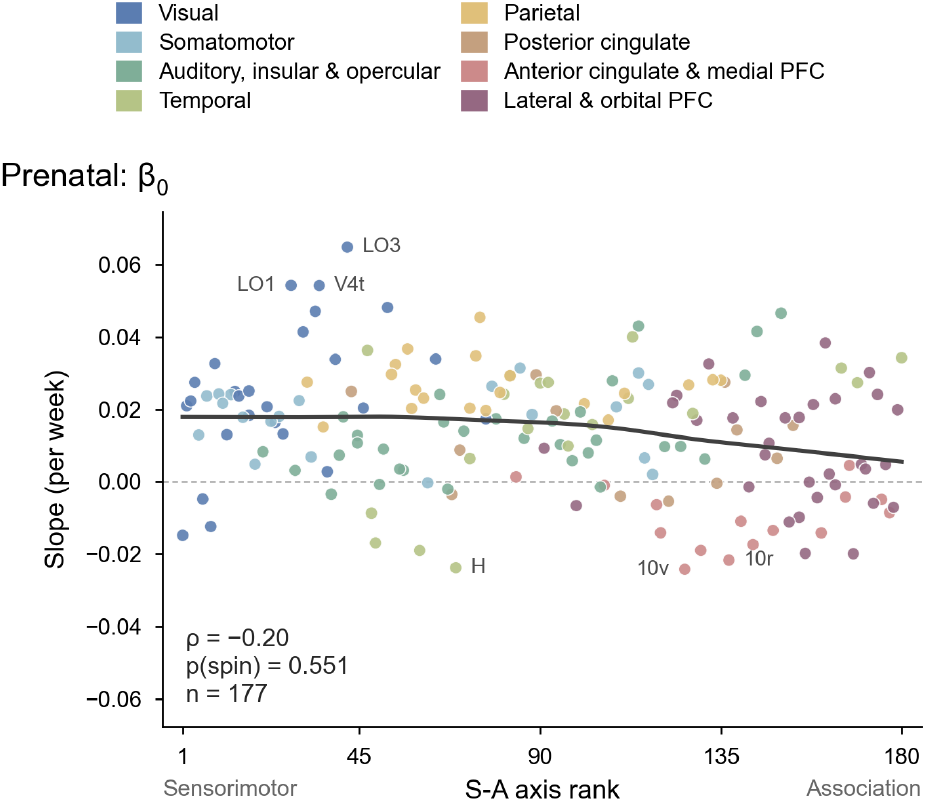
Prenatal slope of cortical innervation entropy along the sensorimotor to association axis. Each point is one area of the HCP multimodal parcellation (left and right hemispheres averaged; *n* = 177), plotted against its sensorimotor to association axis rank (Sydnor et al. 2021) and coloured by cortical division. The prenatal slope (*β*_0_) was not associated with the axis (Spearman *ρ* = −0.20, *p*_spin_ = 0.55; 5,000 rotations).

**Figure S8.**
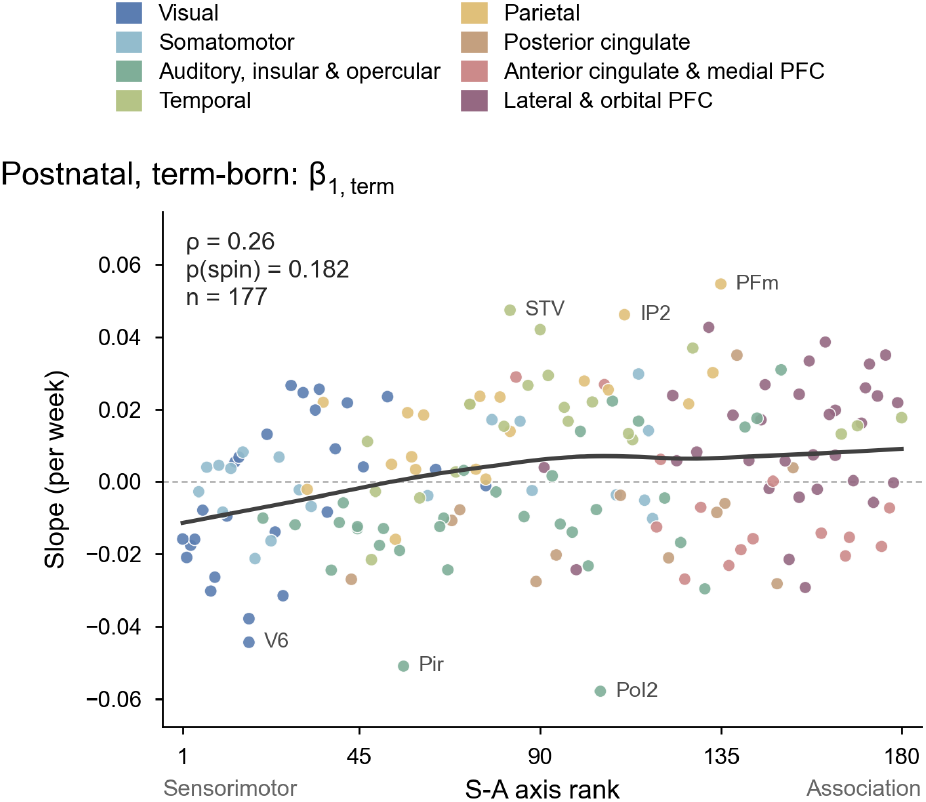
Postnatal slope in term infants along the sensorimotor to association axis. Equivalent plot of Figure S7, for the postnatal slope in term-born infants (*β*_1,term_). The slope was not significantly associated with the axis (*ρ* = 0.26, *p*_spin_ = 0.18).

**Figure S9.**
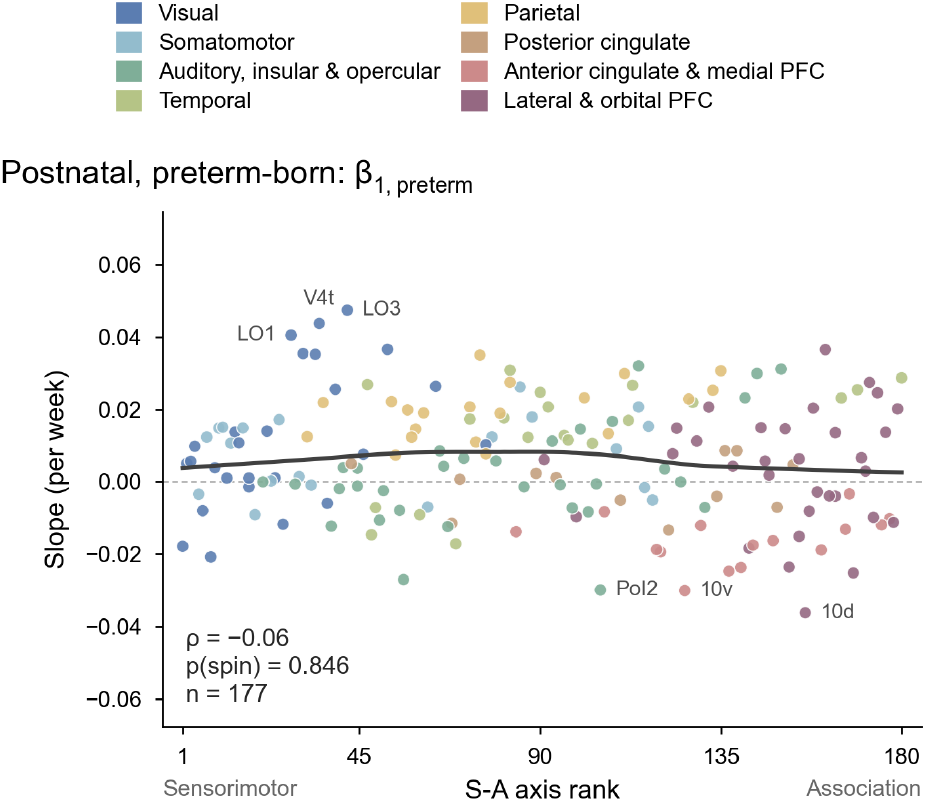
Postnatal slope in preterm-born infants along the sensorimotor to association axis. Equivalent plot of Figure S7 for the postnatal slope in preterm infants (*β*_1,preterm_). The slope was not associated with the axis (*ρ* = −0.06, *p*_spin_ = 0.85).

**Figure S10.**
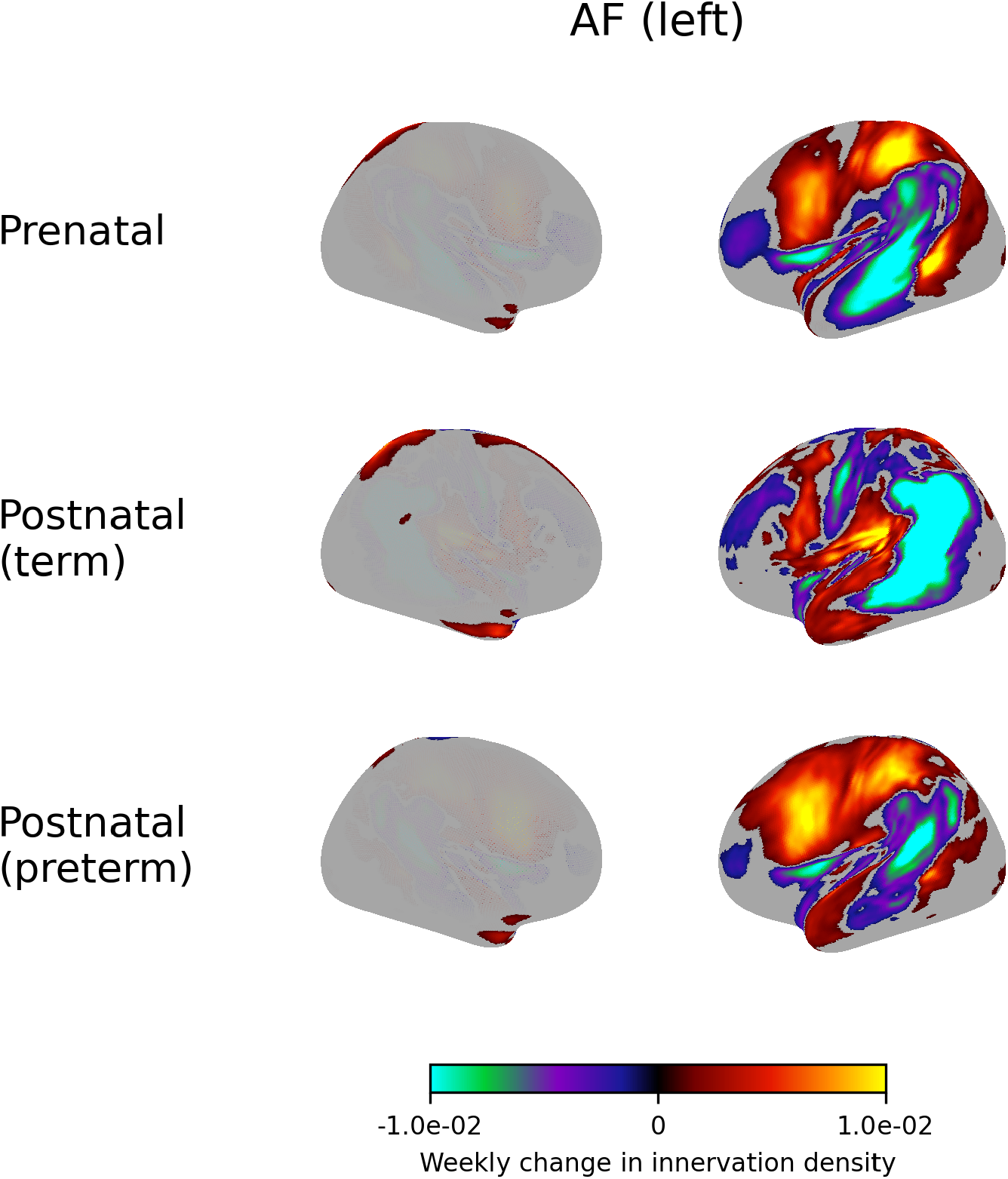
Weekly change in innervation density of the left arcuate fasciculus before birth (prenatal) and after term and preterm birth (postnatal). Each tract’s cortical map is normalised to sum to one, so positive values indicate an increasing share of the tract’s terminations at a vertex and negative values a decreasing share. Increases in some regions are therefore balanced by decreases elsewhere, and the maps describe the redistribution of terminations across the cortex rather than absolute gains or losses in connectivity. Values with magnitude below 10^−3^ are not shown.

**Figure S11.**
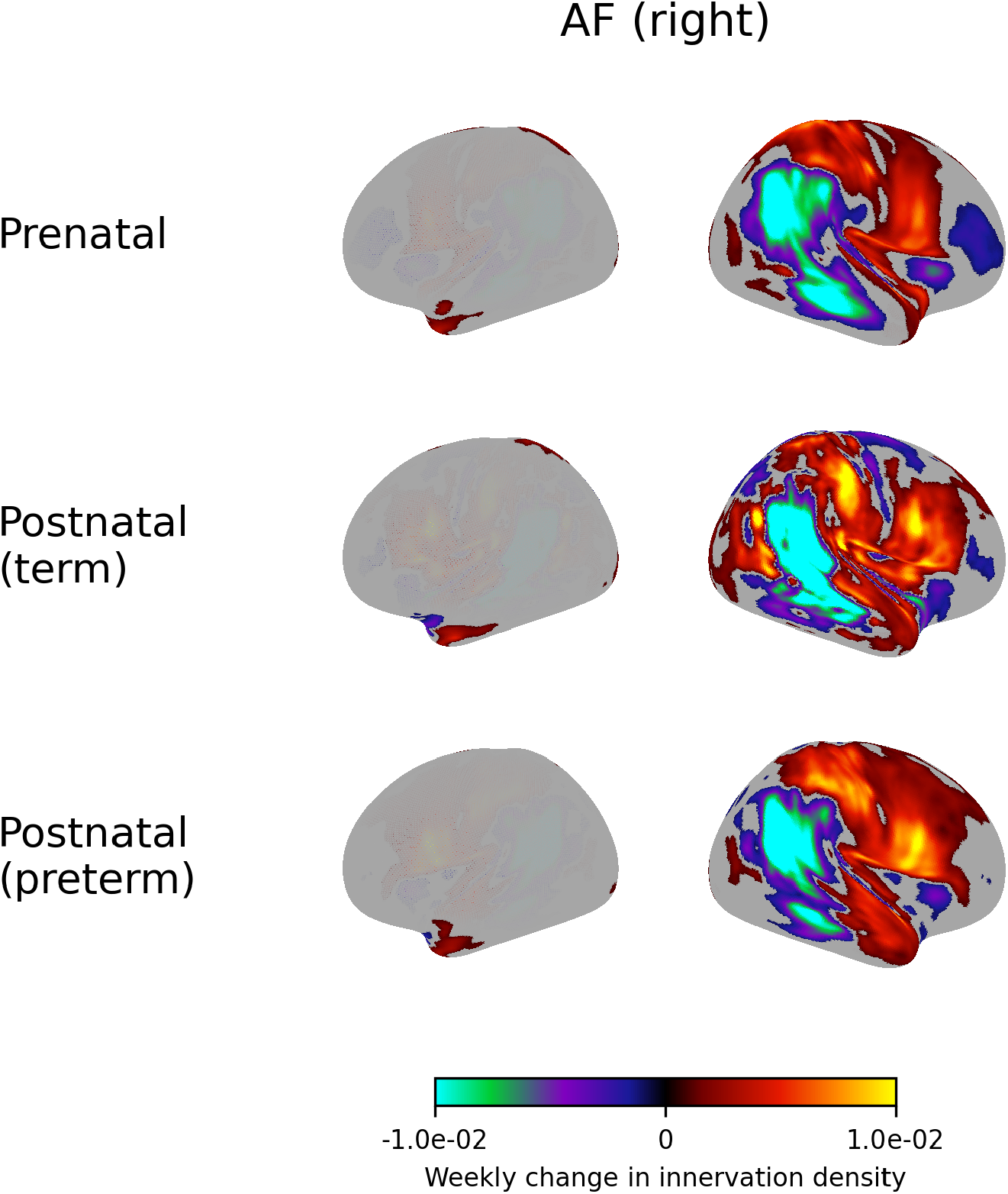
Weekly change in innervation density of the right arcuate fasciculus.

**Figure S12.**
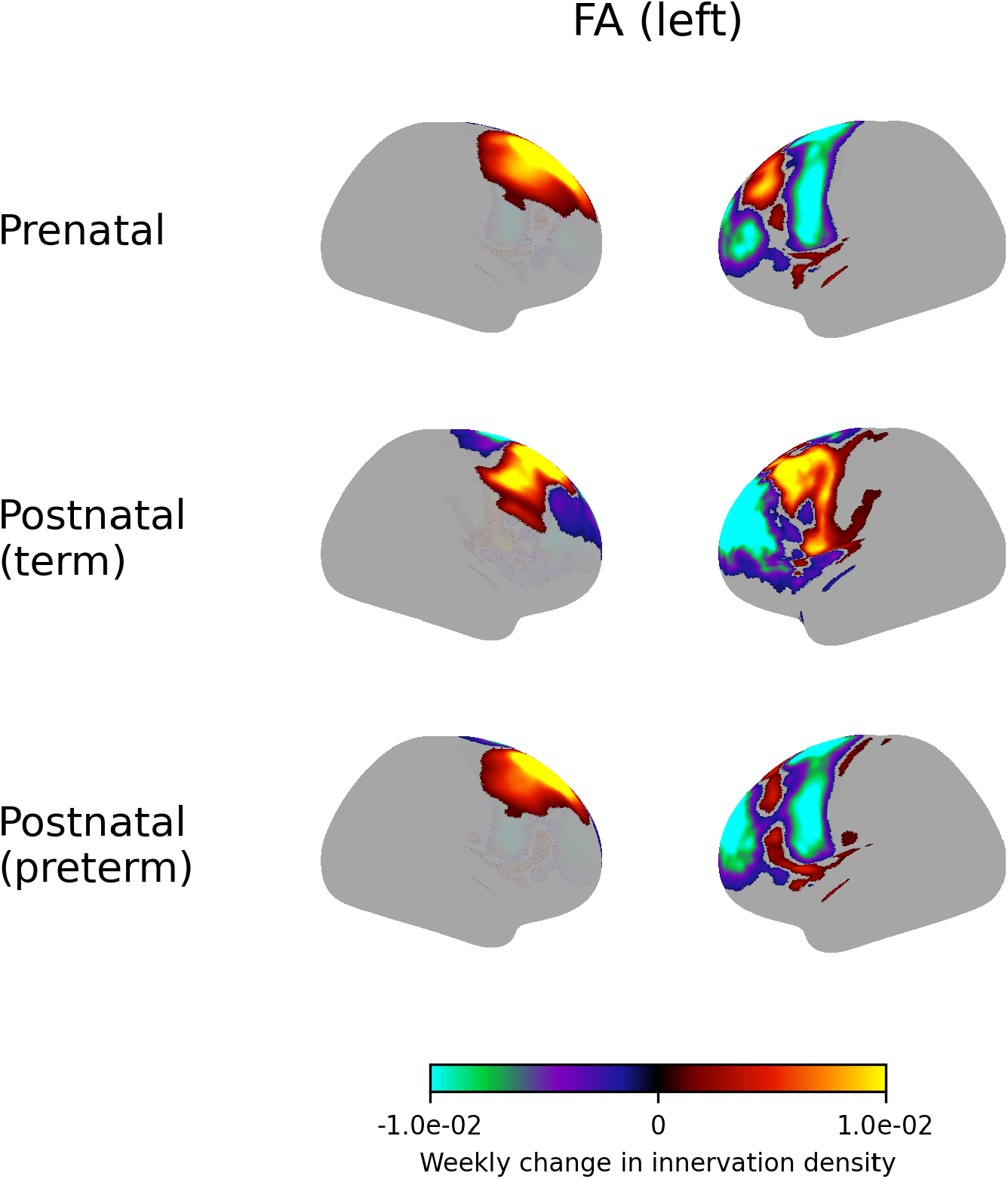
Weekly change in innervation density of the left frontal aslant tract.

**Figure S13.**
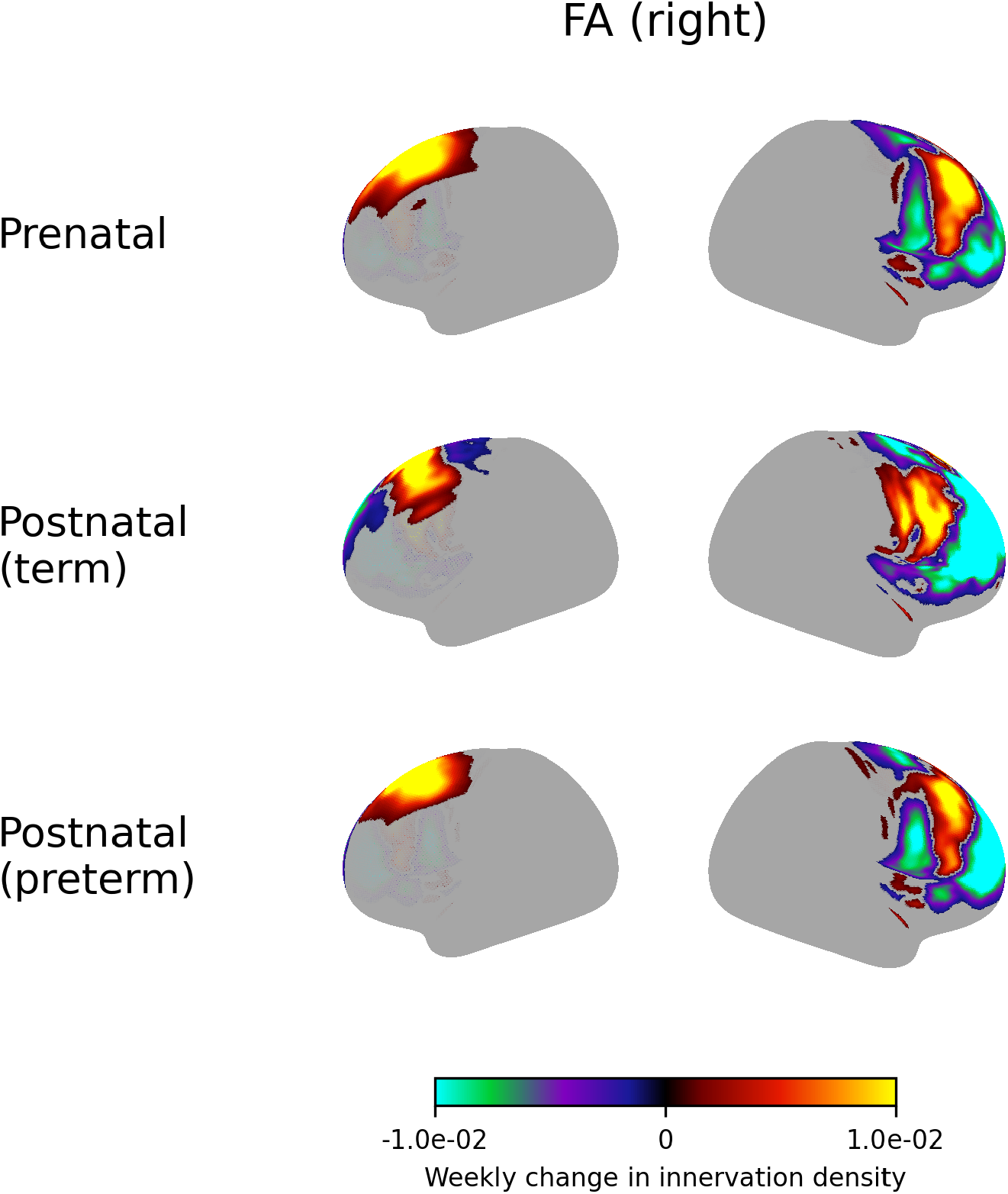
Weekly change in innervation density of the right frontal aslant tract.

**Figure S14.**
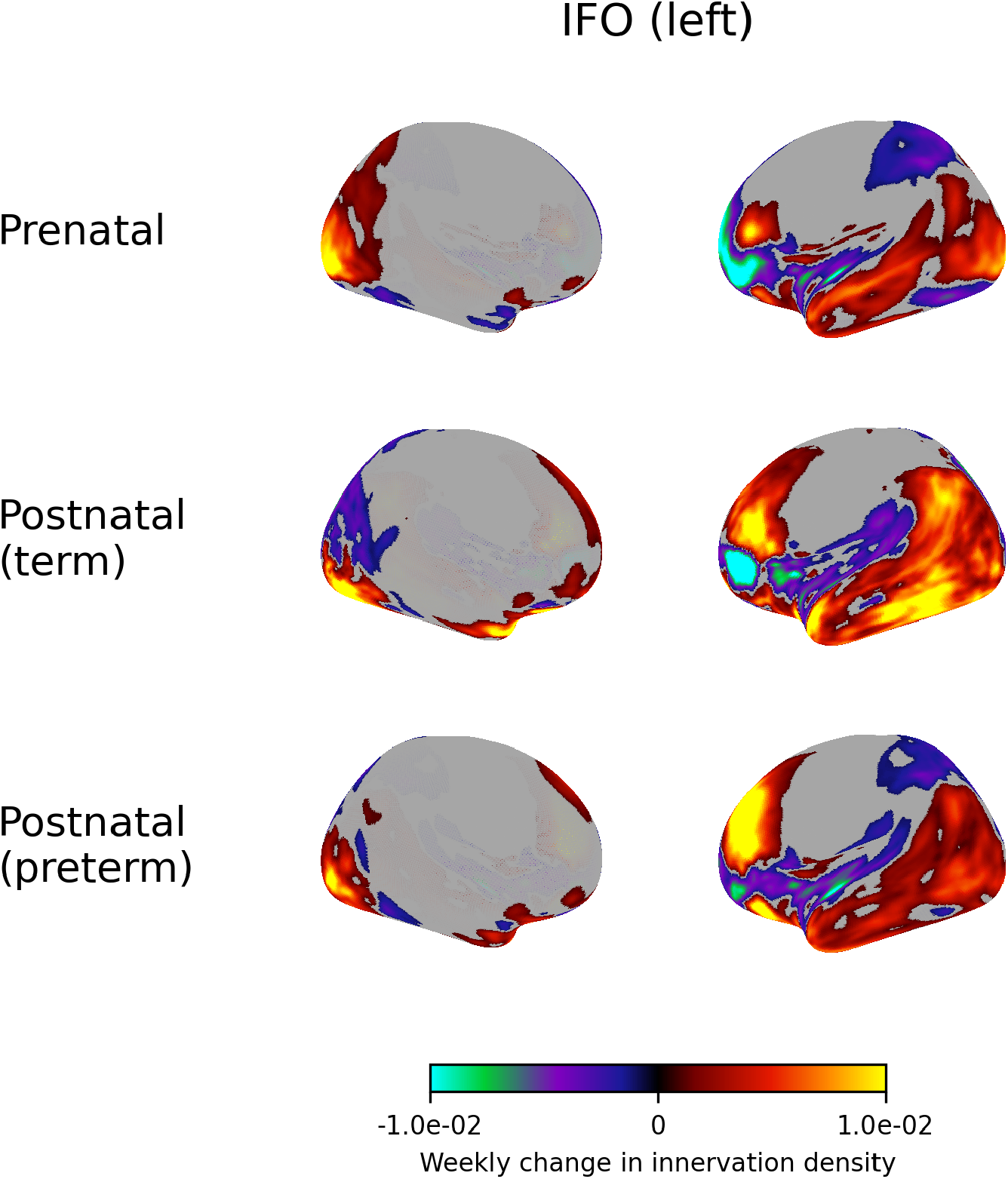
Weekly change in innervation density of the left inferior frontooccipital fasciculus.

**Figure S15.**
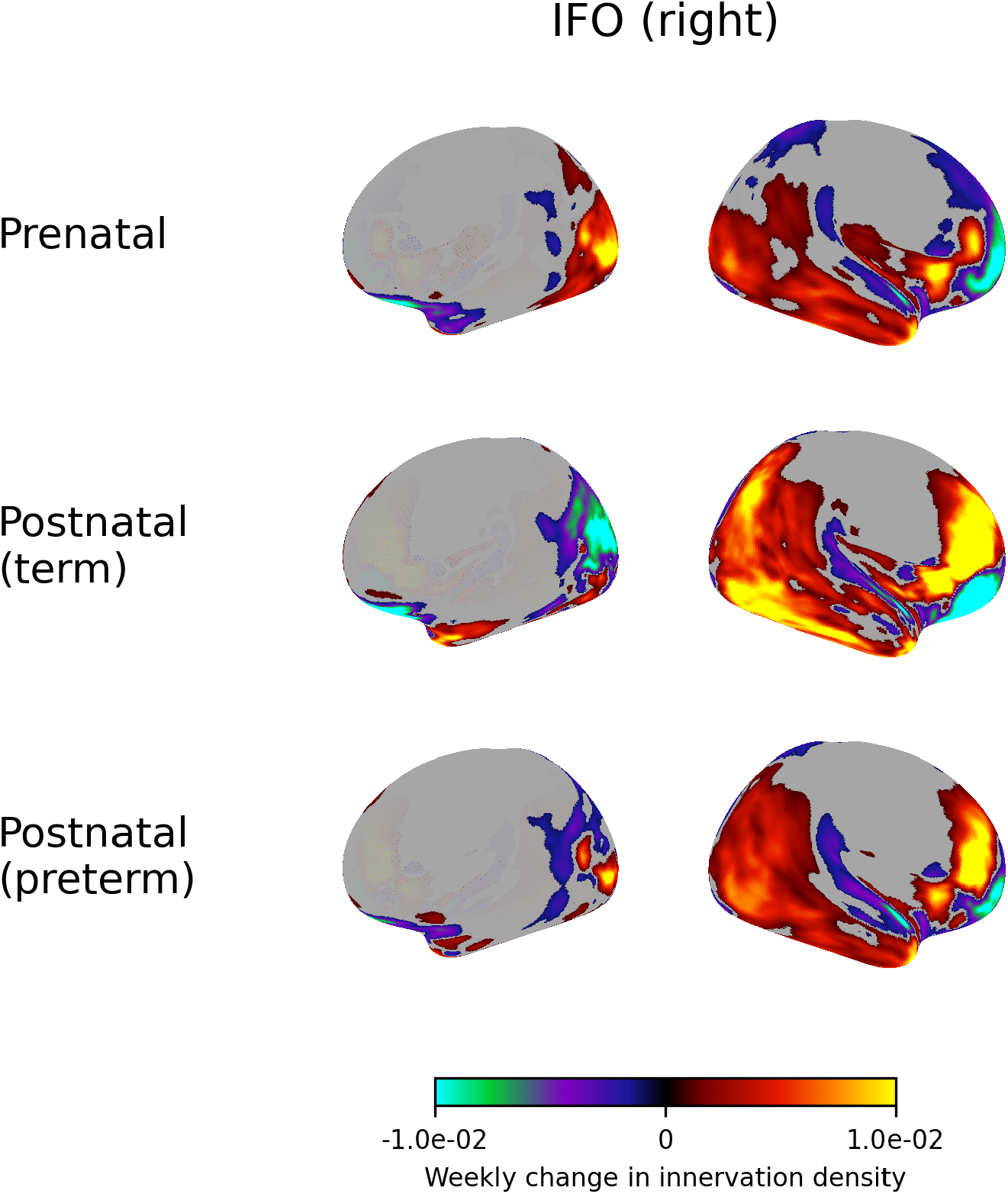
Weekly change in innervation density of the right inferior frontooccipital fasciculus.

**Figure S16.**
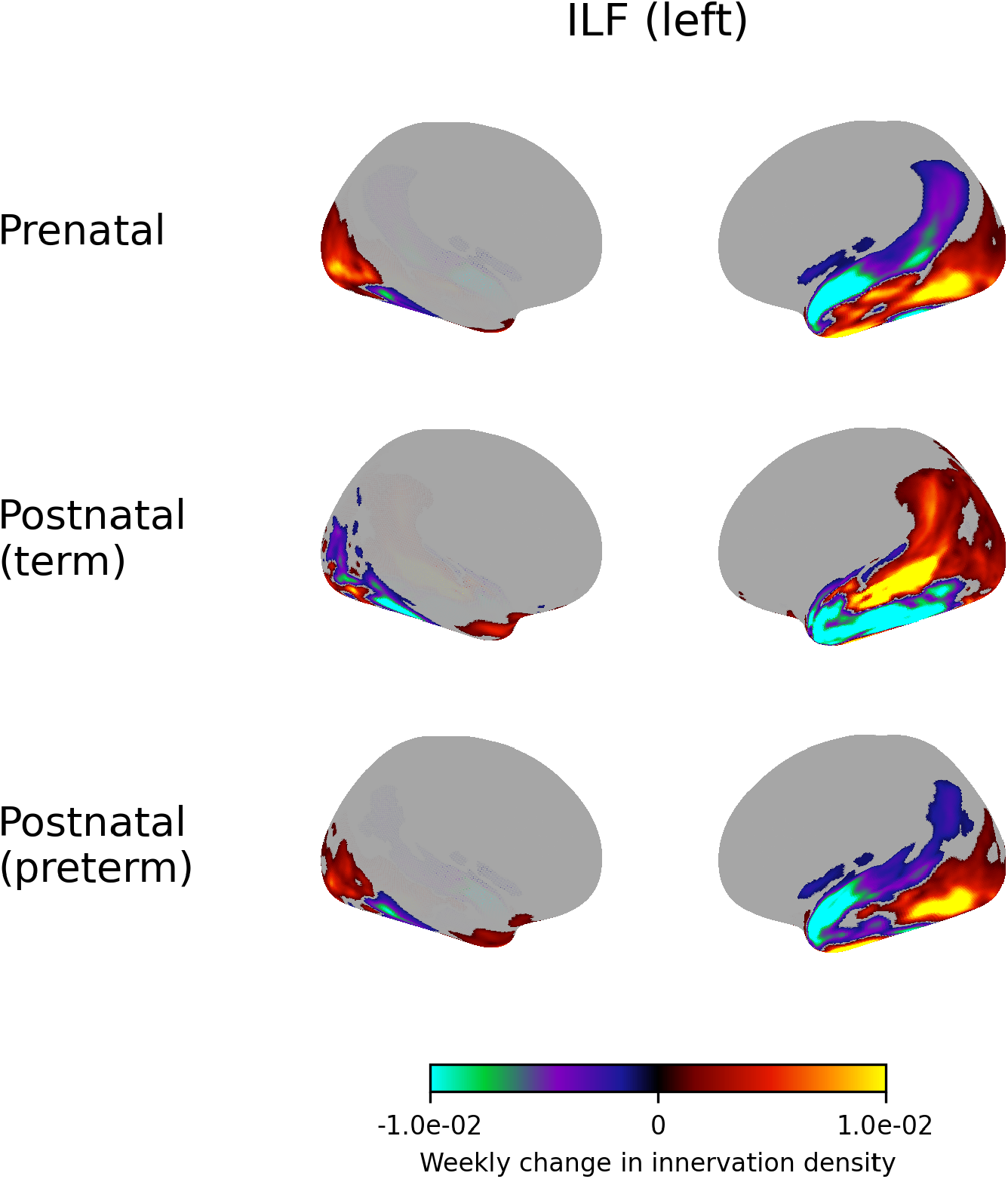
Weekly change in innervation density of the left inferior longitudinal fasciculus.

**Figure S17.**
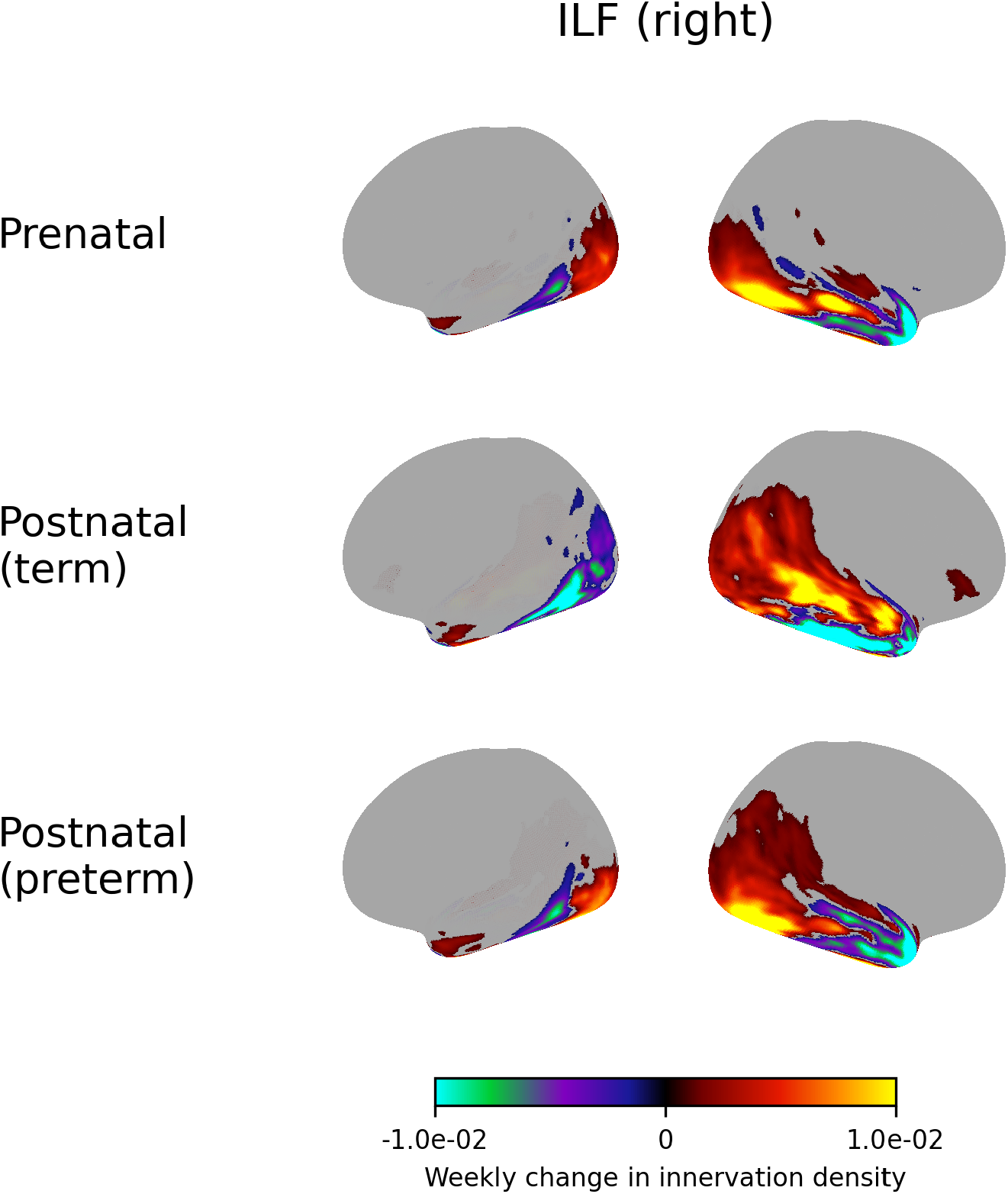
Weekly change in innervation density of the right inferior longitudinal fasciculus.

**Figure S18.**
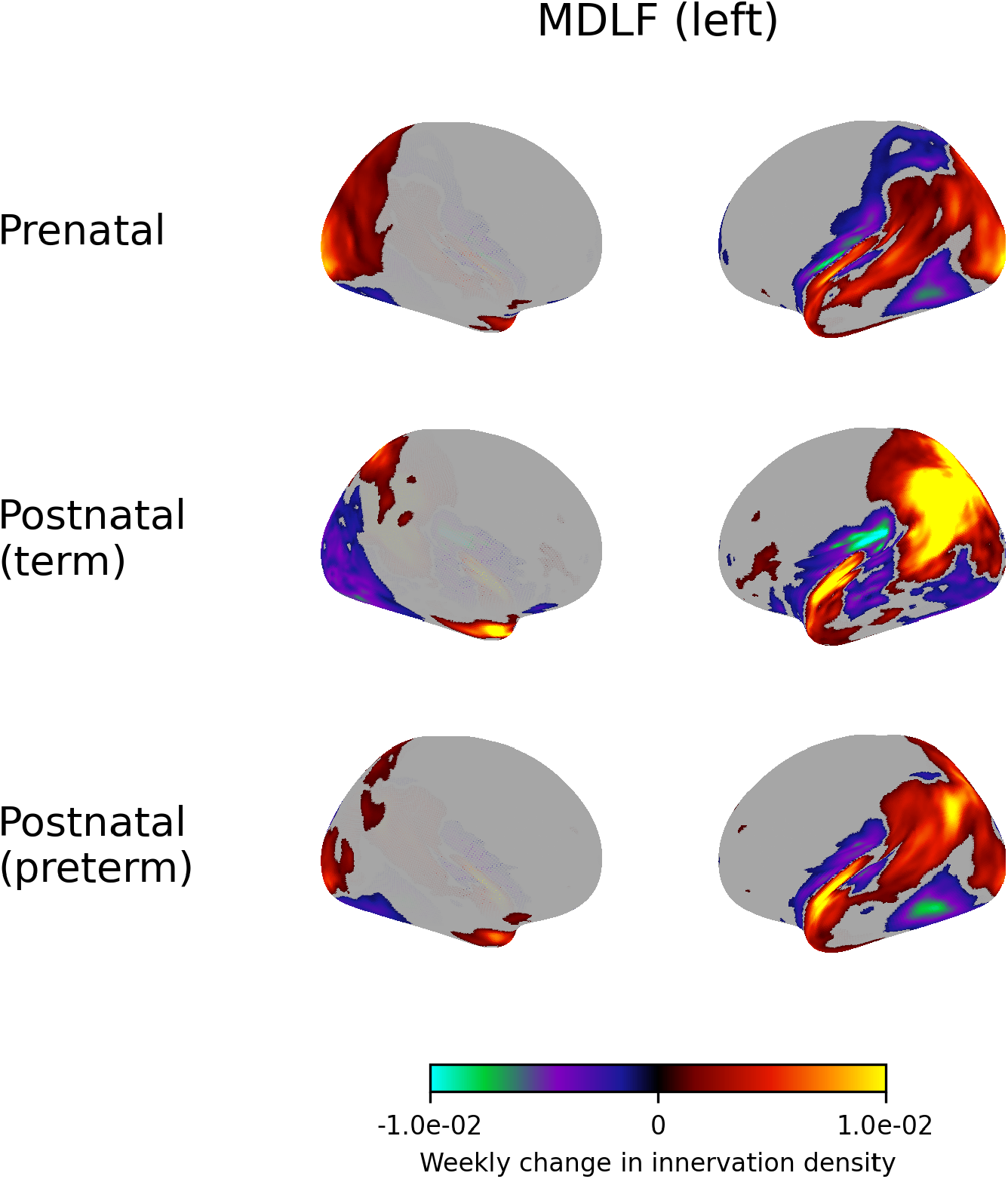
Weekly change in innervation density of the left middle longitudinal fasciculus.

**Figure S19.**
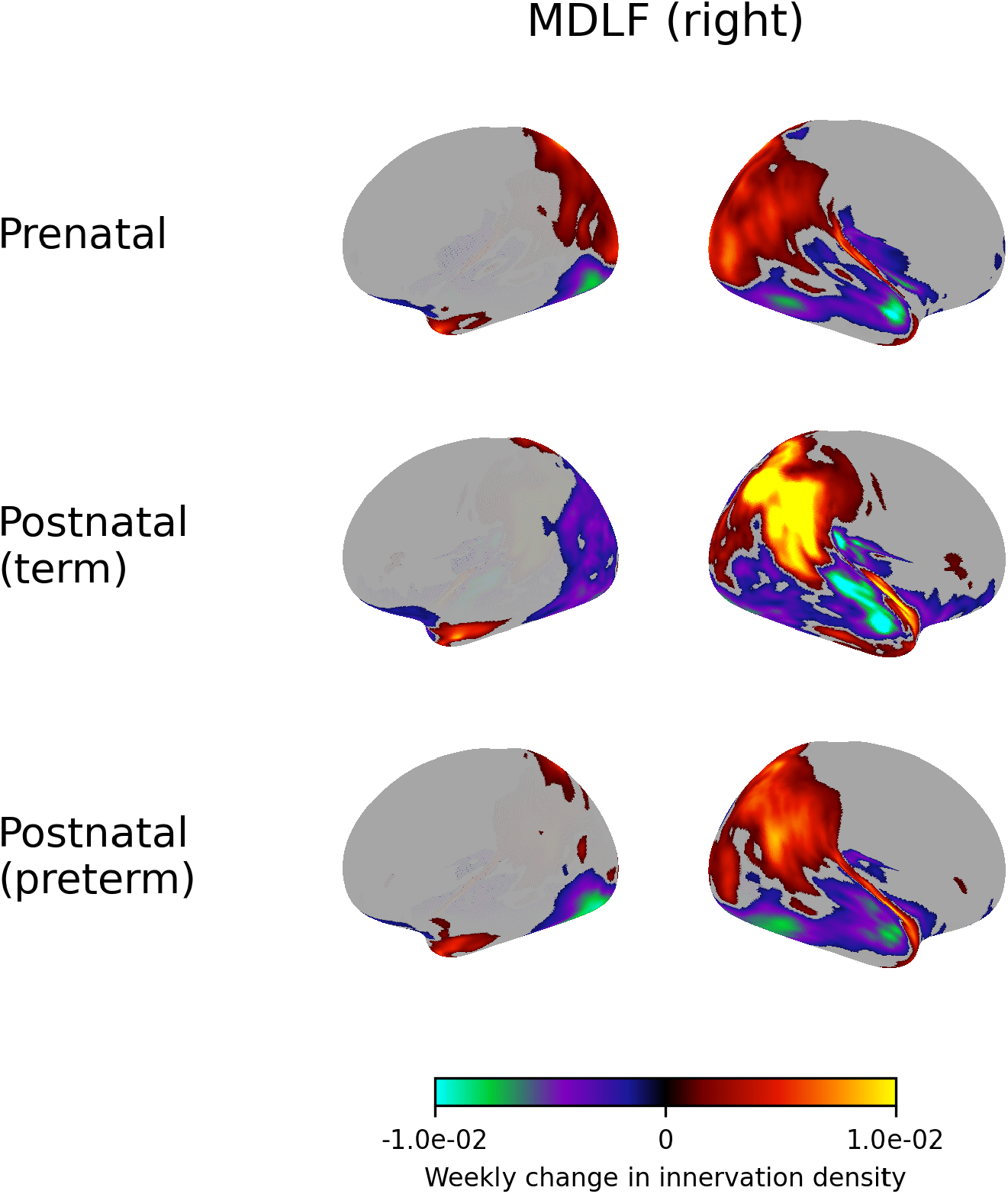
Weekly change in innervation density of the right middle longitudinal fasciculus.

**Figure S20.**
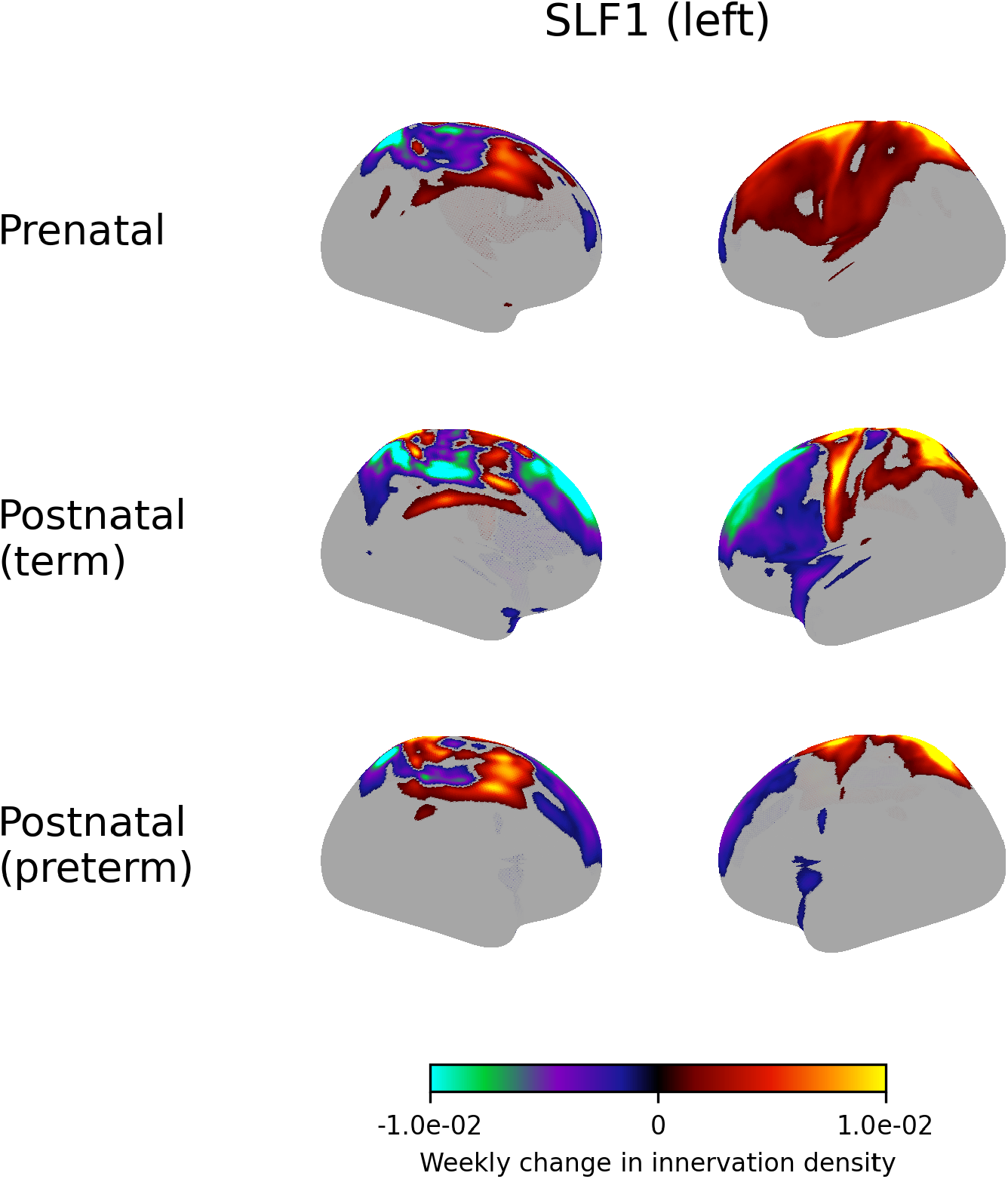
Weekly change in innervation density of the left superior longitudinal fasciculus I.

**Figure S21.**
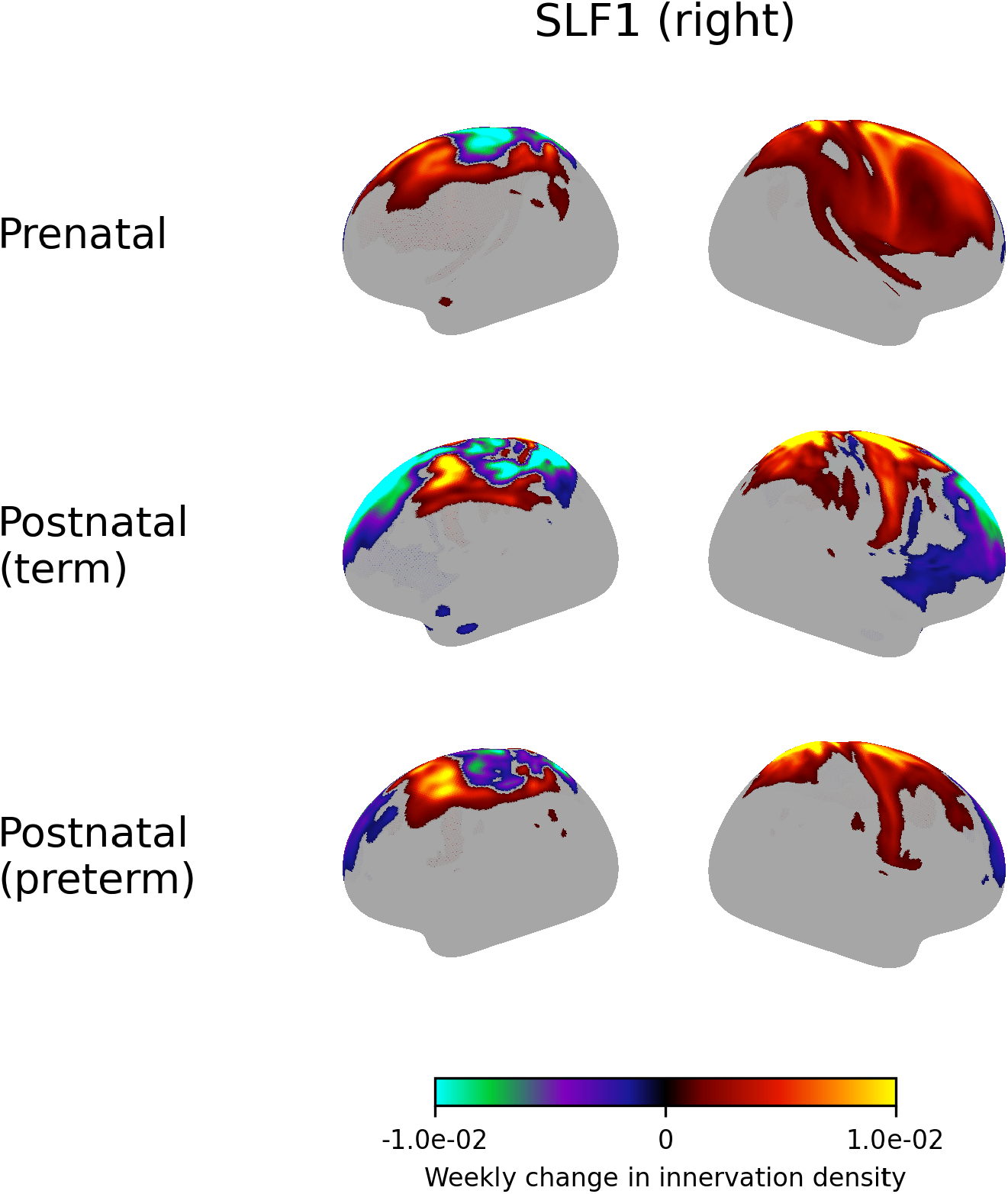
Weekly change in innervation density of the right superior longitudinal fasciculus I.

**Figure S22.**
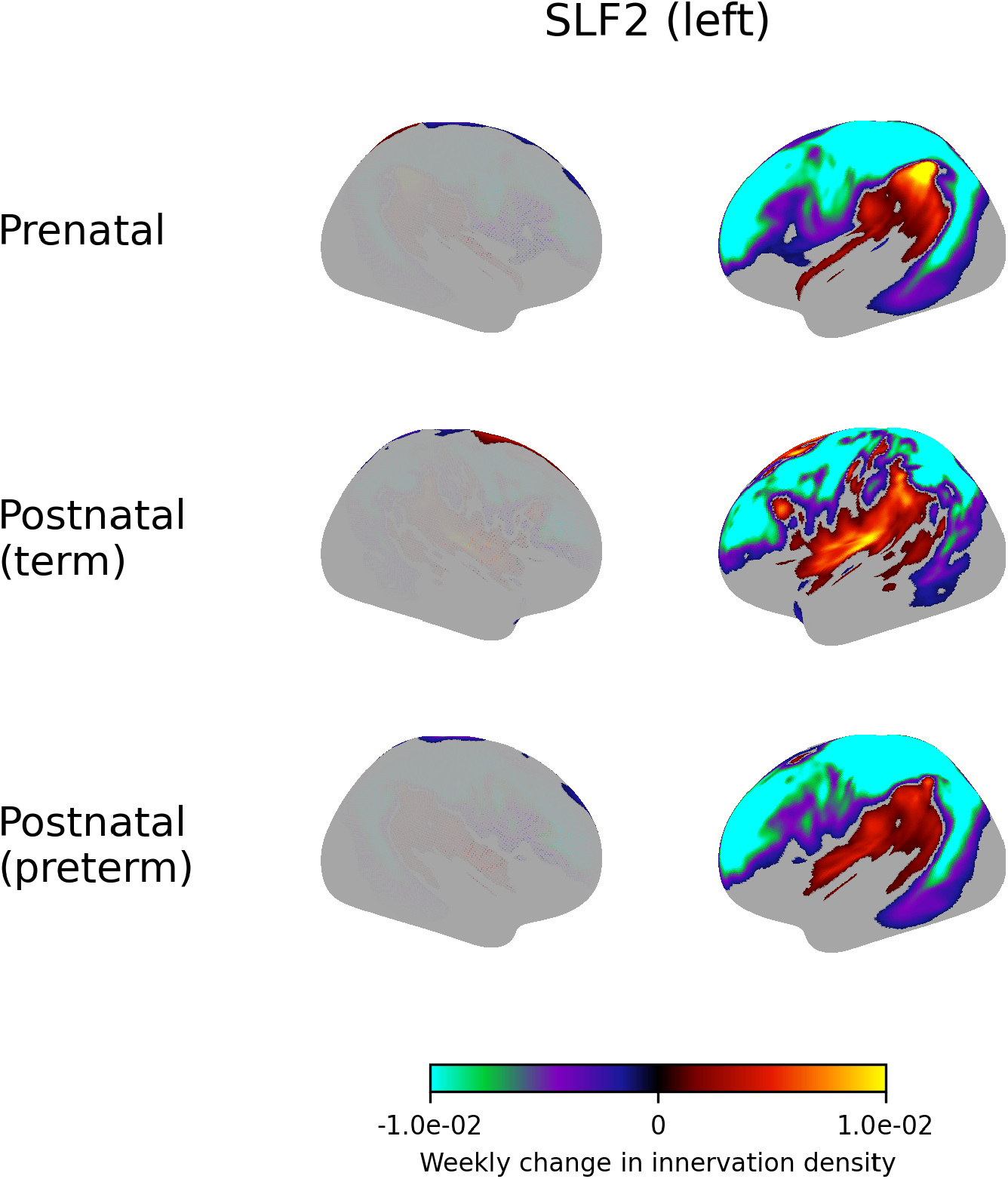
Weekly change in innervation density of the left superior longitudinal fasciculus II.

**Figure S23.**
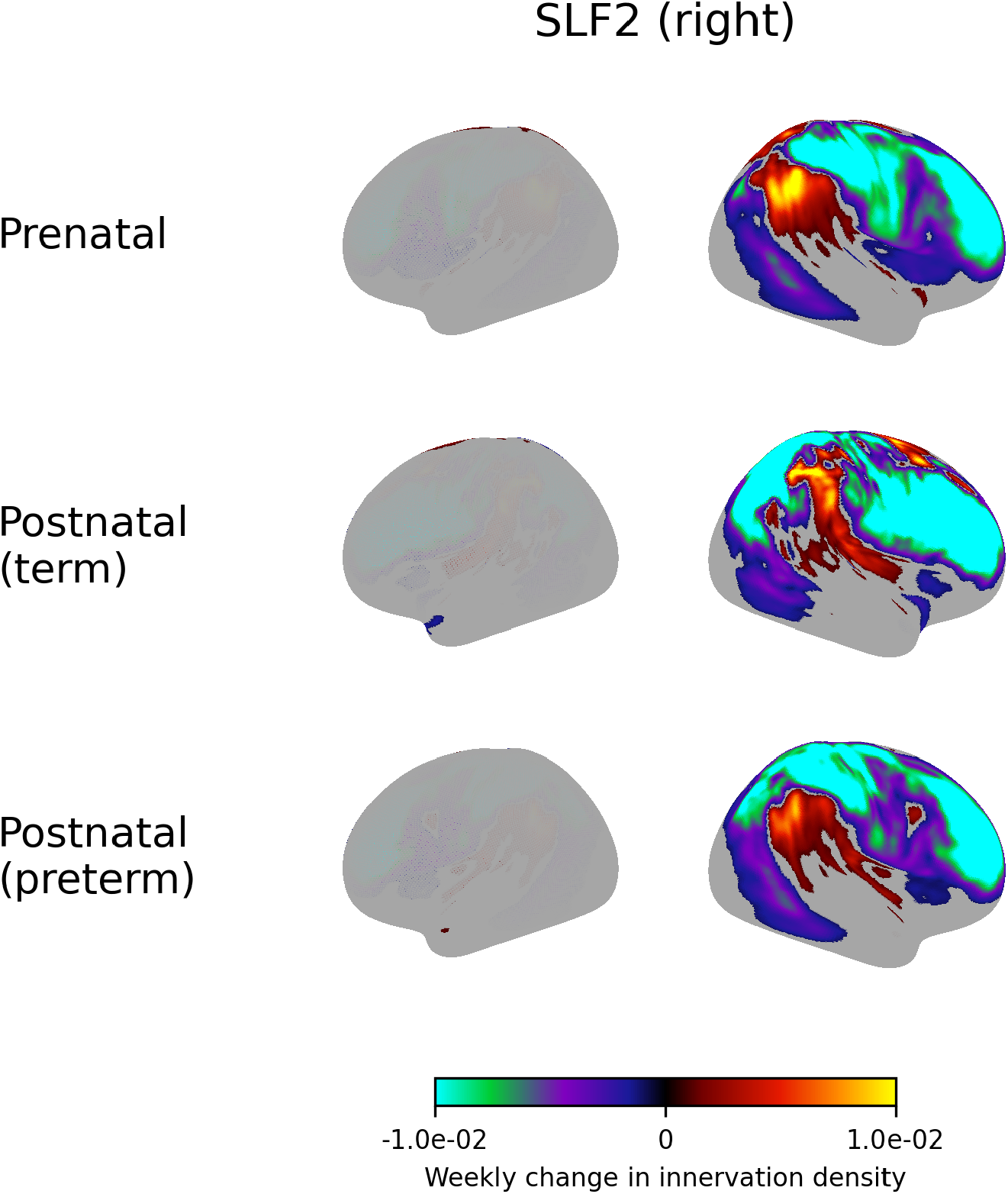
Weekly change in innervation density of the right superior longitudinal fasciculus II.

**Figure S24.**
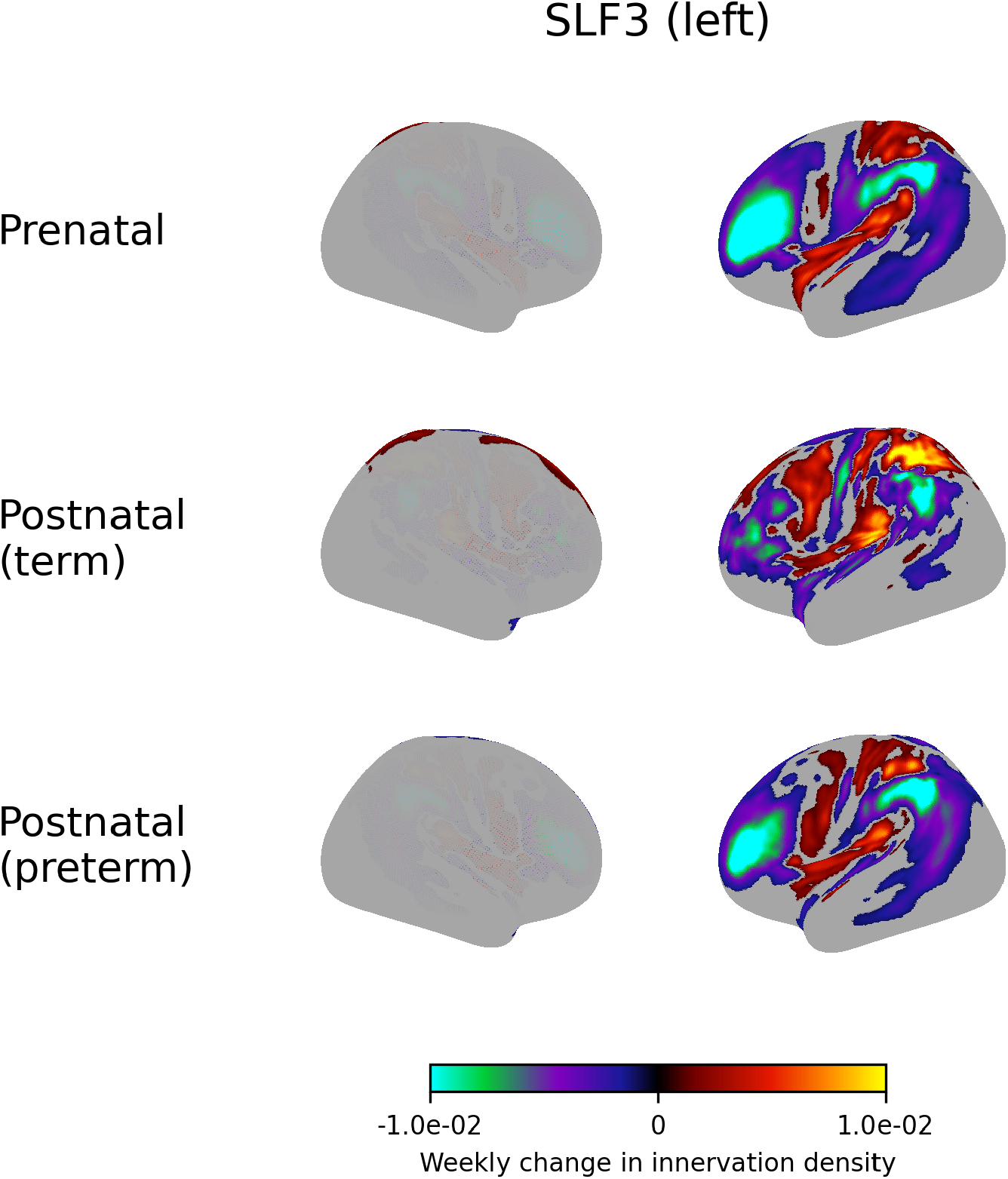
Weekly change in innervation density of the left superior longitudinal fasciculus III.

**Figure S25.**
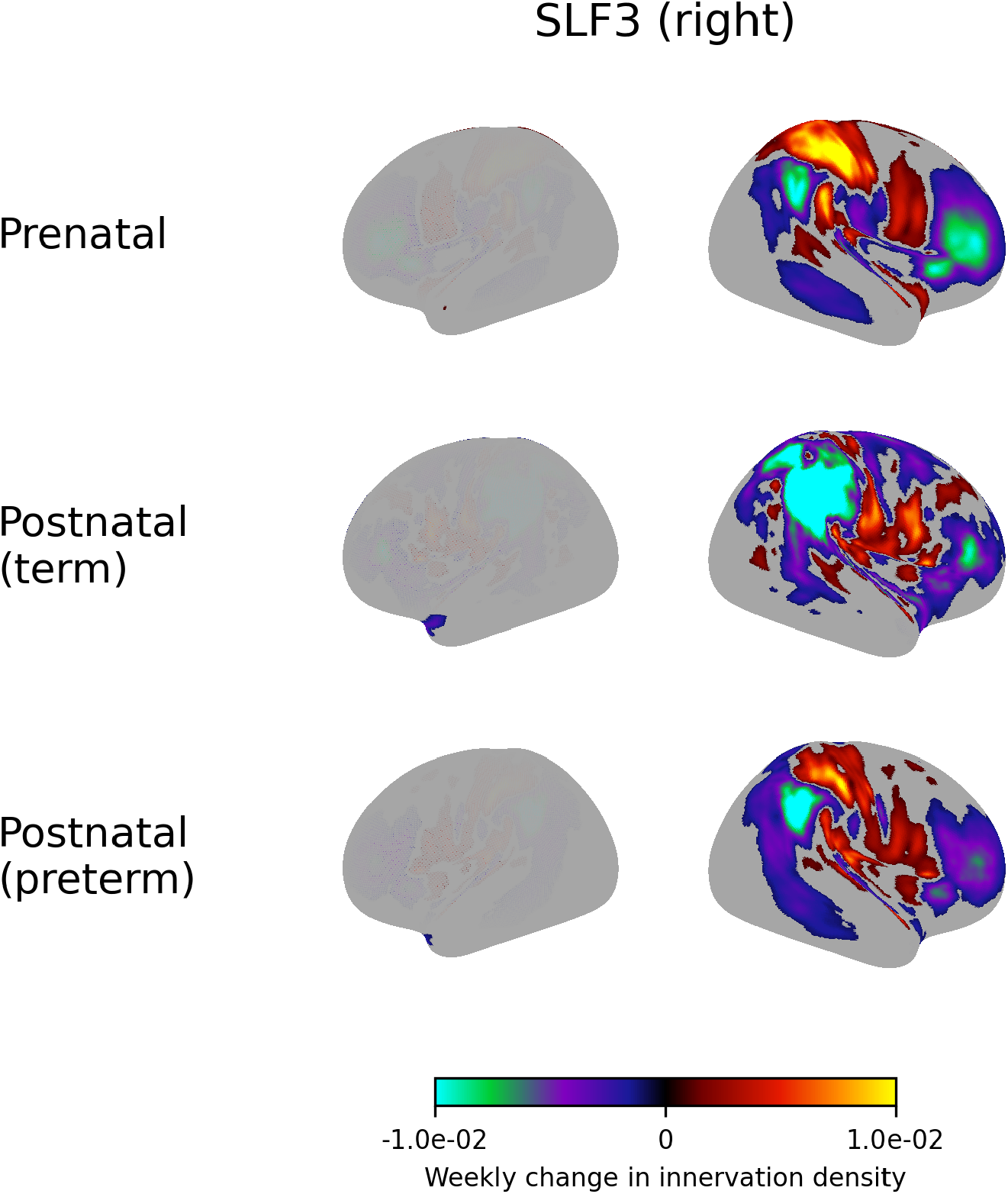
Weekly change in innervation density of the right superior longitudinal fasciculus III.

**Figure S26.**
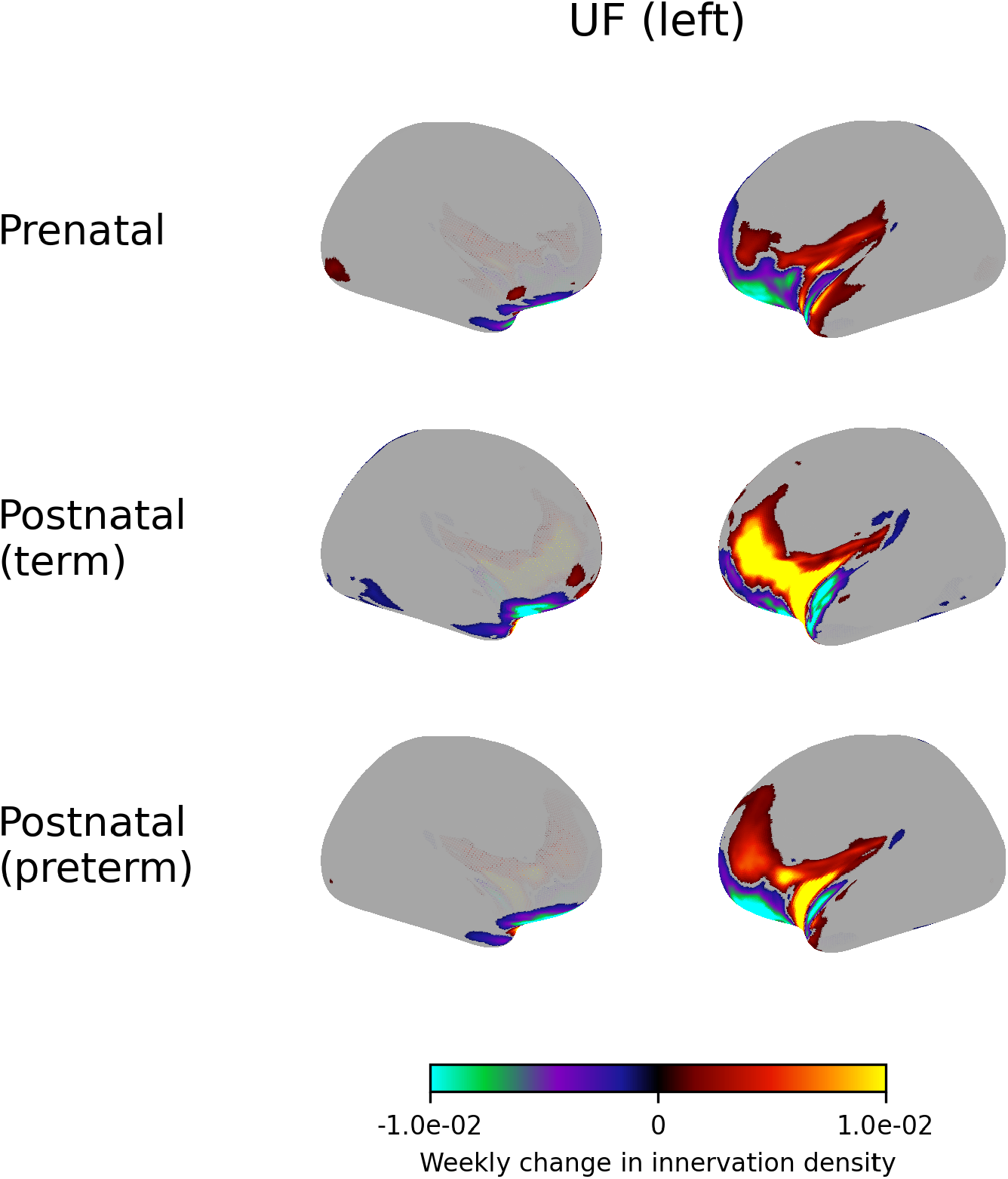
Weekly change in innervation density of the left uncinate fasciculus.

**Figure S27.**
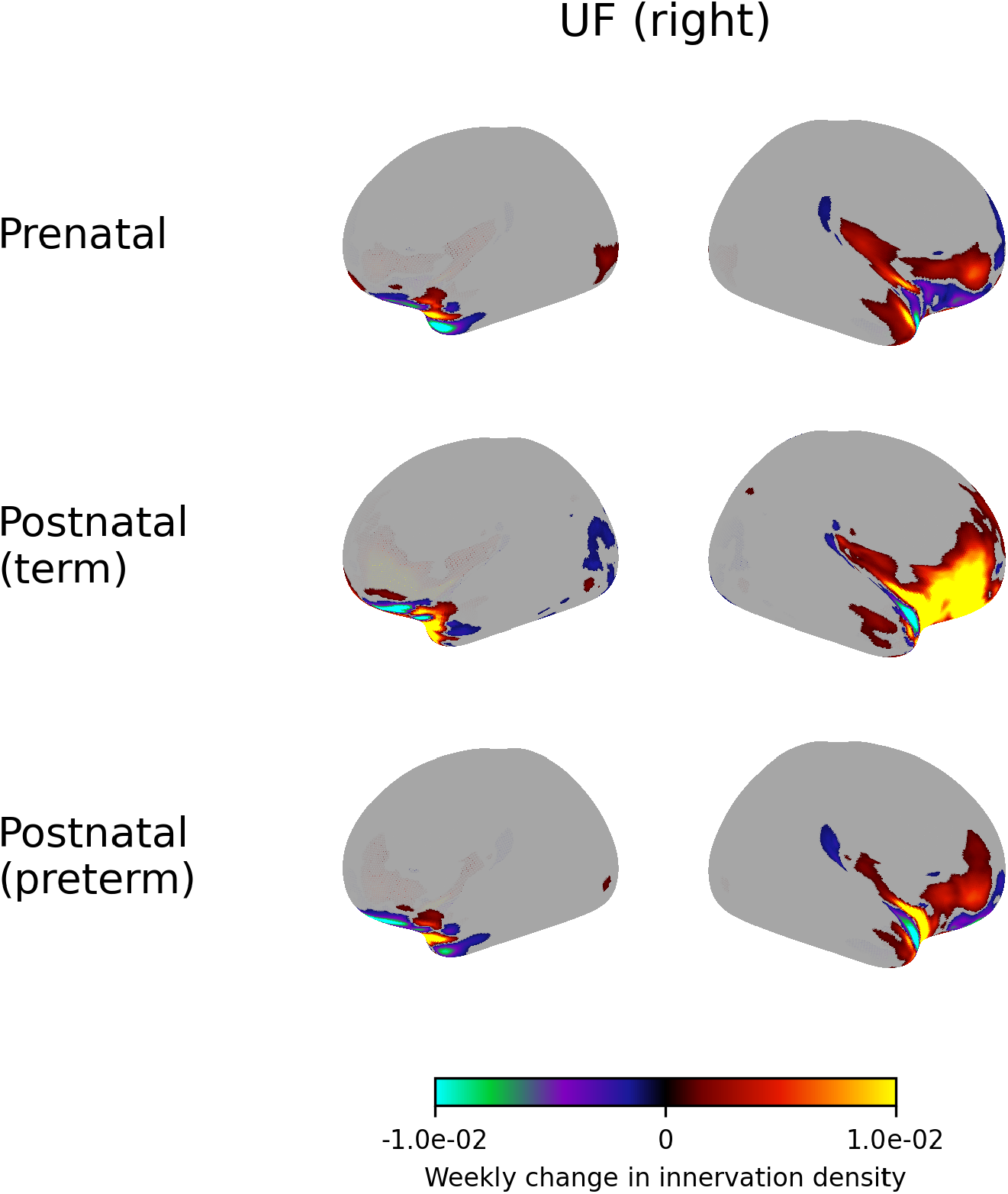
Weekly change in innervation density of the right uncinate fasciculus.

**Figure S28.**
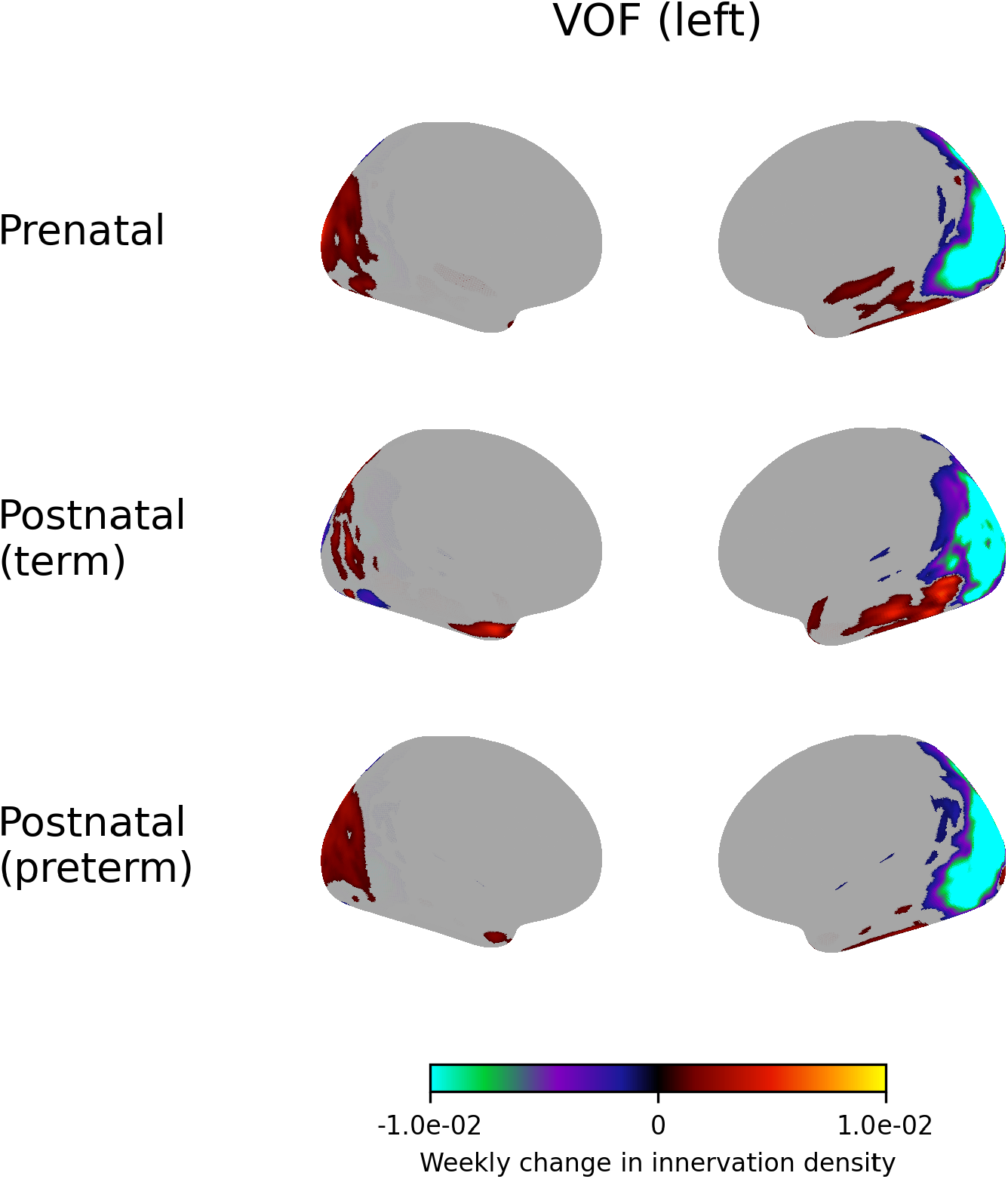
Weekly change in innervation density of the left vertical occipital fasciculus.

**Figure S29.**
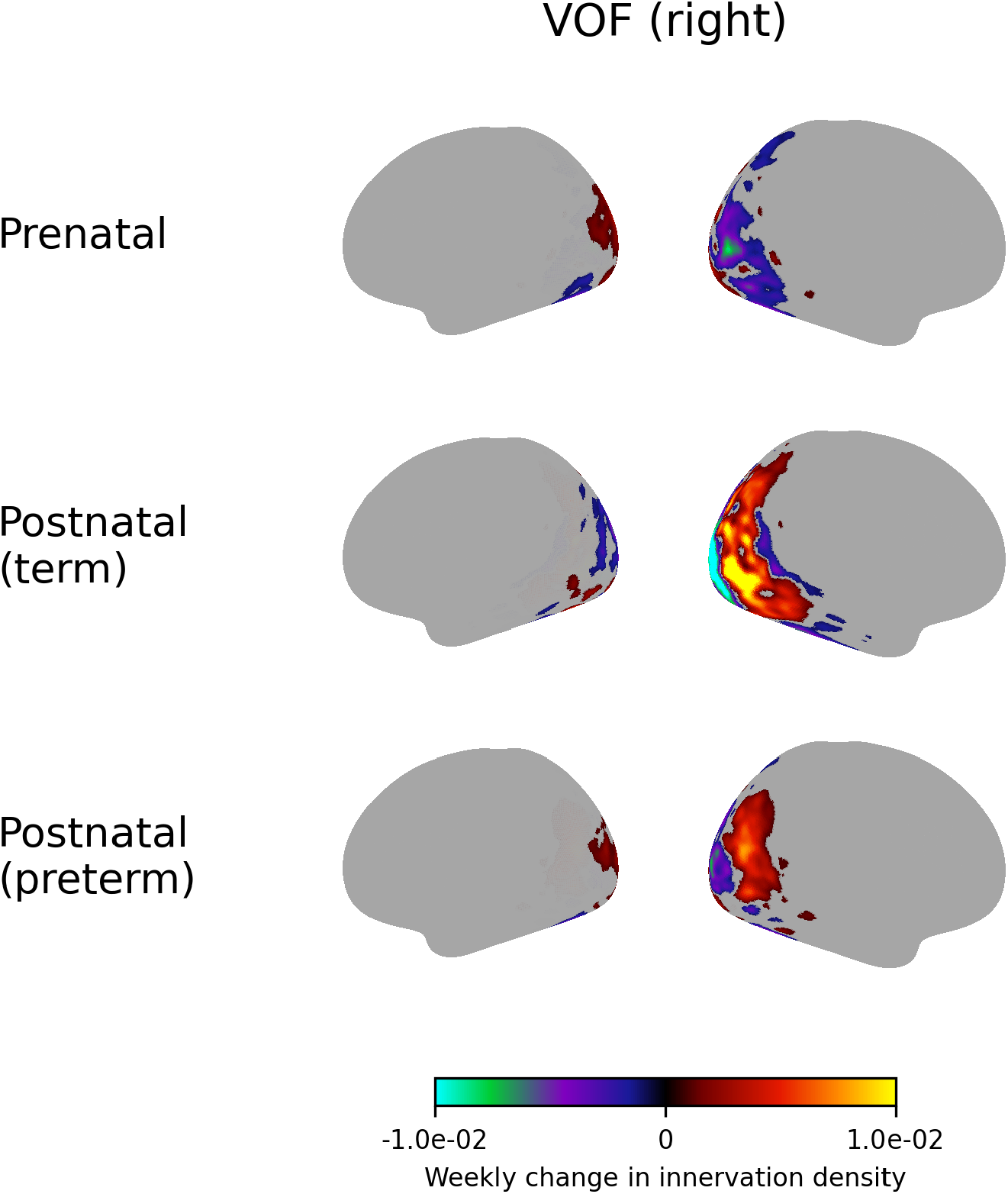
Weekly change in innervation density of the right vertical occipital fasciculus.

**Figure S30.**
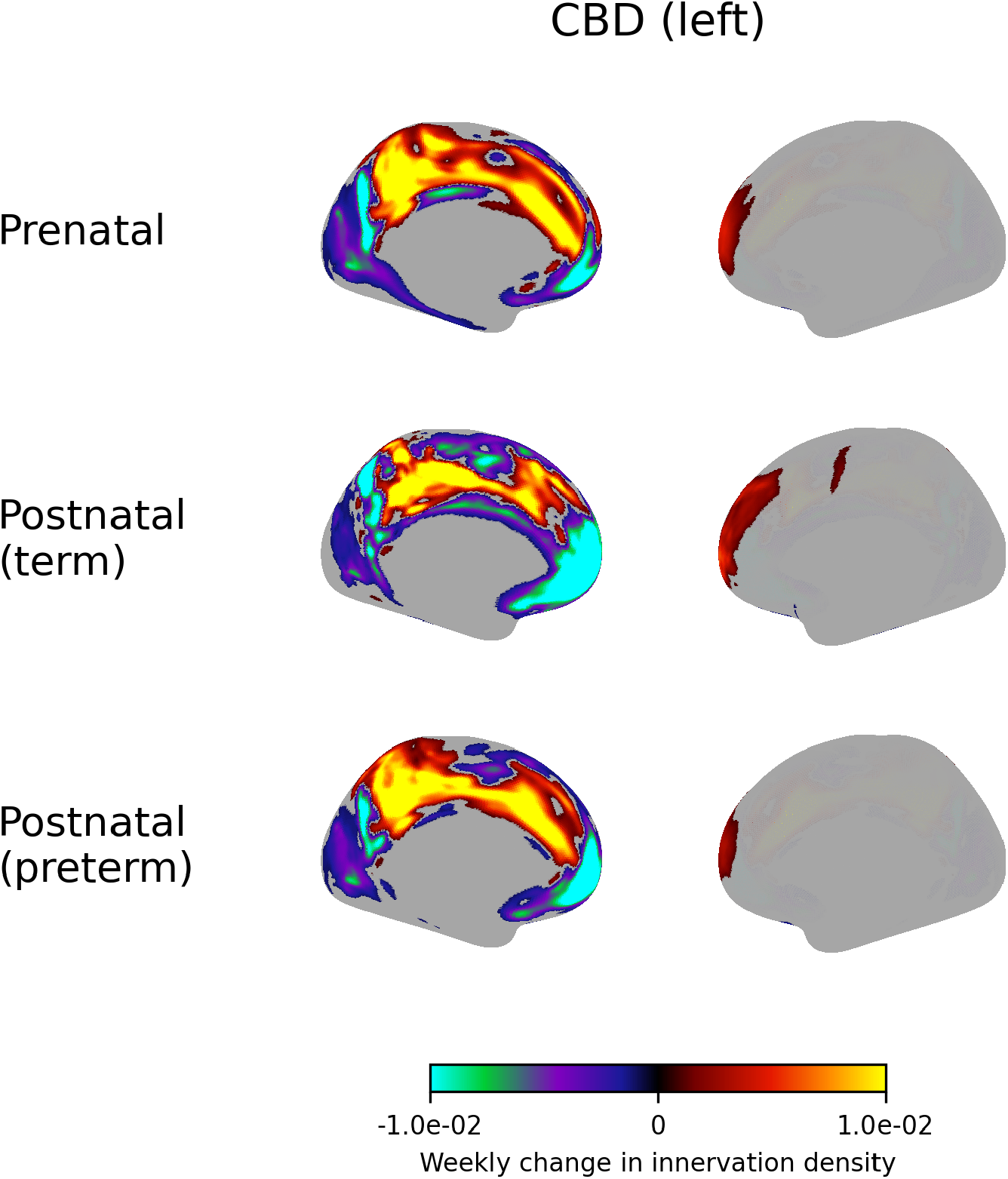
Weekly change in innervation density of the left dorsal cingulum bundle.

**Figure S31.**
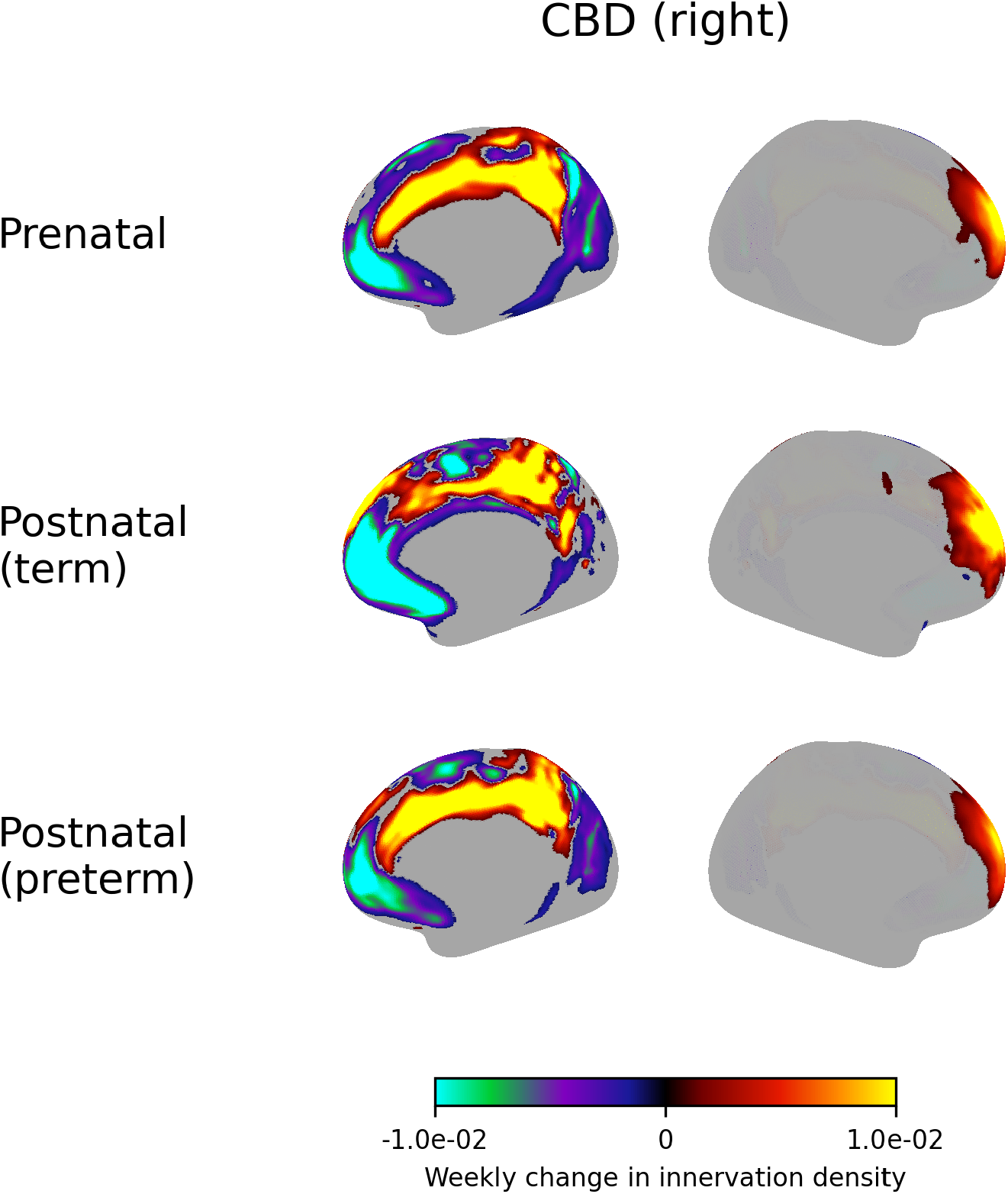
Weekly change in innervation density of the right dorsal cingulum bundle.

**Figure S32.**
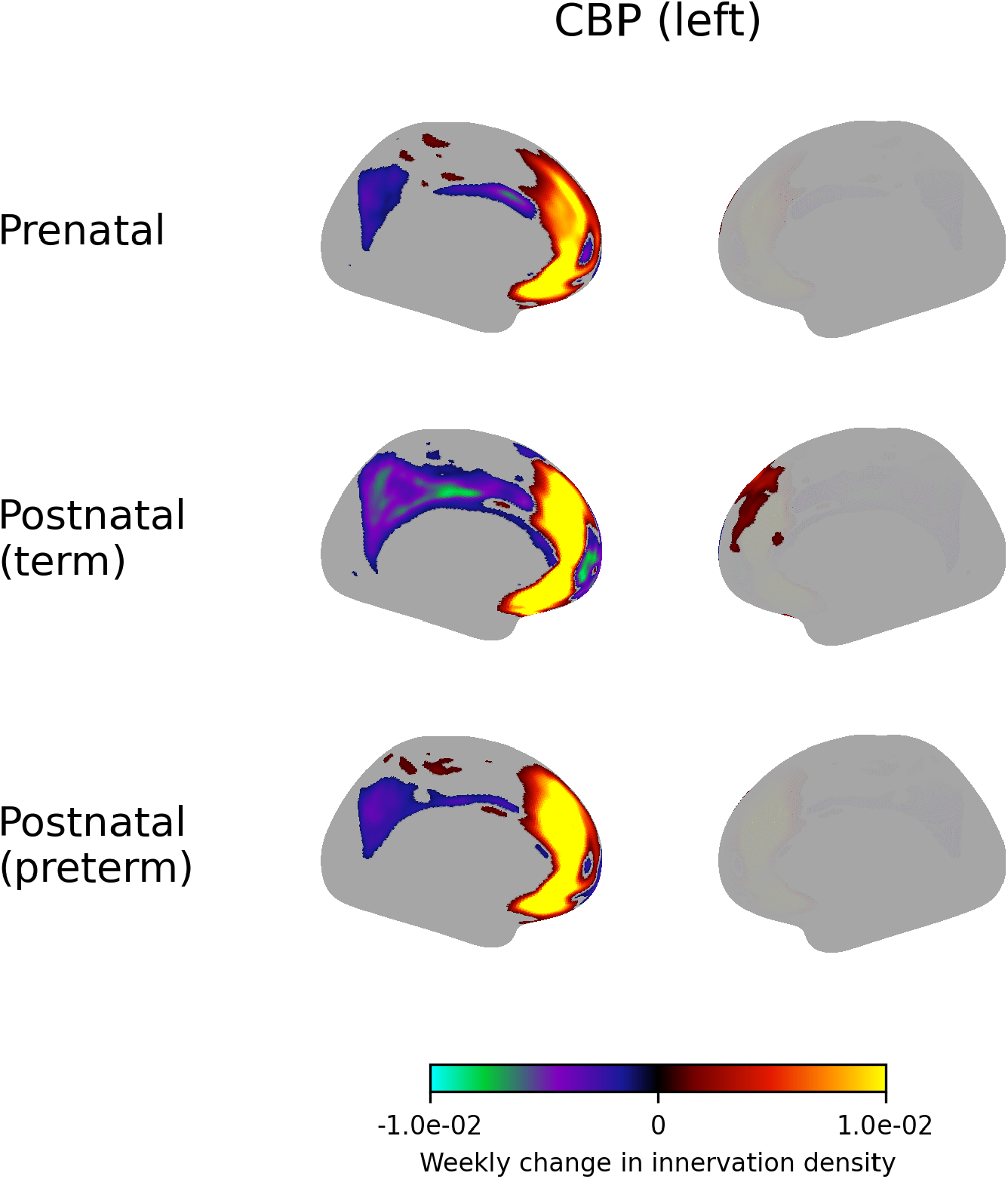
Weekly change in innervation density of the left perigenual cingulum bundle.

**Figure S33.**
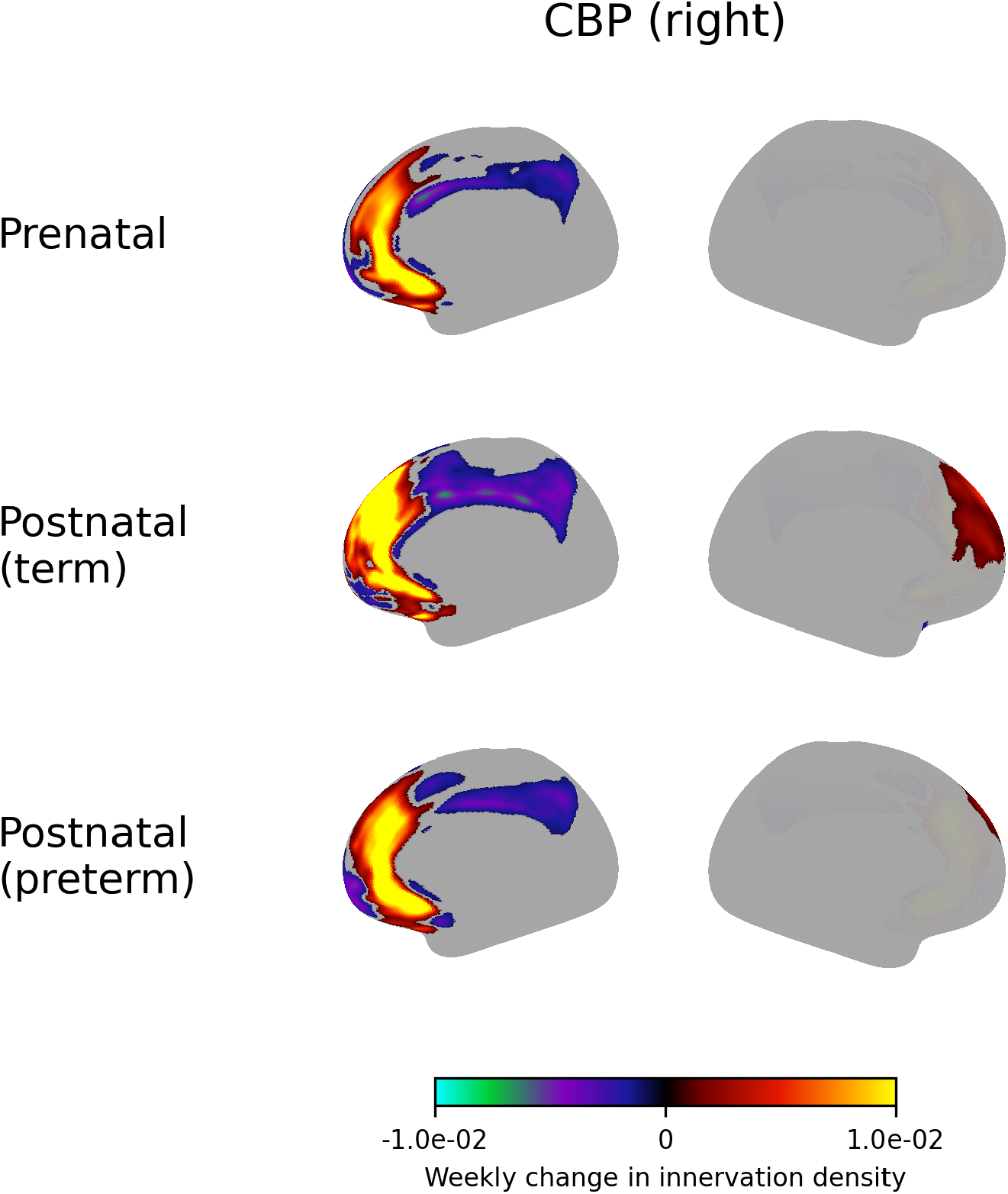
Weekly change in innervation density of the right perigenual cingulum bundle.

**Figure S34.**
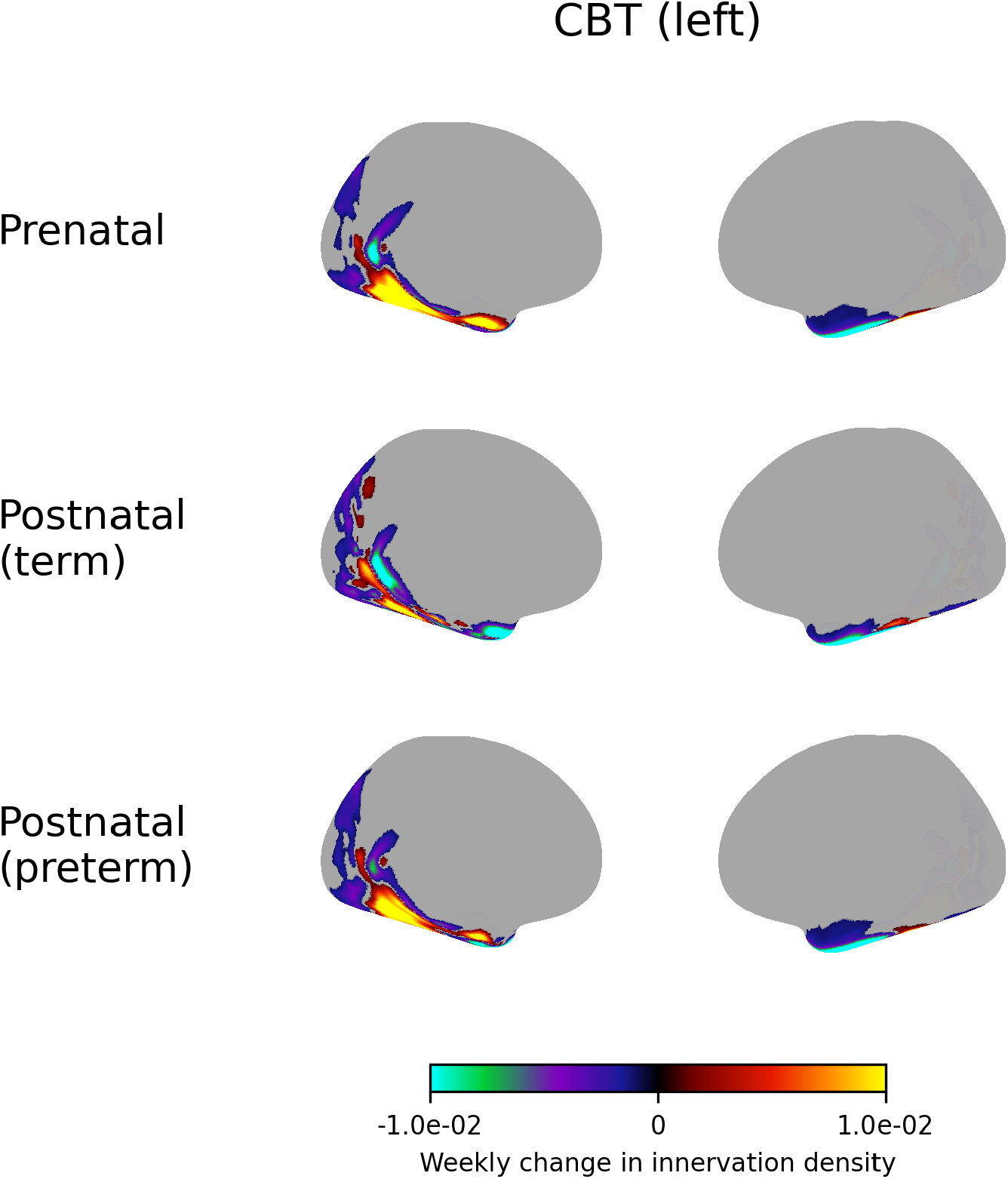
Weekly change in innervation density of the left temporal cingulum bundle.

**Figure S35.**
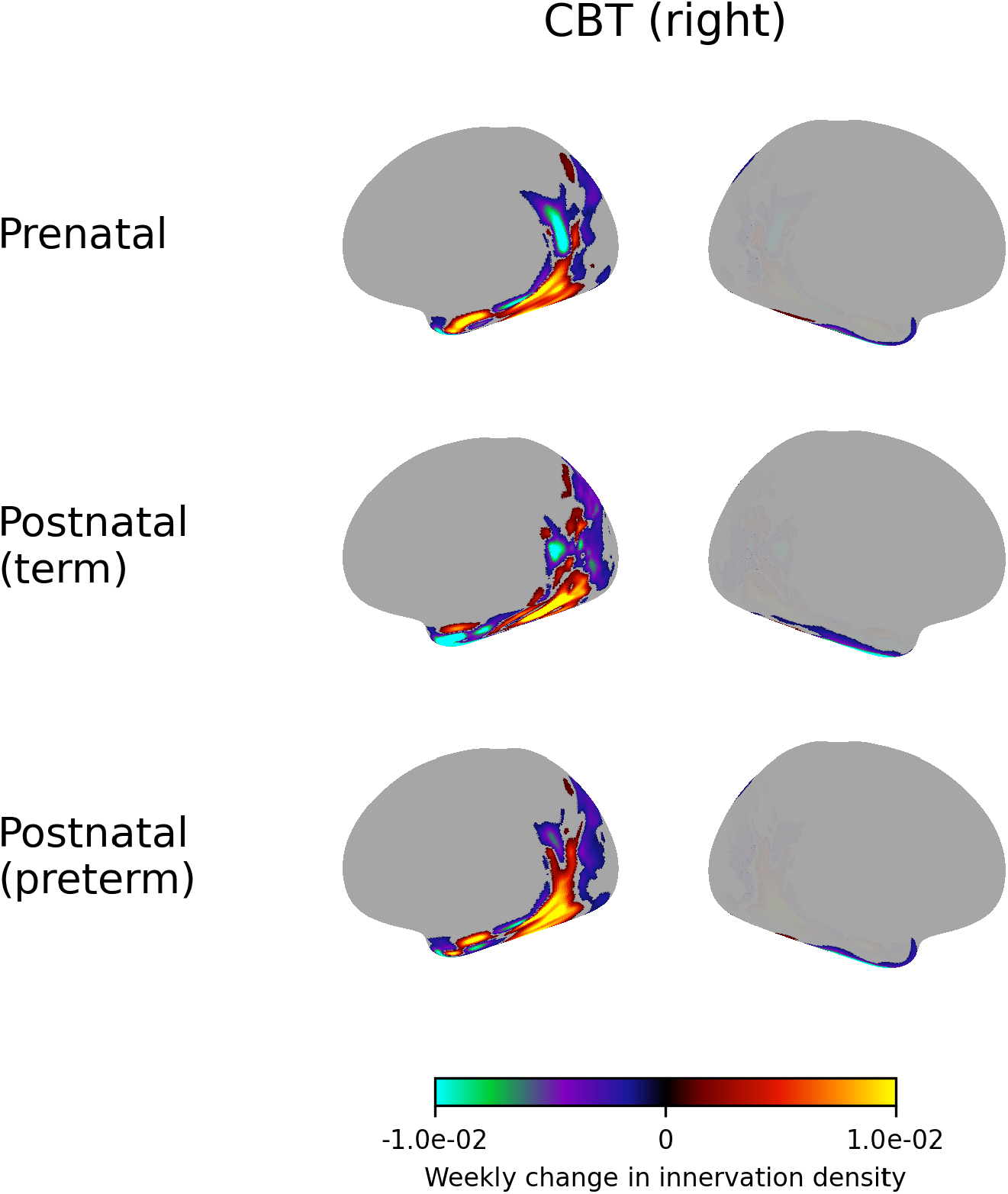
Weekly change in innervation density of the right temporal cingulum bundle.

**Figure S36.**
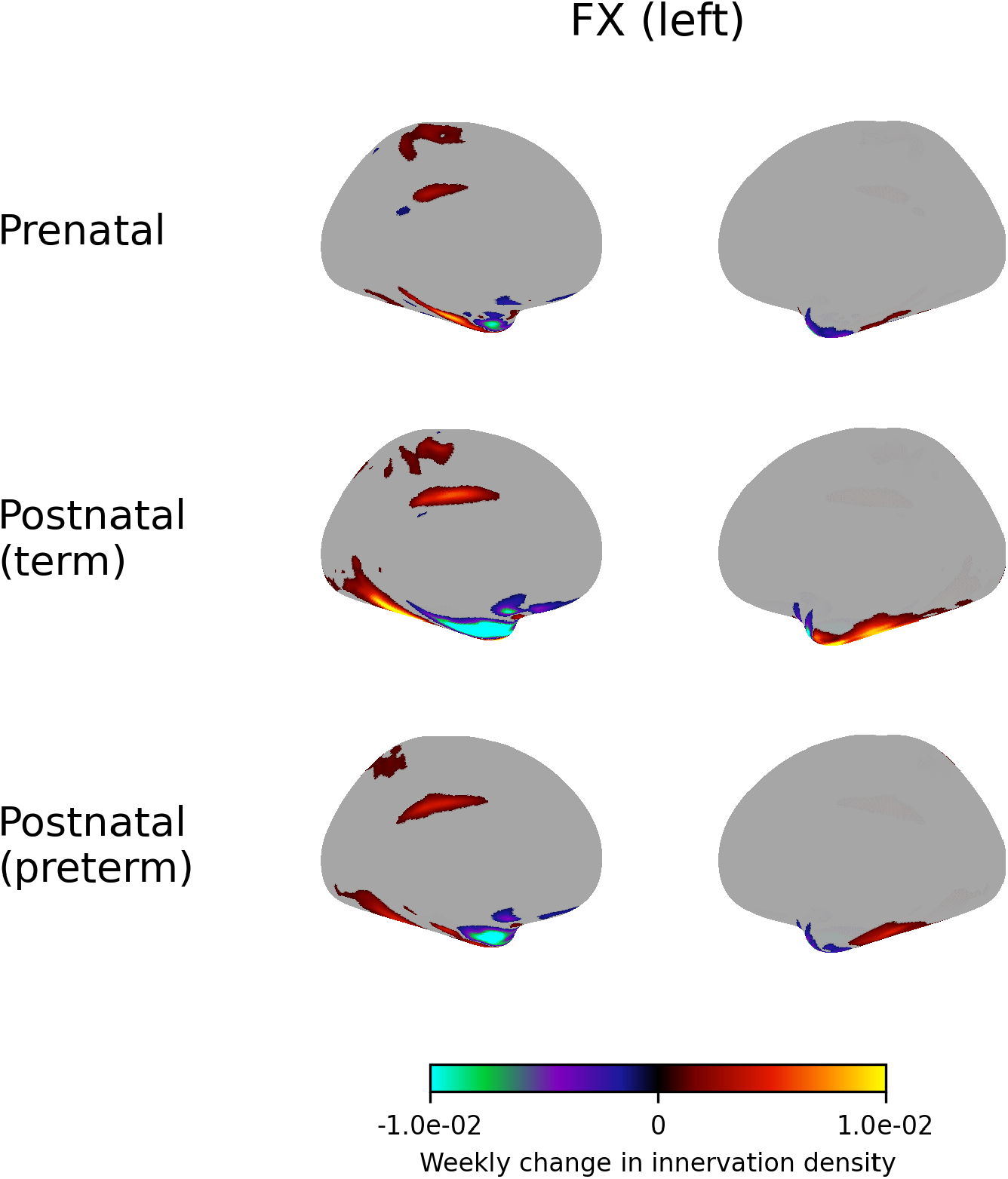
Weekly change in innervation density of the left fornix.

**Figure S37.**
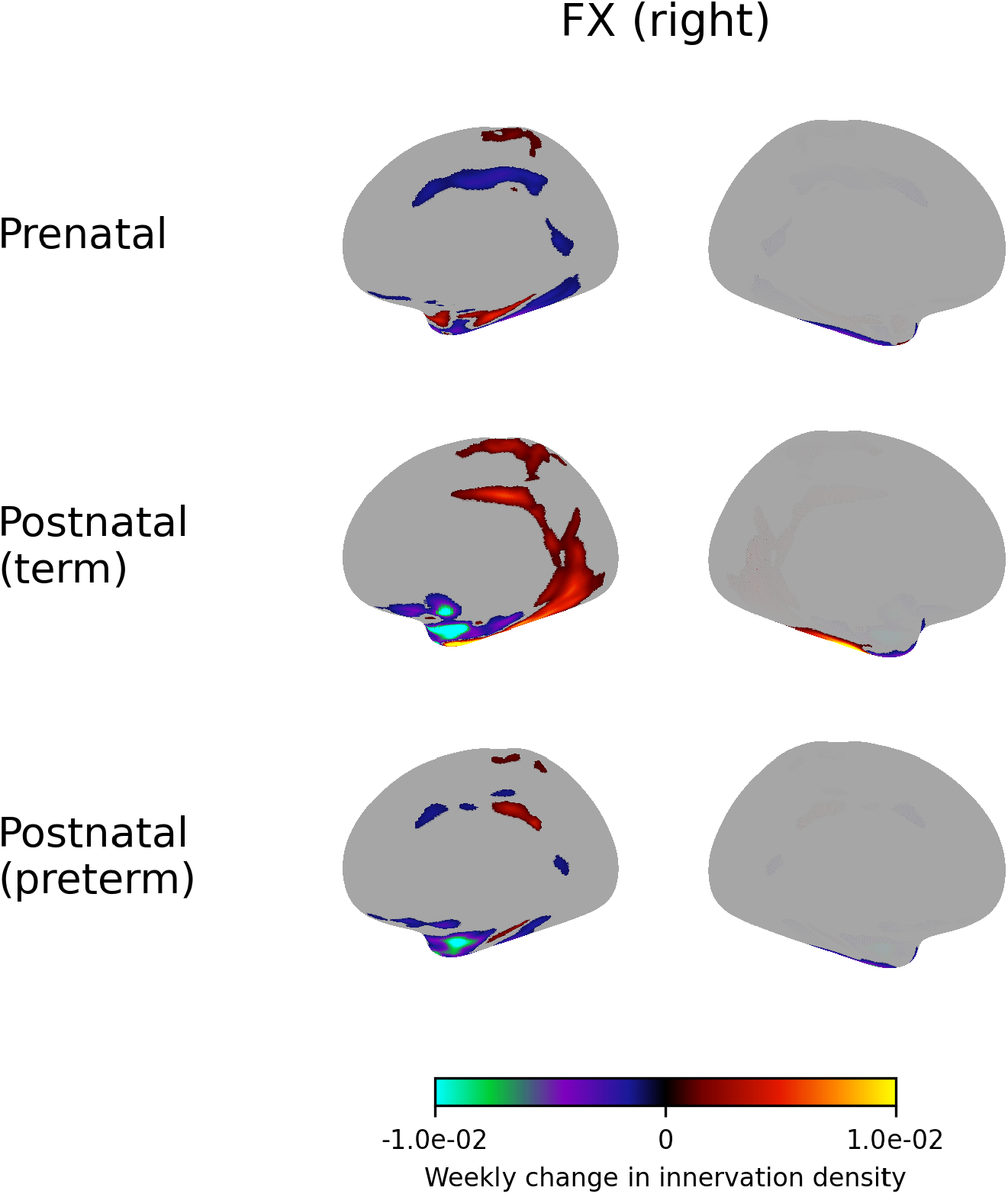
Weekly change in innervation density of the right fornix.

**Figure S38.**
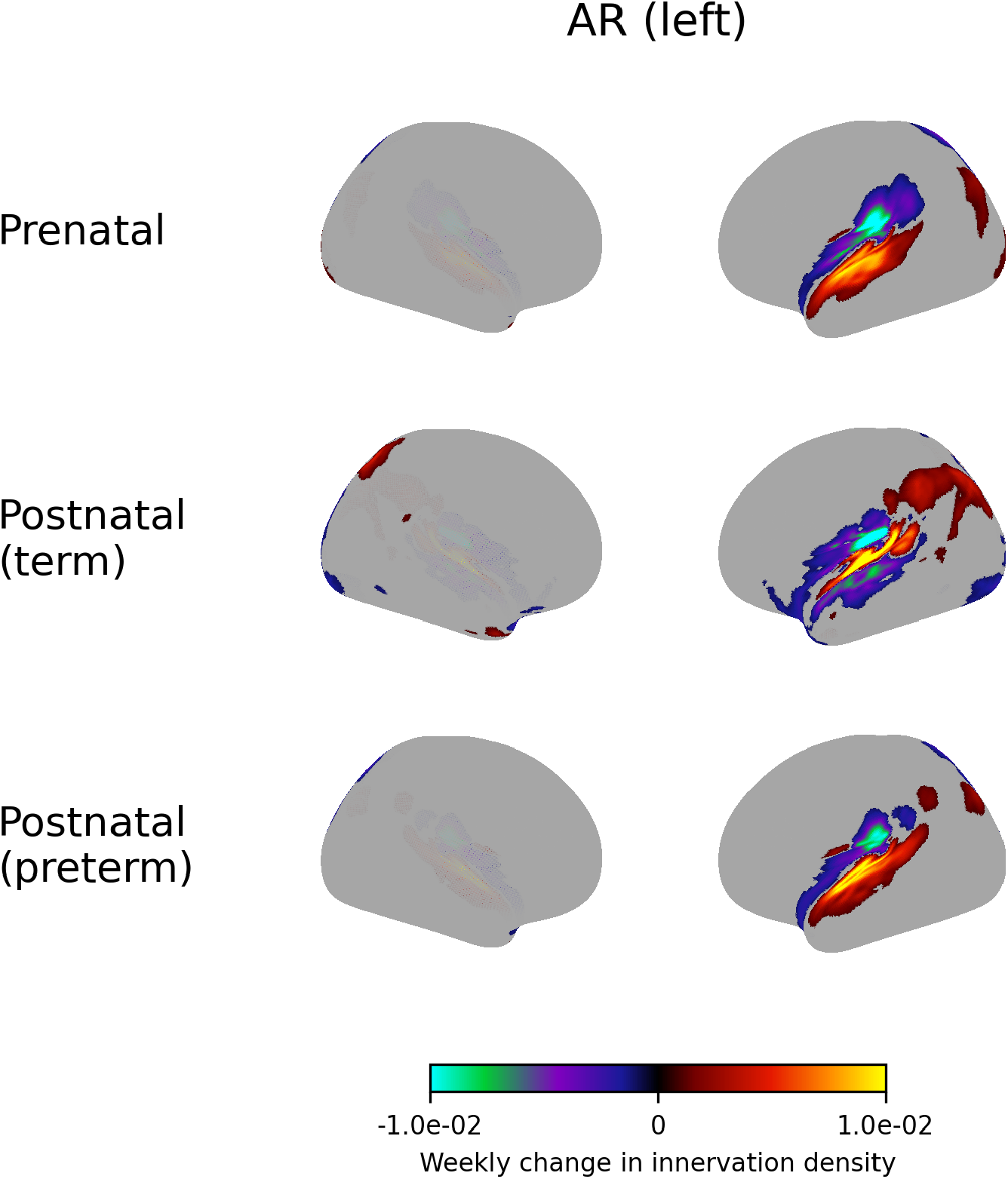
Weekly change in innervation density of the left acoustic radiation.

**Figure S39.**
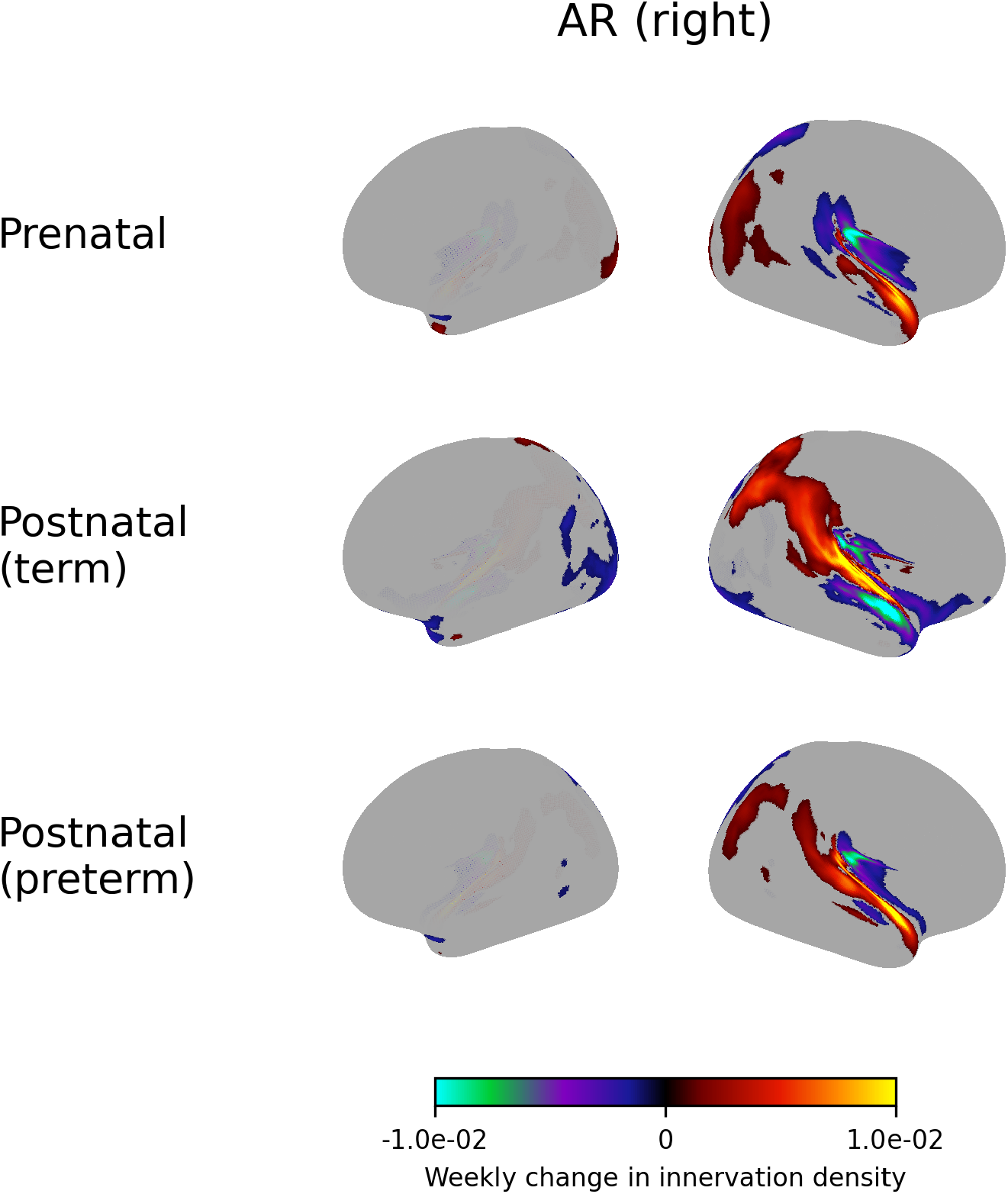
Weekly change in innervation density of the right acoustic radiation.

**Figure S40.**
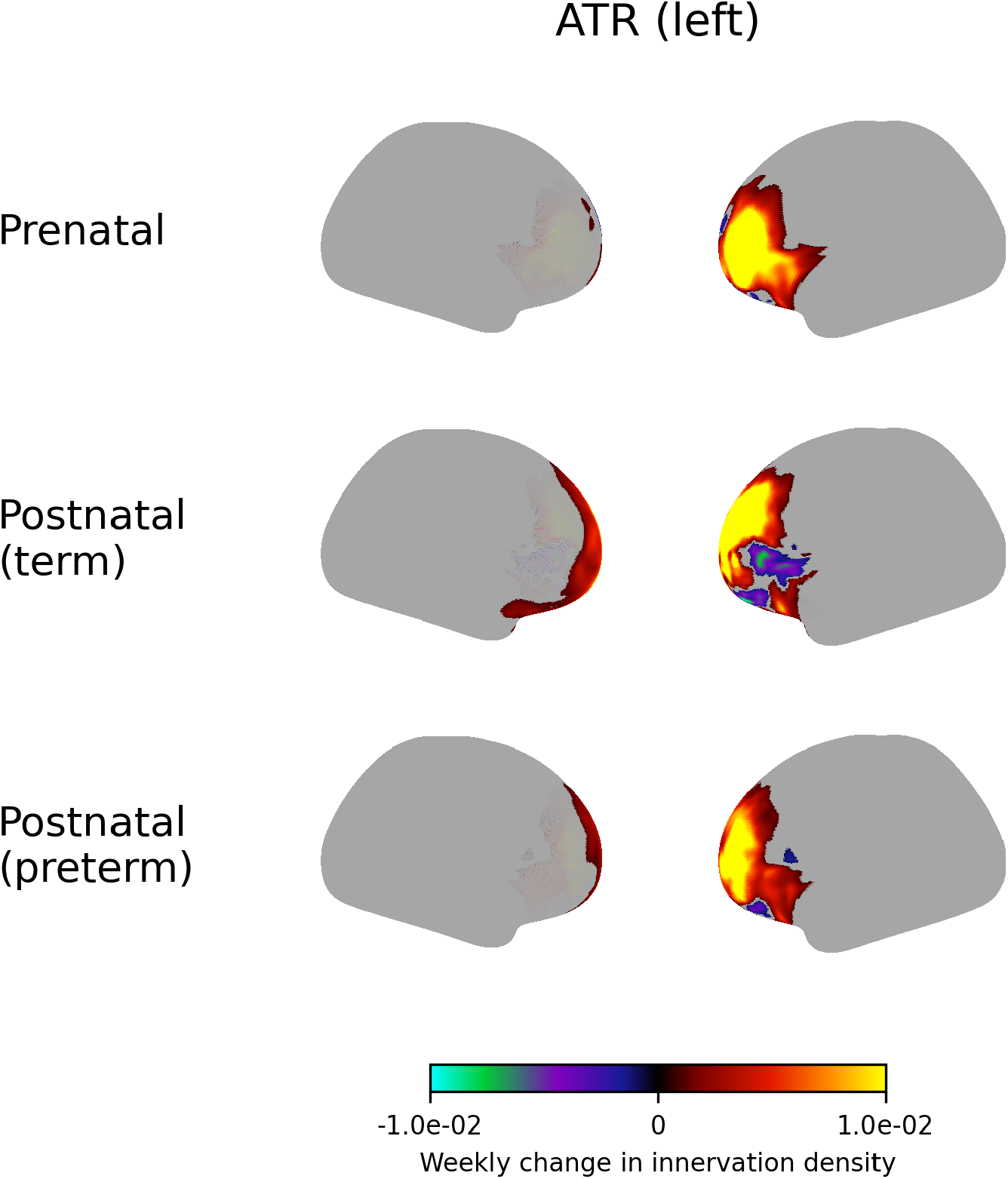
Weekly change in innervation density of the left anterior thalamic radiation.

**Figure S41.**
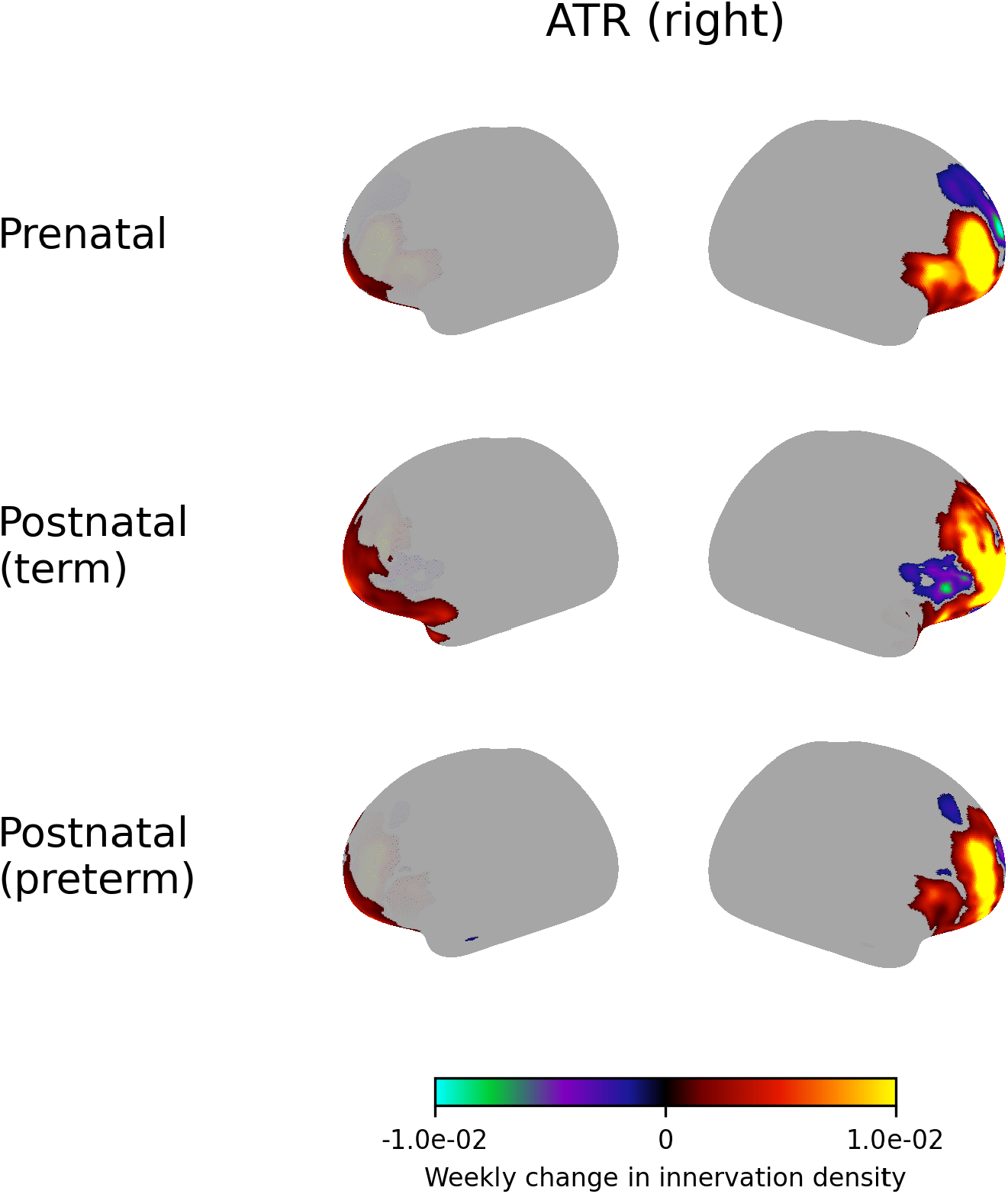
Weekly change in innervation density of the right anterior thalamic radiation.

**Figure S42.**
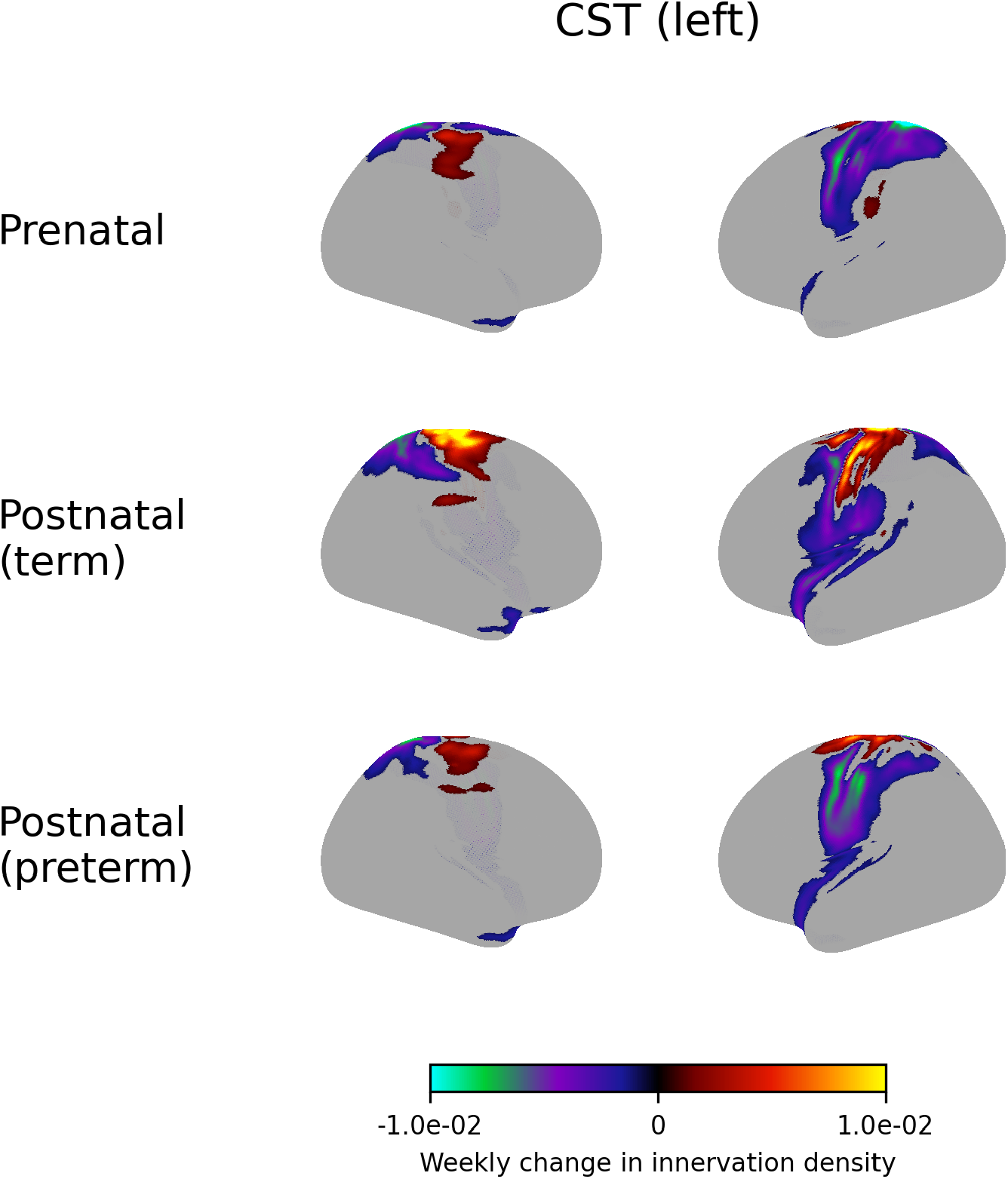
Weekly change in innervation density of the left corticospinal tract.

**Figure S43.**
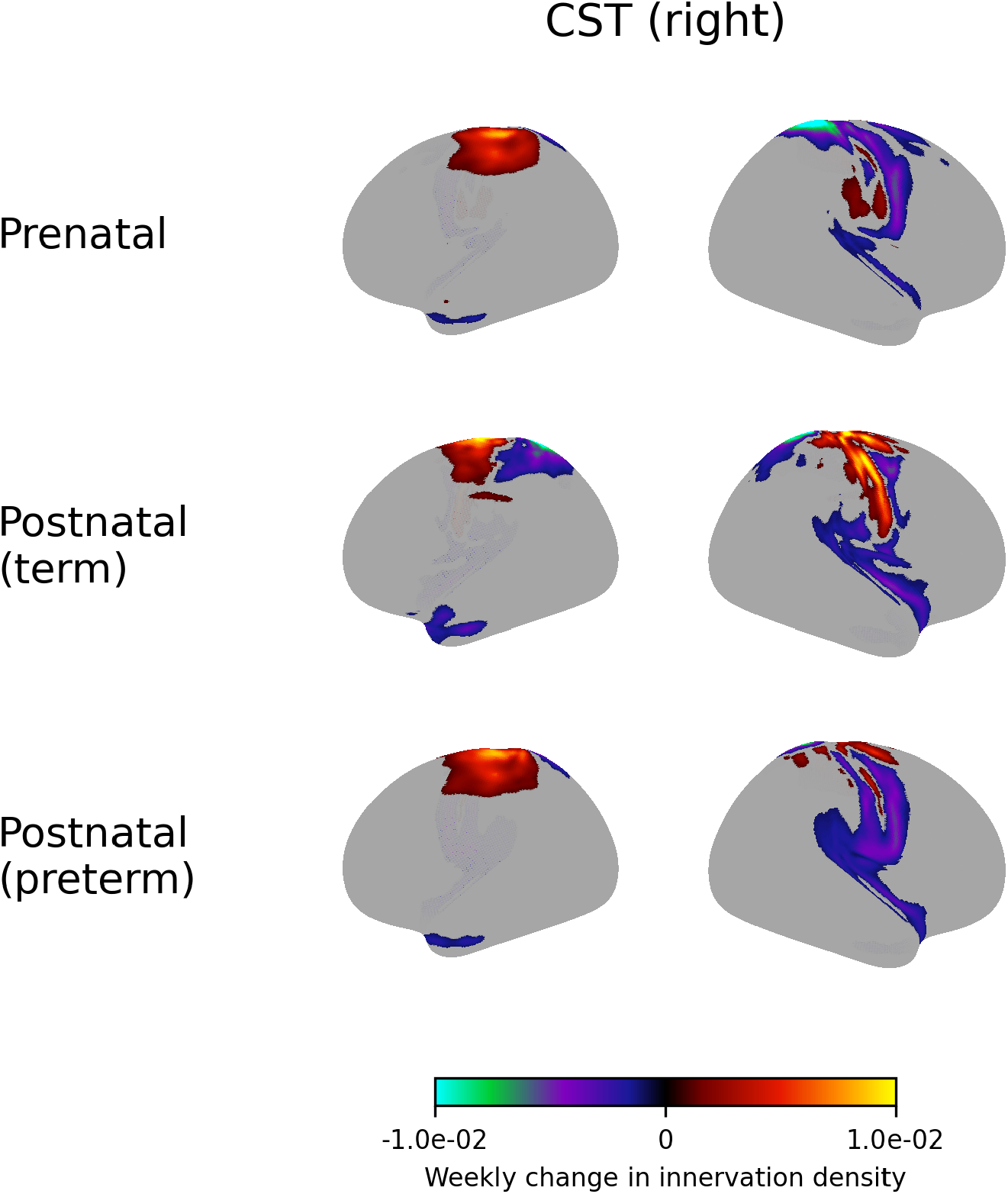
Weekly change in innervation density of the right corticospinal tract.

**Figure S44.**
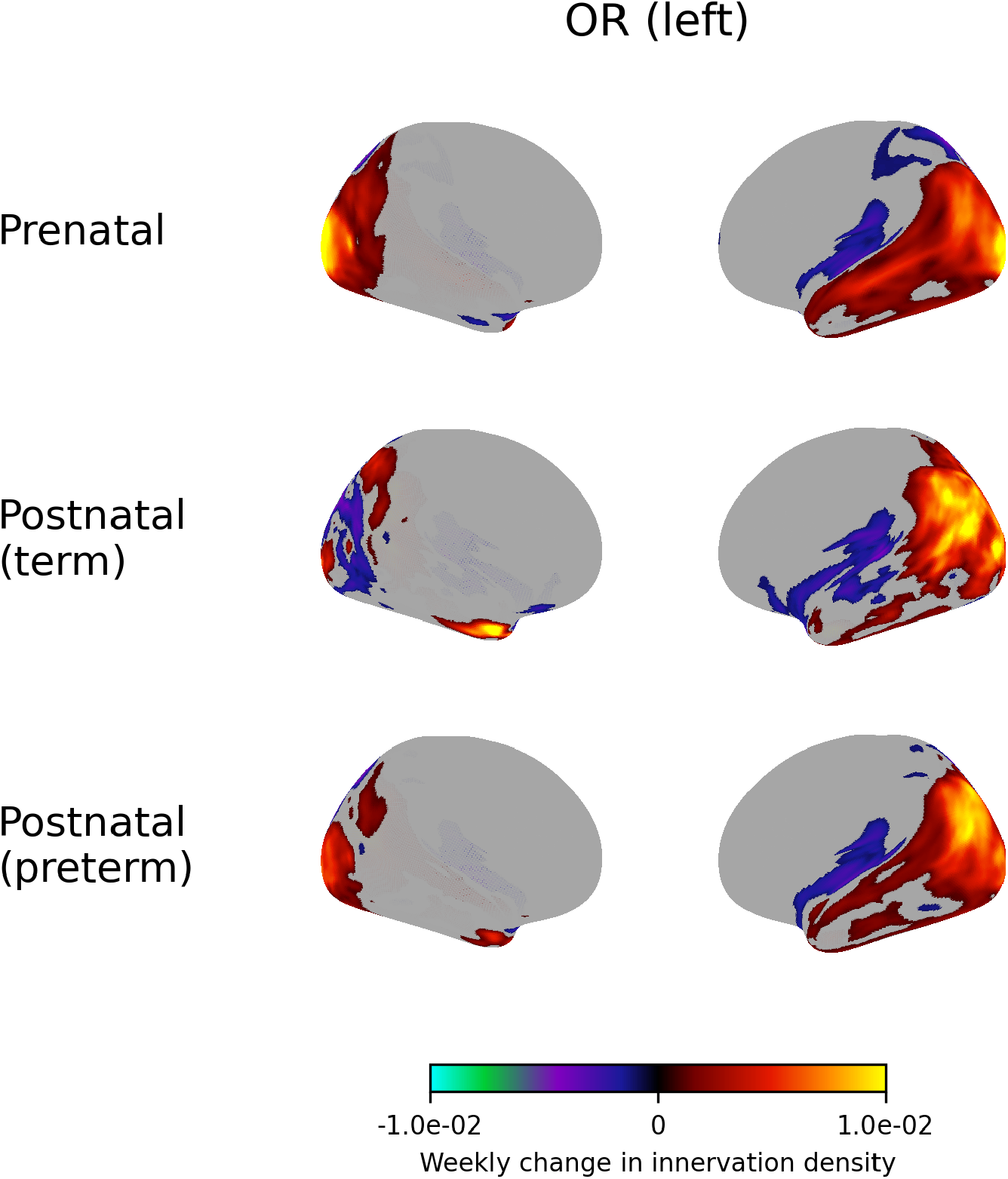
Weekly change in innervation density of the left optic radiation.

**Figure S45.**
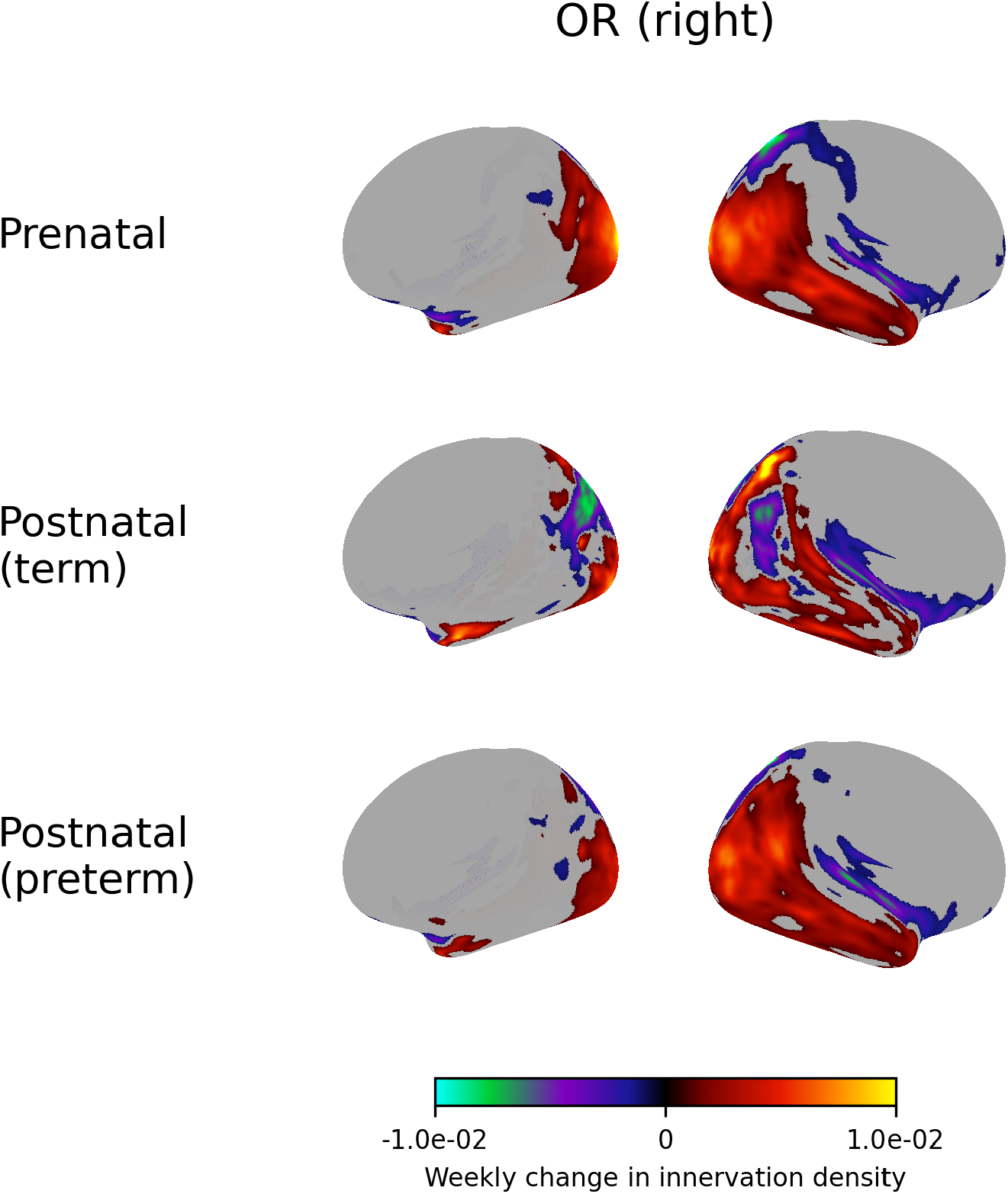
Weekly change in innervation density of the right optic radiation.

**Figure S46.**
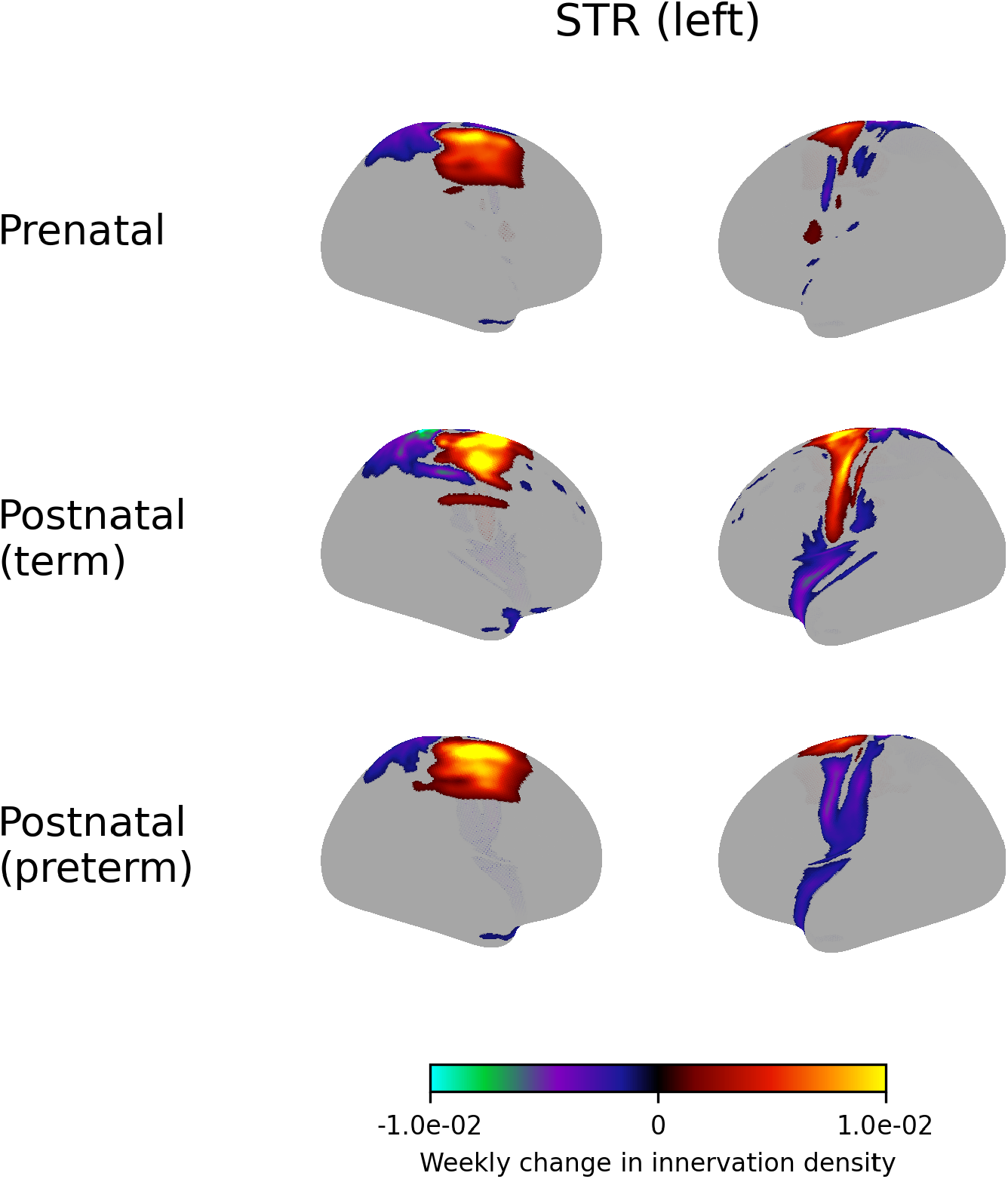
Weekly change in innervation density of the left superior thalamic radiation.

**Figure S47.**
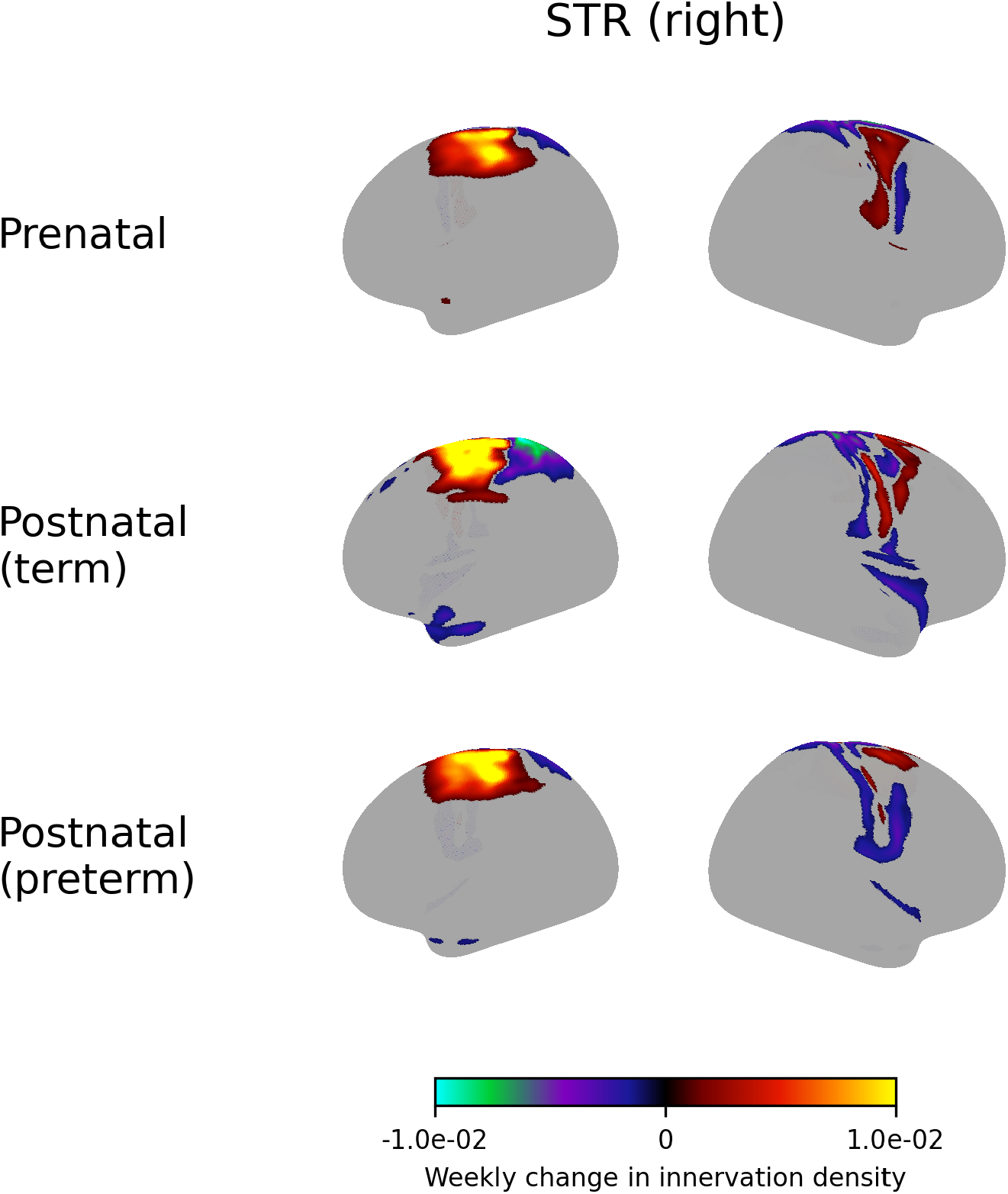
Weekly change in innervation density of the right superior thalamic radiation.

**Figure S48.**
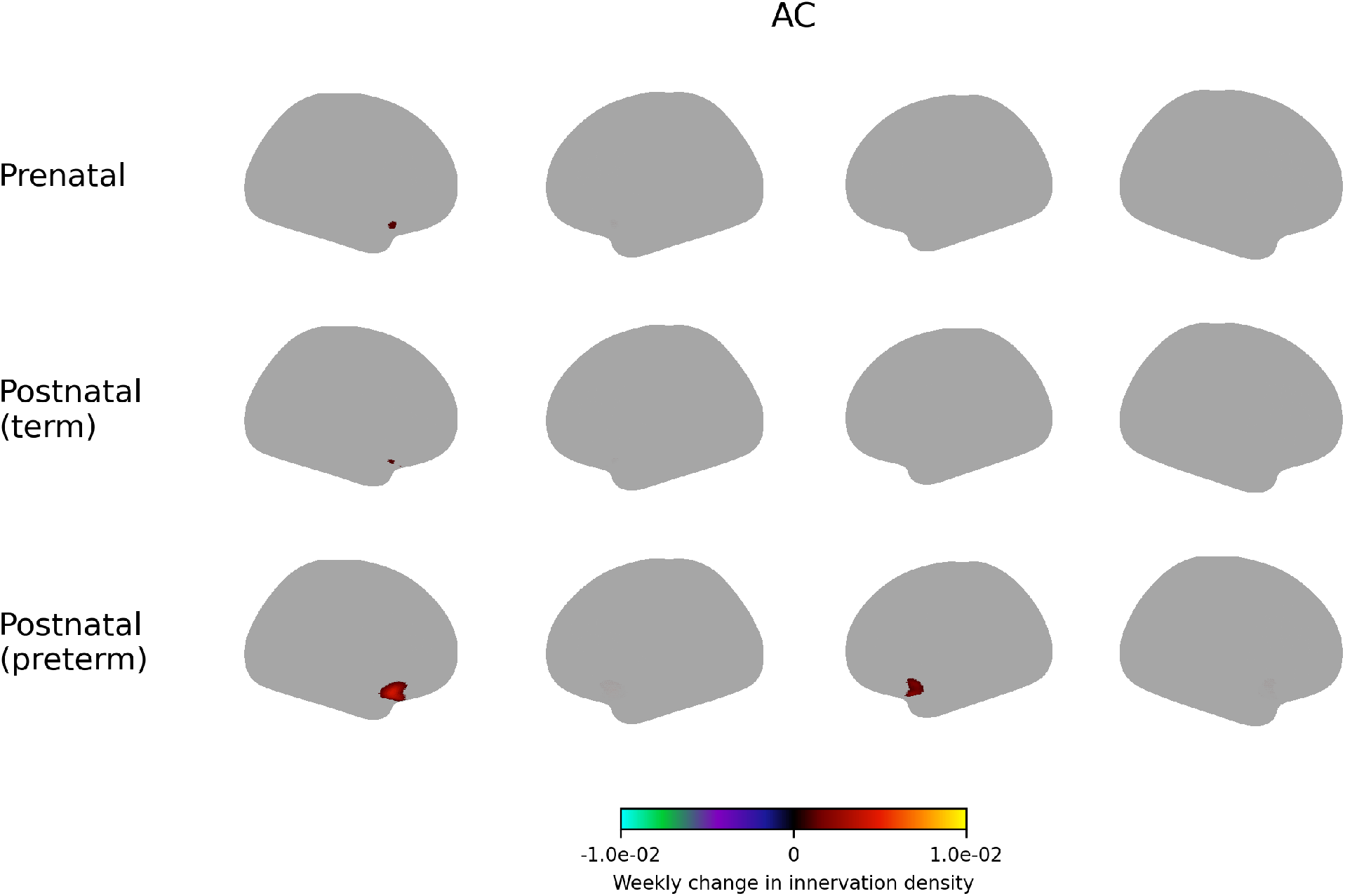
Weekly change in innervation density of the anterior commissure.

**Figure S49.**
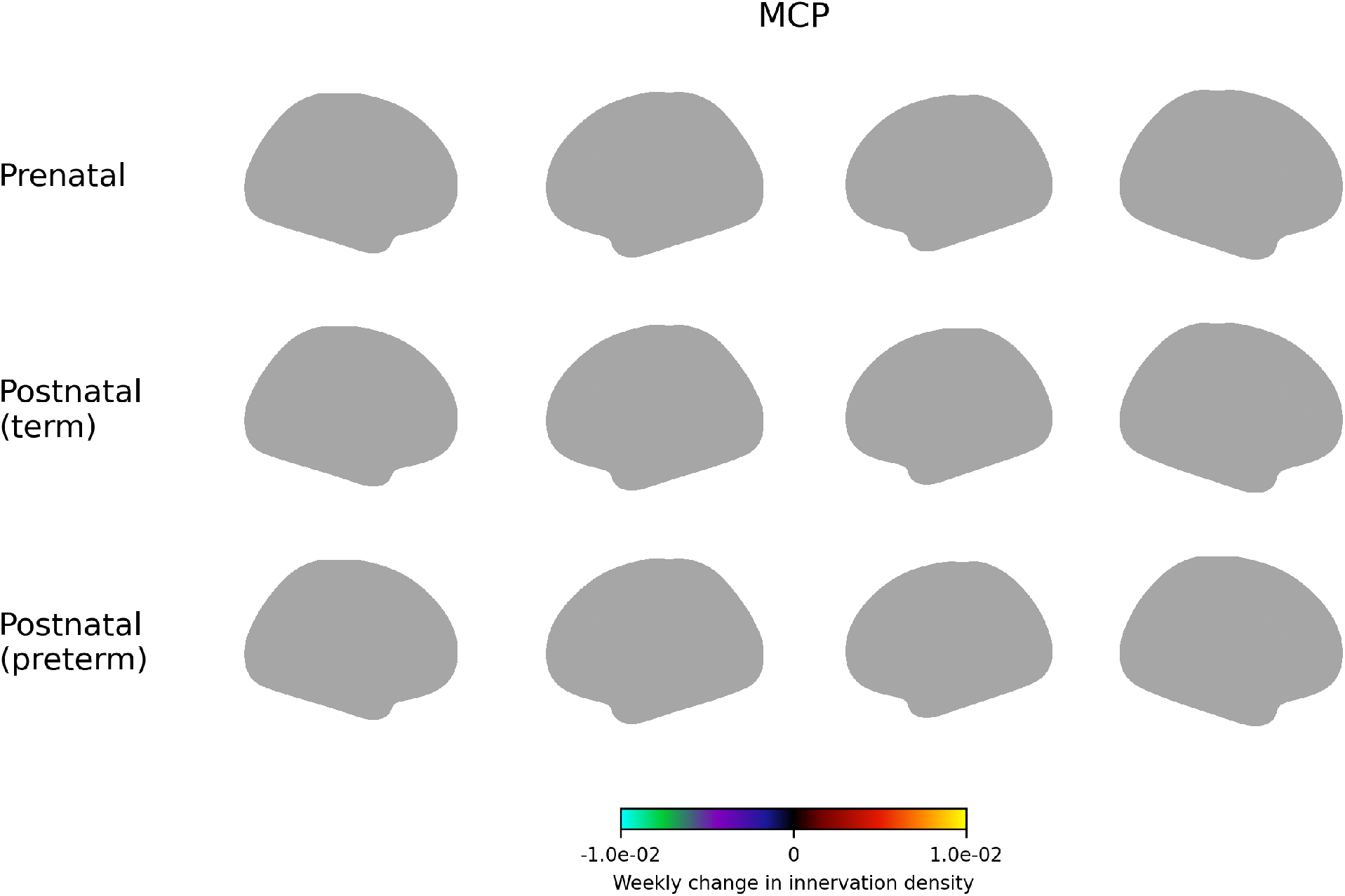
Weekly change in innervation density of the middle cerebellar peduncle. This tract does not reach the cerebral cortex, contributed zero density at all vertices.

**Figure S50.**
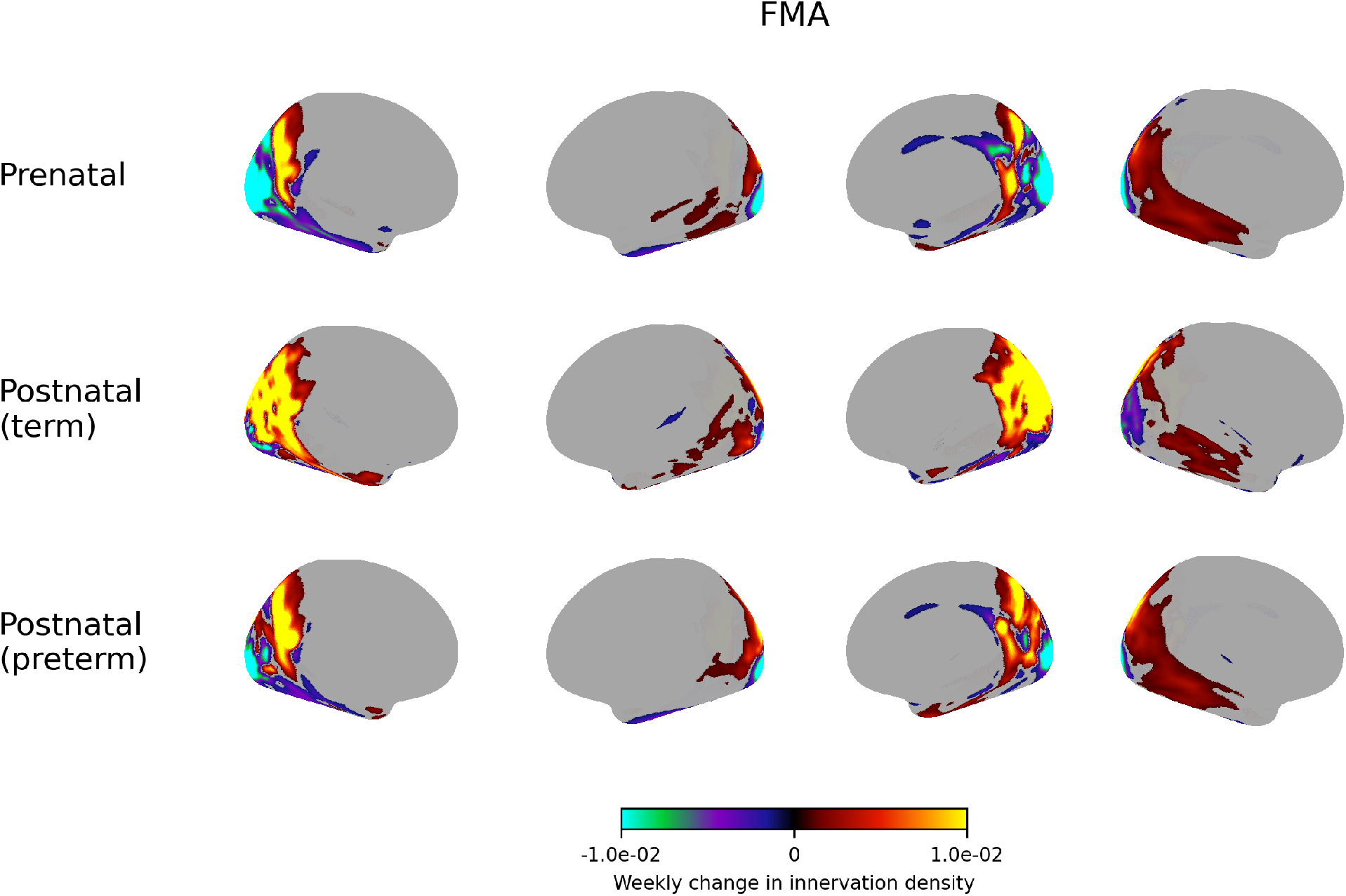
Weekly change in innervation density of the forceps major.

**Figure S51.**
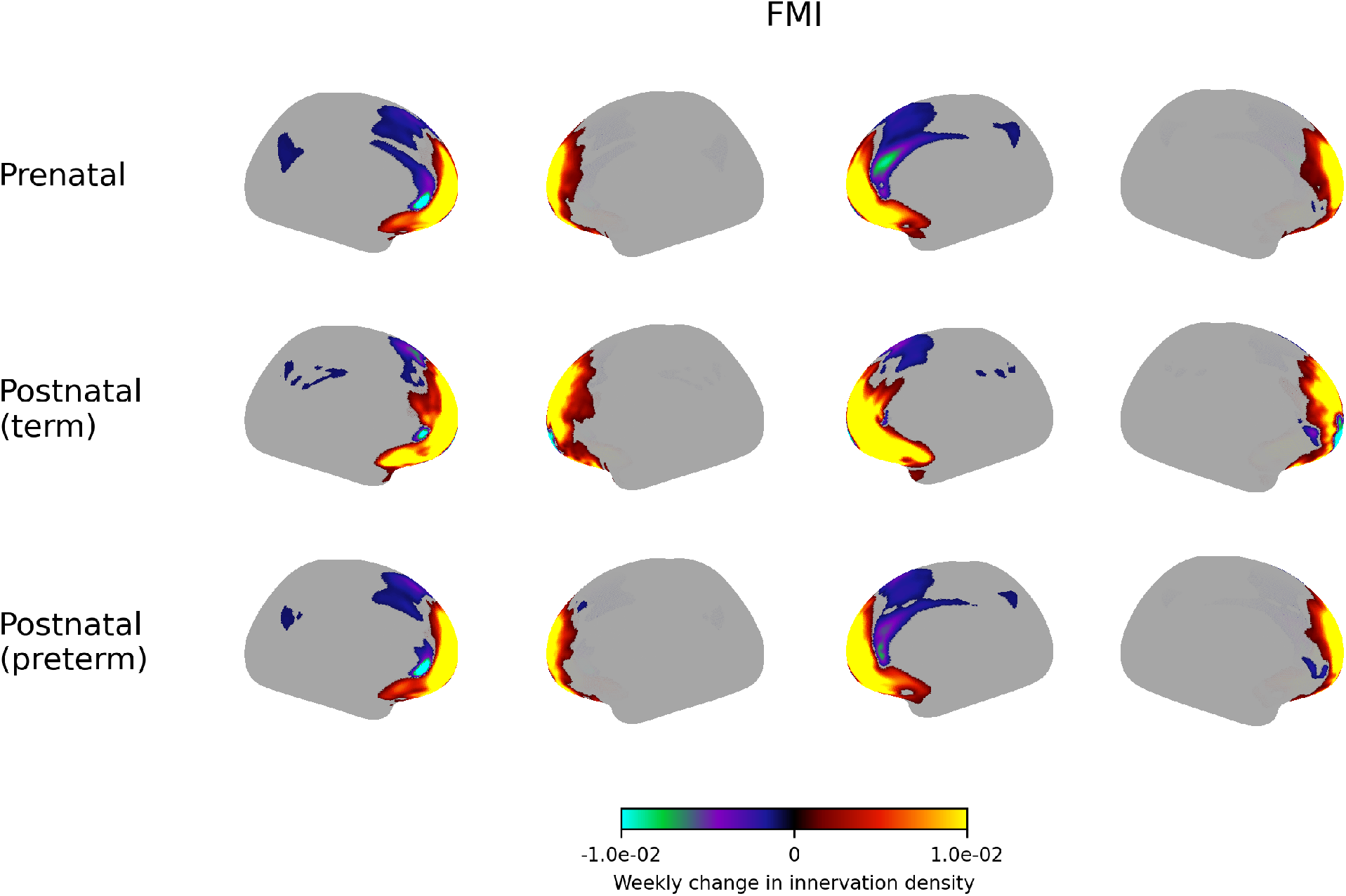
Weekly change in innervation density of the forceps minor.

## REFERENCES

Ajayi-Obe, M., Saeed, N., Cowan, F., Rutherford, M., and Edwards, A. (2000). “Reduced development of cerebral cortex in extremely preterm infants.” The Lancet, 356(9236), 1162–1163.

Alexander-Bloch, A. F., Shou, H., Liu, S., Satterthwaite, T. D., Glahn, D. C., Shinohara, R. T., Vandekar, S. N., and Raznahan, A. (2018). “On testing for spatial correspondence between maps of human brain structure and function.” NeuroImage, 178, 540–551.

Andersson, J. L., Skare, S., and Ashburner, J. (2003). “How to correct susceptibility distortions in spin-echo echo-planar images: application to diffusion tensor imaging.” NeuroImage, 20(2), 870–888.

Andersson, J. L. and Sotiropoulos, S. N. (2016). “An integrated approach to correction for off- resonance effects and subject movement in diffusion MR imaging.” NeuroImage, 125, 1063–1078.

Bastiani, M., Andersson, J. L., Cordero-Grande, L., Murgasova, M., Hutter, J., Price, A. N., Makropoulos, A., Fitzgibbon, S. P., Hughes, E., Rueckert, D., Victor, S., Rutherford, M., Edwards, A. D., Smith, S. M., Tournier, J.-D., Hajnal, J. V., Jbabdi, S., and Sotiropoulos, S. N. (2019). “Automated processing pipeline for neonatal diffusion mri in the developing human connectome project.” NeuroImage, 185, 750–763.

Basu, S. K., Pradhan, S., Kapse, K., McCarter, R., Murnick, J., Chang, T., and Limperopoulos, C. (2019). “Third Trimester Cerebellar Metabolite Concentrations are Decreased in Very Premature Infants with Structural Brain Injury.” Scientific Reports, 9(1).

Behrens, T., Berg, H. J., Jbabdi, S., Rushworth, M., and Woolrich, M. (2007). “Probabilistic diffusion tractography with multiple fibre orientations: What can we gain?.” NeuroImage, 34(1), 144–155.

Bourgeois, J. P., Jastreboff, P. J., and Rakic, P. (1989). “Synaptogenesis in visual cortex of normal and preterm monkeys: evidence for intrinsic regulation of synaptic overproduction..” Proceedings of the National Academy of Sciences, 86(11), 4297–4301.

Bouyssi-Kobar, M., De Asis-Cruz, J., Murnick, J., Chang, T., and Limperopoulos, C. (2019). “Altered functional brain network integration, segregation, and modularity in infants born very preterm at term-equivalent age.” The Journal of Pediatrics, 213, 13–21.e1.

Bouyssi-Kobar, M., du Plessis, A. J., McCarter, R., Brossard-Racine, M., Murnick, J., Tinkleman, L., Robertson, R. L., and Limperopoulos, C. (2016). “Third trimester brain growth in preterm infants compared with in utero healthy fetuses.” Pediatrics, 138(5), e20161640.

Cordero-Grande, L., Hughes, E. J., Hutter, J., Price, A. N., and Hajnal, J. V. (2017). “Three- dimensional motion corrected sensitivity encoding reconstruction for multi-shot multi-slice mri: Application to neonatal brain imaging.” Magnetic Resonance in Medicine, 79(3), 1365–1376.

De Asis-Cruz, J., Kapse, K., Basu, S. K., Said, M., Scheinost, D., Murnick, J., Chang, T., du Plessis, A., and Limperopoulos, C. (2020). “Functional brain connectivity in ex utero premature infants compared to in utero fetuses.” NeuroImage, 219, 117043.

Deoni, S. C., Dean, III, D. C., Remer, J., Dirks, H., and O’Muircheartaigh, J. (2015). “Cortical maturation and myelination in healthy toddlers and young children.” NeuroImage, 115, 147–161.

Dong, H.-M., Margulies, D. S., Zuo, X.-N., and Holmes, A. J. (2021). “Shifting gradients of macroscale cortical organization mark the transition from childhood to adolescence.” Proceedings of the National Academy of Sciences, 118(28).

Dubois, J., Dehaene-Lambertz, G., Kulikova, S., Poupon, C., Hüppi, P., and Hertz-Pannier, L. (2014). “The early development of brain white matter: A review of imaging studies in fetuses, newborns and infants.” Neuroscience, 276, 48–71.

Edwards, A. D., Rueckert, D., Smith, S. M., Abo Seada, S., Alansary, A., Almalbis, J., Allsop, J., Andersson, J., Arichi, T., Arulkumaran, S., Bastiani, M., Batalle, D., Baxter, L., Bozek, J., Braithwaite, E., Brandon, J., Carney, O., Chew, A., Christiaens, D., Chung, R., Colford, K., Cordero-Grande, L., Counsell, S. J., Cullen, H., Cupitt, J., Curtis, C., Davidson, A., Deprez, M., Dillon, L., Dimitrakopoulou, K., Dimitrova, R., Duff, E., Falconer, S., Farahibozorg, S.-R., Fitzgibbon, S. P., Gao, J., Gaspar, A., Harper, N., Harrison, S. J., Hughes, E. J., Hutter, J., Jenkinson, M., Jbabdi, S., Jones, E., Karolis, V., Kyriakopoulou, V., Lenz, G., Makropoulos, A., Malik, S., Mason, L., Mortari, F., Nosarti, C., Nunes, R. G., O’Keeffe, C., O’Muircheartaigh, J., Patel, H., Passerat-Palmbach, J., Pietsch, M., Price, A. N., Robinson, E. C., Rutherford, M. A., Schuh, A., Sotiropoulos, S., Steinweg, J., Teixeira, R. P. A. G., Tenev, T., Tournier, J.-D., Tusor, N., Uus, A., Vecchiato, K., Williams, L. Z. J., Wright, R., Wurie, J., and Hajnal, J. V. (2022). “The developing human connectome project neonatal data release.” Frontiers in Neuroscience, 16.

Gao, W., Lin, W., Chen, Y., Gerig, G., Smith, J., Jewells, V., and Gilmore, J. (2008). “Temporal and spatial development of axonal maturation and myelination of white matter in the developing brain.” American Journal of Neuroradiology, 30(2), 290–296.

Gholipour, A., Rollins, C. K., Velasco-Annis, C., Ouaalam, A., Akhondi-Asl, A., Afacan, O., Ortinau, C. M., Clancy, S., Limperopoulos, C., Yang, E., Estroff, J. A., and Warfield, S. K. (2017). “A normative spatiotemporal mri atlas of the fetal brain for automatic segmentation and analysis of early brain growth.” Scientific Reports, 7(1).

Glasser, M. F., Coalson, T. S., Robinson, E. C., Hacker, C. D., Harwell, J., Yacoub, E., Ugurbil, K., Andersson, J., Beckmann, C. F., Jenkinson, M., Smith, S. M., and Van Essen, D. C. (2016). “A multi-modal parcellation of human cerebral cortex.” Nature, 536(7615), 171–178.

Gogtay, N., Giedd, J. N., Lusk, L., Hayashi, K. M., Greenstein, D., Vaituzis, A. C., Nugent, III, T. F., Herman, D. H., Clasen, L. S., Toga, A. W., Rapoport, J. L., and Thompson, P. M. (2004). “Dynamic mapping of human cortical development during childhood through early adulthood.” Proceedings of the National Academy of Sciences, 101(21), 8174–8179.

Gong, Q., Roberts, N., Garden, A., and Whitehouse, G. (1998). “Fetal and fetal brain volume estimation in the third trimester of human pregnancy using gradient echo mr imaging.” Magnetic Resonance Imaging, 16(3), 235–240.

Greve, D. N. and Fischl, B. (2009). “Accurate and robust brain image alignment using boundary-based registration.” NeuroImage, 48(1), 63–72.

Grotheer, M., Bloom, D., Kruper, J., Richie-Halford, A., Zika, S., Aguilera González, V. A., Yeatman, J. D., Grill-Spector, K., and Rokem, A. (2023). “Human white matter myelinates faster in utero than ex utero.” Proceedings of the National Academy of Sciences, 120(33).

Holland, D., Chang, L., Ernst, T. M., Curran, M., Buchthal, S. D., Alicata, D., Skranes, J., Johansen, H., Hernandez, A., Yamakawa, R., Kuperman, J. M., and Dale, A. M. (2014). “Structural growth trajectories and rates of change in the first 3 months of infant brain development.” JAMA Neurology, 71(10), 1266.

Innocenti, G. M. and Price, D. J. (2005). “Exuberance in the development of cortical networks.” Nature Reviews Neuroscience, 6(12), 955–965.

Jbabdi, S., Sotiropoulos, S. N., Savio, A. M., Graña, M., and Behrens, T. E. J. (2012). “Modelbased analysis of multishell diffusion MR data for tractography: How to get over fitting problems.” Magnetic Resonance in Medicine, 68(6), 1846–1855.

Johnson, S. and Marlow, N. (2011). “Preterm birth and childhood psychiatric disorders.” Pediatric Research, 69(5 Part 2), 11R–18R.

Knickmeyer, R. C., Gouttard, S., Kang, C., Evans, D., Wilber, K., Smith, J. K., Hamer, R. M., Lin, W., Gerig, G., and Gilmore, J. H. (2008). “A structural mri study of human brain development from birth to 2 years.” The Journal of Neuroscience, 28(47), 12176–12182.

Kostović, I. and Jovanov-Milošević, N. (2006). “The development of cerebral connections during the first 20–45 weeks’ gestation.” Seminars in Fetal and Neonatal Medicine, 11(6), 415–422.

Kuklisova-Murgasova, M., Quaghebeur, G., Rutherford, M. A., Hajnal, J. V., and Schnabel, J. A. (2012). “Reconstruction of fetal brain mri with intensity matching and complete outlier removal.” Medical Image Analysis, 16(8), 1550–1564.

Kwon, S. H., Scheinost, D., Lacadie, C., Benjamin, J., Myers, E. H., Qiu, M., Schneider, K. C., Rothman, D. L., Constable, R. T., and Ment, L. R. (2014). “Gaba, resting-state connectivity and the developing brain.” Neonatology, 106(2), 149–155.

Lange, F. J., Arthofer, C., Bartsch, A., Douaud, G., McCarthy, P., Smith, S. M., and Anders son, J. L. R. (2024). “Mmorf—FSL’s MultiMOdal registration framework.” Imaging Neuroscience, 2.

Larsen, B., Sydnor, V. J., Keller, A. S., Yeo, B. T., and Satterthwaite, T. D. (2023). “A critical period plasticity framework for the sensorimotor–association axis of cortical neurodevelopment.” Trends in Neurosciences, 46(10), 847–862.

Lefèvre, J., Germanaud, D., Dubois, J., Rousseau, F., de Macedo Santos, I., Angleys, H., Mangin, J.-F., Hüppi, P. S., Girard, N., and De Guio, F. (2015). “Are developmental trajectories of cortical folding comparable between cross-sectional datasets of fetuses and preterm newborns?.” Cerebral Cortex, 26(7), 3023–3035.

Li, H., Yan, G., Luo, W., Liu, T., Wang, Y., Liu, R., Zheng, W., Zhang, Y., Li, K., Zhao, L., Limperopoulos, C., Zou, Y., and Wu, D. (2021). “Mapping fetal brain development based on automated segmentation and 4d brain atlasing.” Brain Structure and Function, 226(6), 1961–1972.

Limperopoulos, C., Soul, J. S., Gauvreau, K., Huppi, P. S., Warfield, S. K., Bassan, H., Robertson, R. L., Volpe, J. J., and du Plessis, A. J. (2005). “Late gestation cerebellar growth is rapid and impeded by premature birth.” Pediatrics, 115(3), 688–695.

Lyall, A. E., Shi, F., Geng, X., Woolson, S., Li, G., Wang, L., Hamer, R. M., Shen, D., and Gilmore, J. H. (2014). “Dynamic development of regional cortical thickness and surface area in early childhood.” Cerebral Cortex, 25(8), 2204–2212.

Makropoulos, A., Aljabar, P., Wright, R., Hüning, B., Merchant, N., Arichi, T., Tusor, N., Hajnal, J. V., Edwards, A. D., Counsell, S. J., and Rueckert, D. (2016). “Regional growth and atlasing of the developing human brain.” NeuroImage, 125, 456–478.

Makropoulos, A., Gousias, I. S., Ledig, C., Aljabar, P., Serag, A., Hajnal, J. V., Edwards, A. D., Counsell, S. J., and Rueckert, D. (2014). “Automatic whole brain mri segmentation of the developing neonatal brain.” IEEE Transactions on Medical Imaging, 33(9), 1818–1831.

Makropoulos, A., Robinson, E. C., Schuh, A., Wright, R., Fitzgibbon, S., Bozek, J., Counsell, S. J., Steinweg, J., Vecchiato, K., Passerat-Palmbach, J., Lenz, G., Mortari, F., Tenev, T., Duff, E. P., Bastiani, M., Cordero-Grande, L., Hughes, E., Tusor, N., Tournier, J.-D., Hutter, J., Price, A. N., Teixeira, R. P. A., Murgasova, M., Victor, S., Kelly, C., Rutherford, M. A., Smith, S. M., Edwards, A. D., Hajnal, J. V., Jenkinson, M., and Rueckert, D. (2018). “The developing human connectome project: A minimal processing pipeline for neonatal cortical surface reconstruction.” NeuroImage, 173, 88–112.

Malik, S., Vinukonda, G., Vose, L. R., Diamond, D., Bhimavarapu, B. B. R., Hu, F., Zia, M. T., Hevner, R., Zecevic, N., and Ballabh, P. (2013). “Neurogenesis continues in the third trimester of pregnancy and is suppressed by premature birth.” The Journal of Neuroscience, 33(2), 411–423.

Mars, R. B., Sotiropoulos, S. N., Passingham, R. E., Sallet, J., Verhagen, L., Khrapitchev, A. A., Sibson, N., and Jbabdi, S. (2018). “Whole brain comparative anatomy using connectivity blueprints.” eLife, 7.

Natu, V. S., Rosenke, M., Wu, H., Querdasi, F. R., Kular, H., Lopez-Alvarez, N., Grotheer, M., Berman, S., Mezer, A. A., and Grill-Spector, K. (2021). “Infants’ cortex undergoes microstructural growth coupled with myelination during development.” Communications Biology, 4(1).

Nishio, M., Liu, X., Mackey, A. P., and Arcaro, M. J. (2025). “Myelination across cortical hierarchies and depths in humans and macaques.” Report no., openRxiv, <10.1101/2025.02.06.636851> (feb 6).

Nosarti, C., Reichenberg, A., Murray, R. M., Cnattingius, S., Lambe, M. P., Yin, L., MacCabe, J., Rifkin, L., and Hultman, C. M. (2012). “Preterm birth and psychiatric disorders in young adult life.” Archives of General Psychiatry, 69(6).

Orasanu, E., Melbourne, A., Cardoso, M. J., Lomabert, H., Kendall, G. S., Robertson, N. J., Marlow, N., and Ourselin, S. (2016). “Cortical folding of the preterm brain: A longitudinal analysis of extremely preterm born neonates using spectral matching.” Brain and Behavior, 6(8).

Padilla, N., Alexandrou, G., Blennow, M., Lagercrantz, H., and Ådén, U. (2014). “Brain growth gains and losses in extremely preterm infants at term.” Cerebral Cortex, 25(7), 1897–1905.

Rajagopalan, V., Deoni, S., Panigrahy, A., and Thomason, M. E. (2021). “Is fetal MRI ready for neuroimaging prime time? An examination of progress and remaining areas for development.” Developmental Cognitive Neuroscience, 51, 100999.

Ricci, D., Cesarini, L., Romeo, D. M., Gallini, F., Serrao, F., Groppo, M., De Carli, A., Cota, F., Lepore, D., Molle, F., Ratiglia, R., De Carolis, M. P., Mosca, F., Romagnoli, C., Guzzetta, F., Cowan, F., Ramenghi, L. A., and Mercuri, E. (2008). “Visual function at 35 and 40 weeks’ postmenstrual age in low-risk preterm infants.” Pediatrics, 122(6), e1193–e1198.

Schmidt Mellado, G., Pillay, K., Adams, E., Alarcon, A., Andritsou, F., Cobo, M. M., Evans Fry, R., Fitzgibbon, S., Moultrie, F., Baxter, L., and Slater, R. (2022). “The impact of premature extrauterine exposure on infants’ stimulus-evoked brain activity across multiple sensory systems.” NeuroImage: Clinical, 33, 102914.

Schmierer, K., Wheeler-Kingshott, C. A., Boulby, P. A., Scaravilli, F., Altmann, D. R., Barker, G. J., Tofts, P. S., and Miller, D. H. (2007). “Diffusion tensor imaging of post mortem multiple sclerosis brain.” NeuroImage, 35(2), 467–477.

Schuh, A., Makropoulos, A., Robinson, E. C., Cordero-Grande, L., Hughes, E., Hutter, J., Price, A. N., Murgasova, M., Teixeira, R. P. A. G., Tusor, N., Steinweg, J. K., Victor, S., Rutherford, M. A., Hajnal, J. V., Edwards, A. D., and Rueckert, D. (2018). “Unbiased construction of a temporally consistent morphological atlas of neonatal brain development.” Report no., openRxiv, <10.1101/251512> (jan 22).

Schwindt, E., Giordano, V., Rona, Z., Czaba-Hnizdo, C., Olischar, M., Waldhoer, T., Werther, T., Fuiko, R., Berger, A., and Klebermass-Schrehof, K. (2018). “The impact of extrauterine life on visual maturation in extremely preterm born infants.” Pediatric Research, 84(3), 403–410.

Shaw, P., Kabani, N. J., Lerch, J. P., Eckstrand, K., Lenroot, R., Gogtay, N., Greenstein, D., Clasen, L., Evans, A., Rapoport, J. L., Giedd, J. N., and Wise, S. P. (2008). “Neurodevelopmental trajectories of the human cerebral cortex.” The Journal of Neuroscience, 28(14), 3586–3594.

Smyser, C. D., Inder, T. E., Shimony, J. S., Hill, J. E., Degnan, A. J., Snyder, A. Z., and Neil, J. J. (2010). “Longitudinal analysis of neural network development in preterm infants.” Cerebral Cortex, 20(12), 2852–2862.

Sydnor, V. J., Larsen, B., Bassett, D. S., Alexander-Bloch, A., Fair, D. A., Liston, C., Mackey, A. P., Milham, M. P., Pines, A., Roalf, D. R., Seidlitz, J., Xu, T., Raznahan, A., and Satterthwaite, T. D. (2021). “Neurodevelopment of the association cortices: Patterns, mechanisms, and implications for psychopathology.” Neuron, 109(18), 2820–2846.

Tau, G. Z. and Peterson, B. S. (2009). “Normal development of brain circuits.” Neuropsychophar-macology, 35(1), 147–168.

Tibrewal, M., Cheng, B., Dohare, P., Hu, F., Mehdizadeh, R., Wang, P., Zheng, D., Ungvari, Z., and Ballabh, P. (2017). “Disruption of interneuron neurogenesis in premature newborns and reversal with estrogen treatment.” The Journal of Neuroscience, 38(5), 1100–1113.

Toda, T., Homma, D., Tokuoka, H., Hayakawa, I., Sugimoto, Y., Ichinose, H., and Kawasaki, H. (2013). “Birth regulates the initiation of sensory map formation through serotonin signaling.” Developmental Cell, 27(1), 32–46.

Uus, A., Kyriakopoulou, V., Cordero Grande, L., Christiaens, D., Pietsch, M., Price, A., Wilson, S., Patkee, P., Karolis, S., Schuh, A., Gartner, A., Williams, L., Hughes, E., Arichi, T., O’Muircheartaigh, J., Hutter, J., Robinson, E., Tournier, J. D., Rueckert, D., Counsell, S., Rutherford, M., Deprez, M., Hajnal, J. V., and Edwards, A. D. (2023). “Multi-channel spatiotemporal mri atlas of the normal fetal brain development from the developing human connectome project, <https://doi.gin.g-node.org/10.12751/g-node.ysgsy1>.

Vahedifard, F., Adepoju, J. O., Supanich, M., Ai, H. A., Liu, X., Kocak, M., Marathu, K. K., and Byrd, S. E. (2023). “Review of deep learning and artificial intelligence models in fetal brain magnetic resonance imaging.” World Journal of Clinical Cases, 11(16), 3725–3735.

Valk, S. L., Xu, T., Margulies, D. S., Masouleh, S. K., Paquola, C., Goulas, A., Kochunov, P., Smallwood, J., Yeo, B. T. T., Bernhardt, B. C., and Eickhoff, S. B. (2020). “Shaping brain structure: Genetic and phylogenetic axes of macroscale organization of cortical thickness.” Science Advances, 6(39).

Vinall, J., Grunau, R. E., Brant, R., Chau, V., Poskitt, K. J., Synnes, A. R., and Miller, S. P. (2013). “Slower postnatal growth is associated with delayed cerebral cortical maturation in preterm newborns.” Science Translational Medicine, 5(168).

Volpe, J. J. (2018). Volpe’s neurology of the newborn.

Warrington, S., Bryant, K. L., Khrapitchev, A. A., Sallet, J., Charquero-Ballester, M., Douaud, G., Jbabdi, S., Mars, R. B., and Sotiropoulos, S. N. (2020). “Xtract - Standardised protocols for automated tractography in the human and macaque brain.” NeuroImage, 217, 116923.

Wu, J., Sun, T., Yu, B., Li, Z., Wu, Q., Wang, Y., Qian, Z., Zhang, Y., Jiang, L., and Wei, H. (2021). “Age-specific structural fetal brain atlases construction and cortical development quantification for chinese population.” NeuroImage, 241, 118412.

